## Supplementary Information for "Resilience of genetic diversity in forest trees over the Quaternary"

**This PDF file includes:**

Supplementary Materials and Methods

GenTree consortium author list

Figs. S1 to S28

Tables S2, S9 to S11, S15 to S20

**Other Supplementary Materials for this manuscript include the following:**

Tables, S1, S3 to S8 and S12 to S14

**Table_S1.xlsx** Sampling location IDs, species, coordinates (WGS84) and number of individuals in vcf v.5.3.1.

**Table_S3.xlsx.** Coordinates, expected heterozygosity at all, silent, 4-fold and 0-fold degenerate sites, and population-specific F_ST_ per population and species.

**Table_S4.xlsx.** Genetic distance and geographic distance (as pairwise F_ST_) among all pairs of populations.

**Table_S5.xlsx.** AIC, delta AIC and w_i_ computed for each species, sampling design and model.

**Table_S6.xlsx.** Confidence intervals (95%) of the parameter obtained with the panmictic models (2-epoch and 3-epoch, with one and two demographic changes, respectively) across all species and the two sampling designs (AllSamples = all samples included, OnePerPop = one haplotype per location included). The median and the 2.5 and 97.5 percentiles are provided for each parameter. NANC = population effective size of the ancestral population; NCUR = population effective size of the current population; TLATE = time in generations of the demographic event in the 2-epoch model and the most ancient demographic event in the 3-epoch model; TEARLY = time in generations of the most recent demographic event in the 3-epoch model; TLATE_year = same as TLATE but expressed in years; TEARLY_year = same as TLATE but expressed in year; Ratio NCUR/NANC = ratio of NCUR and NANC.

**Table_S7.xlsx.** Single-point estimates for several parameters obtained from simulations under seven models (SNM, 2-epoch, 3-epoch, div-e1-iso, div-e2-iso, div-e1-mig, div-e2-mig) across four sampling designs (AllSamples, OnePerPop, PairPop, SinglePop) with fastsimcoal2. These estimates were taken from the best run out of 100 independent runs (see Materials and Methods).

**Table_S8.tsv.** All results obtained with Stairway Plot 2 concatenated into a single file with columns ‘species’ and ‘population’ specific to each individual analysis. Species-wide analyses were run using two sampling designs, AllSamples and OnePerPop, and are identified as such in the ‘population’ column. For population-wide analyses the population ID is given in the ‘population’ column.

**Table_S12.xlsx.** Gene Ontology (GO) and Kyoto Encyclopedia of Genes and Genomes (KEGG) terms for selected genes.

**Table_S13.xlsx.** The set of 811 orthogroups that included at least one gene for at least six of the seven tree species.

**Table_S14.xlsx.** Other ‘orthogroups’ including genes that were found only in a reduced set of species (e.g. 59 additional genes across the three conifer species).

Supplementary Materials and Methods

Dataset descriptions

As different types of analyses require SNP sets curated and filtered based on their specific needs, we provide four versions of SNP sets as vcf files available at <https://doi.org/10.57745/DV2X0M>. Note that since several species were observed to have experienced various levels and extents of hybridization, some admixed populations were included/excluded based on the purpose of a particular analysis. For example, in the analyses of population structure, we aimed to identify the potential admixed individuals and populations. In contrast, admixed individuals and populations can have a disproportionate effect on the measures of diversity and on the site frequency spectrum and were thus excluded from the corresponding analyses.

**v.5.3** Known other species and clear hybrids removed, samples and SNPs with poor coverage or other low quality removed, organelle contigs removed (described in SNP filtering), *P. nigra* clones and cultivars removed, vcf format.

**v.5.3.1** Master dataset derived from v.5.3, without samples with incorrect taxon assignment as indicated by genetic analysis for *F. sylvatica* and *Q. petraea*, identical to v.5.3 for the other five species, vcf format.

**v.5.3.2** Derived from v.5.3.1, excludes *P. abies* populations RU_PA_19 and RU_PA_20, vcf format.

**v.6.3.1** Only includes four-fold degenerate sites, intron and intergenic sites, SNPs in high LD (1 kb windows, *r* > 0.5, PLINK v.1.90b4.9, (Gaunt et al, 2007, Chang et al. 2015) were excluded, derived from v.5.3.1

**v.6.3.2** Derived from v.6.3.1, excludes singletons, ped- and map- format.

Clone and incorrect taxon identification and filtering

We identified *Populus* x *canadensis* genotypes among *P. nigra* samples based on an excess of heterozygous positions (> 5% of calls). We confirmed these by comparing their genotype at SSR markers with the INRAE database of common *Populus* x *canadensis* cultivars (<https://urgi.versailles.inrae.fr/faidare/studies/dXJuOlVSR0kvc3R1ZHkvNjU%3D>). We detected introgression from the ornamental *P. nigra* ‘Italica’ cultivar by using this cultivar as a control in the targeted capture experiment and performing ancestry analysis using ADMIXTURE software (Alexander & Lange 2009) on the whole SNP dataset. We discarded samples with a percentage of co-ancestry with ‘Italica’ > 85%, except for one Spanish genotype (ES_PO_01_02) that is identical to ‘Italica’ and represents the genotype. We identified clones through an identity by state (IBS) calculation on the whole SNP dataset using plink software (v.1.9).

For *Q. petraea*, we removed samples with an incorrect taxon identification by comparing cluster membership in a preliminary admixture analysis (see below) with the leaf hair density and type of herbarium proofs (Rellstab et al. 2019). We identified as *Q. robur* 38 individuals forming their own cluster at K=2 and removed them for further analysis. These included seven individuals from population DE_QP_17, two from GB_QP_12, two from PL_QP_19, five from SE_QP_16, and the complete population SE_QP_15 (22 individuals).

We removed *F. sylvatica* population GR_FS_10 from the dataset because preliminary analyses of genotypic data by PCA showed it to be entirely different from the remaining populations. This stand is possibly composed of hybrids between *F. sylvatica* and *F. orientalis*.

For *P. abies*, *Picea obovata* served as an outgroup for interpreting admixture results. We excluded these samples from subsequent analyses, as well as samples from populations RU_PA_19 and RU_PA_20, which showed excessive admixture with *P. obovata*, and two *P. omorika* samples.

**GenTree consortium author list**

Audrey Albet^1^, Ricardo Alía^2^, Paraskevi Alizoti^3^, Olivier Ambrosio^4^, Filippos A. Aravanopoulos^2^, Francisco Auñón^2^, Camilla Avanzi^5^, Evangelia Avramidou^3^, Marko Bajc^6^, Eduardo Ballesteros^2^, Evangelos Barbas^3^, Cristina C. Bastias^7^, Catherine Bastien^8^, Giorgia Beffa^9,10^, Raquel Benavides^11^, Vanina Benoit^12^, Frédéric Bernier^1^, Henri Bignalet^1^, Guillaume Bodineau^13^, Damien Bouic^1^, Sabine Brodbeck^9^, William Brunetto^4^, Jurata Buchovska^14^, Corinne Buret^12^, Melanie Buy^15^, Ana M. Cabanillas-Saldaña^16^, Bárbara Carvalho^11^, Nicolas Cheval^1^, José M. Climent^2^, Marianne Correard^17^, Eva Cremer^18^, Darius Danusevičius^14^, Nicolas De Girardi^9^, Fernando Del Caño^2^, Jean-Luc Denou^1^, Bernard Dokhelar^1^, Remi Dourthe^1^, Alexis Ducousso^20^, Anna-Maria Farsakoglou^3^, Patrick Fonti^9^, Ioannis Ganopoulos^21^, José M. García del Barrio^2^, Olivier Gilg^17^, René Graf^9^, Alan Gray^22^, Christoph Hartleitner^23^, Katrin Heer^24^, Enja Hollenbach^25^, Agathe Hurel^19^, Bernard Issenhuth^1^, Florence Jean^4^, Arnaud Jouineau^3^, Jan-Philipp Kappner^25^, Katri Kärkkäinen^26^, Robert Kesälahti^27^, Florian Knutzen^18^, Sonja T. Kujala^26^, Timo A. Kumpula^27^, Mariaceleste Labriola^28^, Celine Lalanne^19^, Johannes Lambertz^25^, Gregoire Le-Provost^19^, Vincent Lejeune^13^, Joseph Levillain^29^, Mirko Liesebach^30^, David López-Quiroga^11^, Ermioni Malliarou^3^, Jérémy Marchon^9^, Nicolas Mariotte^4^, Elisabet Martínez-Sancho^9^, Antonio Mas^11^, Silvia Matesanz^31^, Benjamin Meier^9^, Helge Meischner^25^, Célia Michotey^15^, Sandro Morganti^9^, Tor Myking^20^, Daniel Nievergelt^9^, Anne Eskild Nilsen^20^, Eduardo Notivol^32^, Geir Ostreng^21^, Birte Pakull^31^, Annika Perry^23^, Andrea Piotti^6^, Nicolas Poinot^5^, Mehdi Pringarbe^18^, Luc Puzos^5^, Annie Raffin^5^, José A. Ramírez-Valiente^3^, Oliver Reutimann^9,33^, Sebastian Richter^25^, Juan J. Robledo-Arnuncio^2^, Sergio San Segundo^2^, Outi Savolainen^27^, Volker Schneck^34^, Silvio Schueler^35^, Vladimir Semerikov^36^, Lenka Slámová^9,37^, Jørn Henrik Sønstebø^20^, Ilaria Spanu^5^, Jean Thevenet^17^, Mari Mette Tollefsrud^20^, Norbert Turion^17^, Fernando Valladares^11^, Marc Villar^12^, Georg von Arx^9^, Johan Westin^38^

Affiliations:

^1^INRAE, UEFP, 33610 Cestas, France

^2^Institute of Forest Sciences (ICIFOR-INIA), CSIC, 28040 Madrid, Spain

^3^Aristotle University of Thessaloniki, School of Forestry and Natural Environment, Laboratory of Forest Genetics and Tree Improvement, 54124 Thessaloniki, Greece

^4^INRAE, URFM, 84914 Avignon, France

^5^Institute of Biosciences and Bioresources, National Reasearch Council of Italy (IBBR-CNR), 50019 Sesto Fiorentino, Italy

^6^Slovenian Forestry Institute, 1000 Ljubljana, Slovenia

^7^CEFE, CNRS, UMR 5175, 34090 Montpellier, France

^8^INRAE, ECODIV, 45075 Orléans, France

^9^Swiss Federal Research Institute WSL, 8903 Birmensdorf, Switzerland

^10^Institute of Plant Sciences, University of Bern, 3013 Bern, Switzerland

^11^Department of Biogeography and Global Change, Museo Nacional de Ciencias Naturales, CSIC, 28006 Madrid, Spain

^12^INRAE, ONF, BioForA, 45075 Orléans, France

^13^INRAE, GBFOR, 45075 Orléans, France

^14^Vytautas Magnus University, 53361 Akademija, Lithuania

^15^INRAE, URGI, 78026 Versailles, France

^16^Departamento de Agricultura, Ganadería y Medio Ambiente, Gobierno de Aragón, 50071 Zaragoza, Spain

^17^INRAE, UEFM, 84914 Avignon, France

^18^Bavarian Institute for Forest Genetics, 83317 Teisendorf, Germany

^19^University of Bordeaux, INRAE, BIOGECO, 33770 Cestas, France

^20^Division of Forestry and Forest Resources, Norwegian Institute of Bioeconomy Research (NIBIO), 1431 Ås, Norway

^21^Institute of Plant Breeding and Genetic Resources, Hellenic Agricultural Organization DEMETER, 57001 Thermi, Greece

^22^UK Centre for Ecology & Hydrology (UKCEH), EH26 0QB Bush Estate, United Kingdom

^23^LIECO GmbH & Co KG, 8775 Kalwang, Austria

^24^Forest Genetics, Albert-Ludwigs Universität Freiburg, 79098 Freiburg, Germany

^25^Faculty of Biology, Plant Ecology and Geobotany, Philipps University Marburg, 35043 Marburg, Germany

^26^Natural Resources Institute Finland, University of Oulu, 90014 Oulu, Finland

^27^University of Oulu, 90014 Oulu, Finland

^28^Institute of Biosciences and Bioresources, National Reasearch Council of Italy (IBBR-CNR), 50019 Sesto Fiorentino, Italy

^29^Université de Lorraine, AgroParisTech, INRAE, SILVA, 54000 Nancy, France

^30^Thünen Institute of Forest Genetics, 22927 Grosshansdorf, Germany

^31^Área de Biodiversidad y Conservación, Universidad Rey Juan Carlos, 28933 Móstoles, Spain

^32^Centro de Investigación y Tecnología Agroalimentaria de Aragón, Dpto. de Sistemas Agrarios, Forestales y Medio Ambiente (CITA), 50059 Zaragoza, Spain

^33^Department of Environmental Systems Sciences, ETH Zurich, 8092 Zurich, Switzerland

^34^Thünen Institute of Forest Genetics, 15377 Waldsieversdorf, Germany

^35^Austrian Research Centre for Forests (BFW), 1131 Wien, Austria

^36^Institute of Plant and Animal Ecology, Ural branch of RAS, 620144 Ekaterinburg, Russia

^37^Institute for Environmental Sciences, University of Geneva, 1205 Geneva, Switzerland

^38^Skogforsk, 91821 Sävar, Sweden

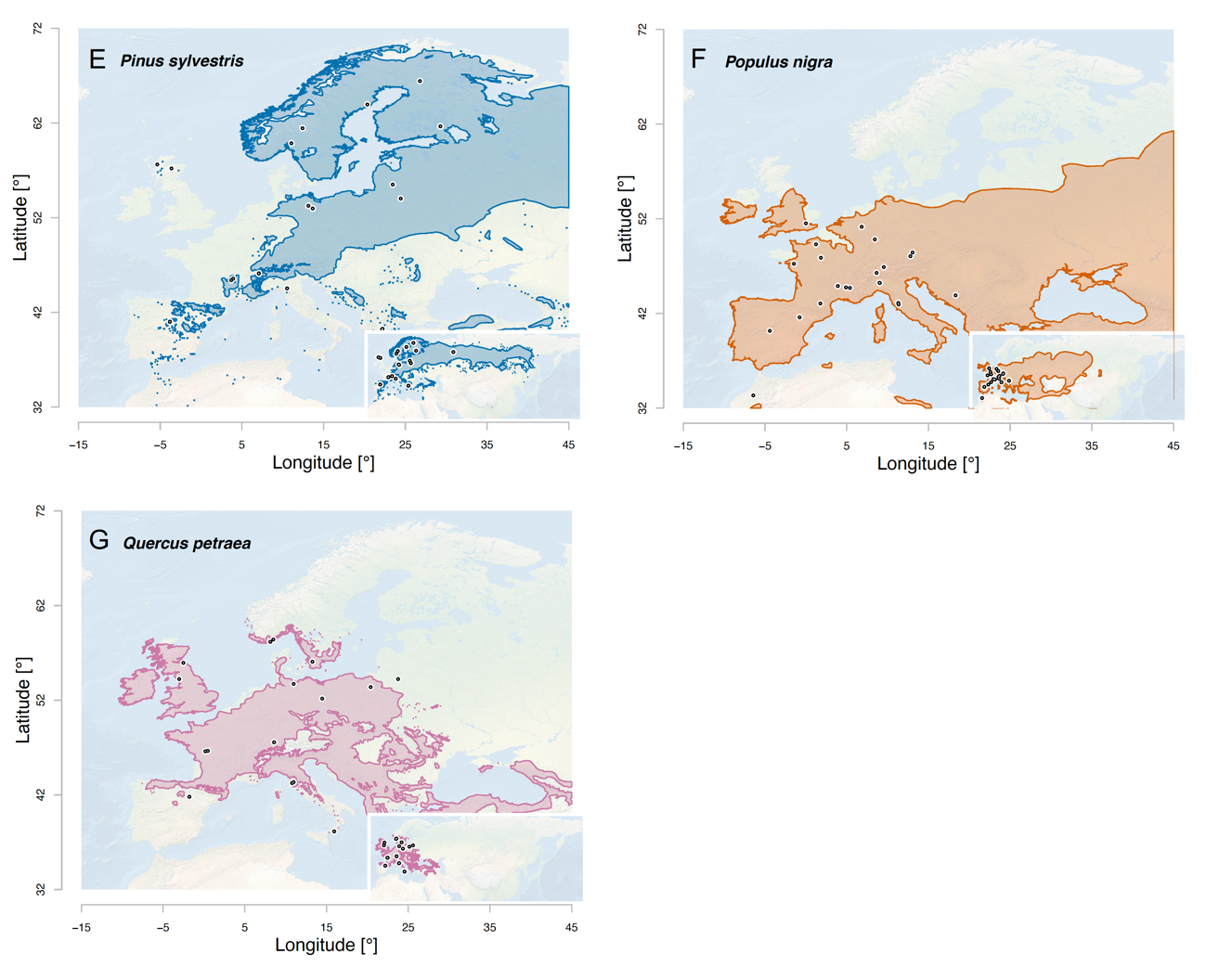

Figure S1. Sampling design. Species range and sampling locations (black dots) for the seven species analyzed in this study. Species ranges are based on the EUFORGEN database ([www.euforgen.org](http://www.euforgen.org)).

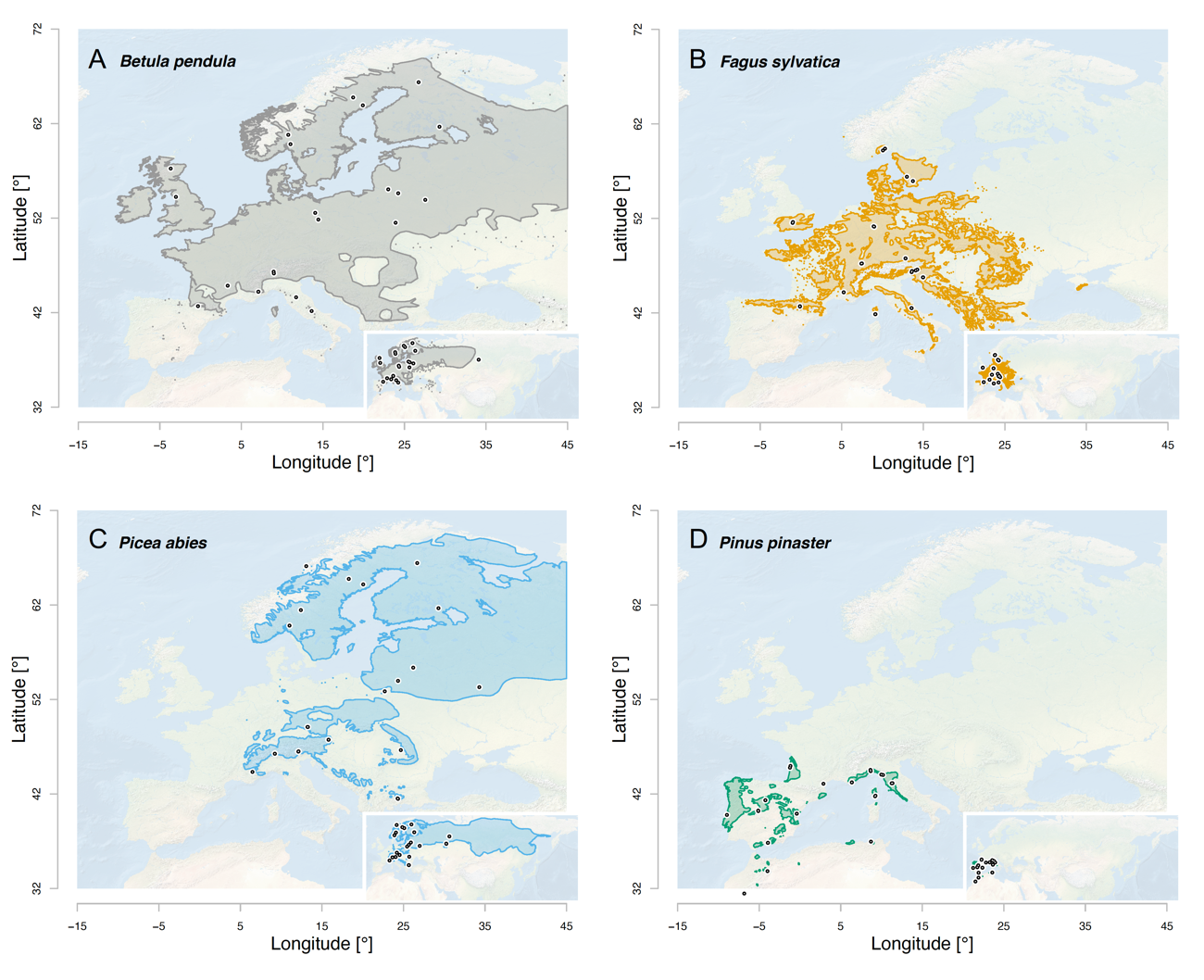

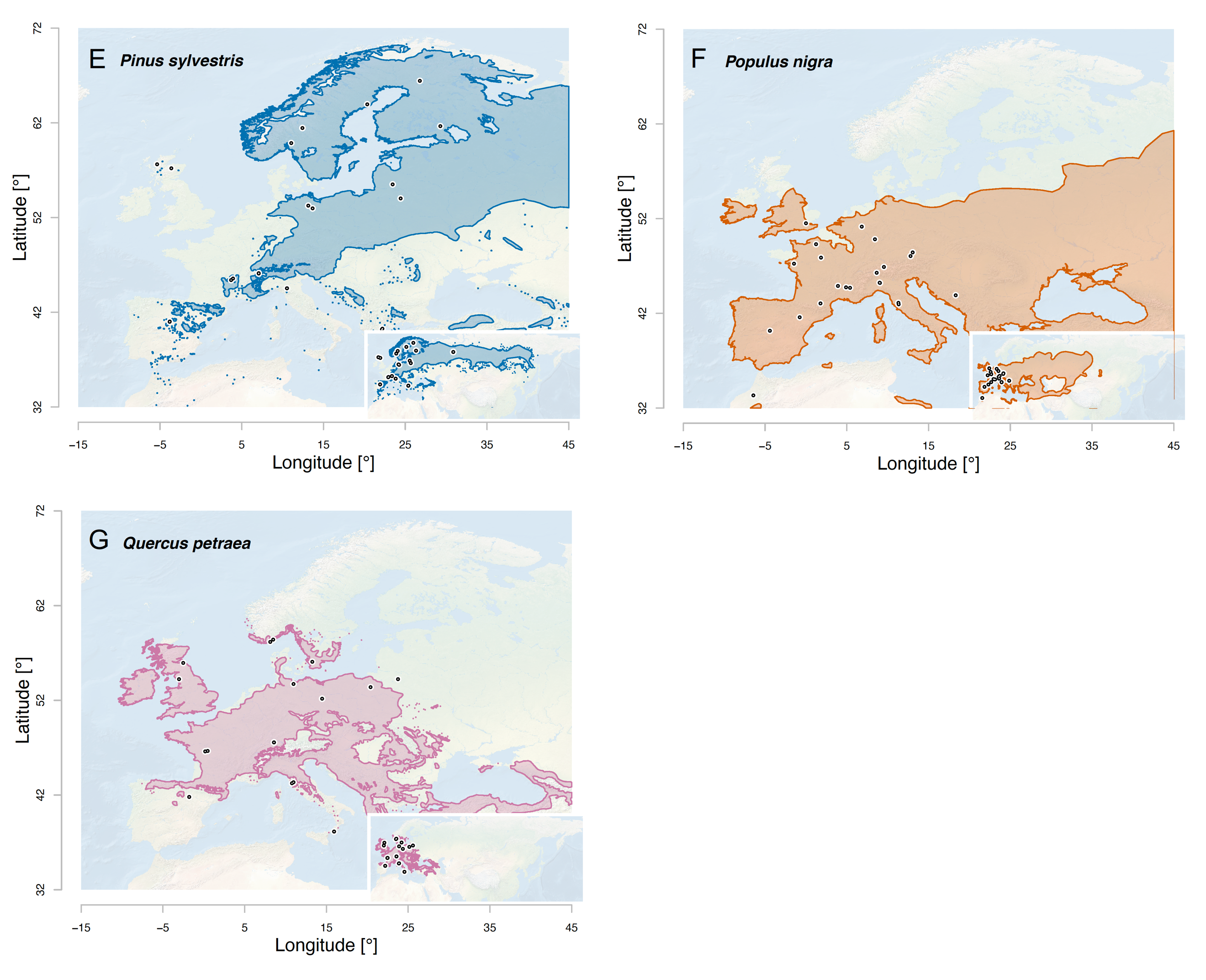

Figure S1. Sampling design. Species range and sampling locations (black dots) for the seven species analyzed in this study. Species ranges are based on the EUFORGEN database ([www.euforgen.org](http://www.euforgen.org)).

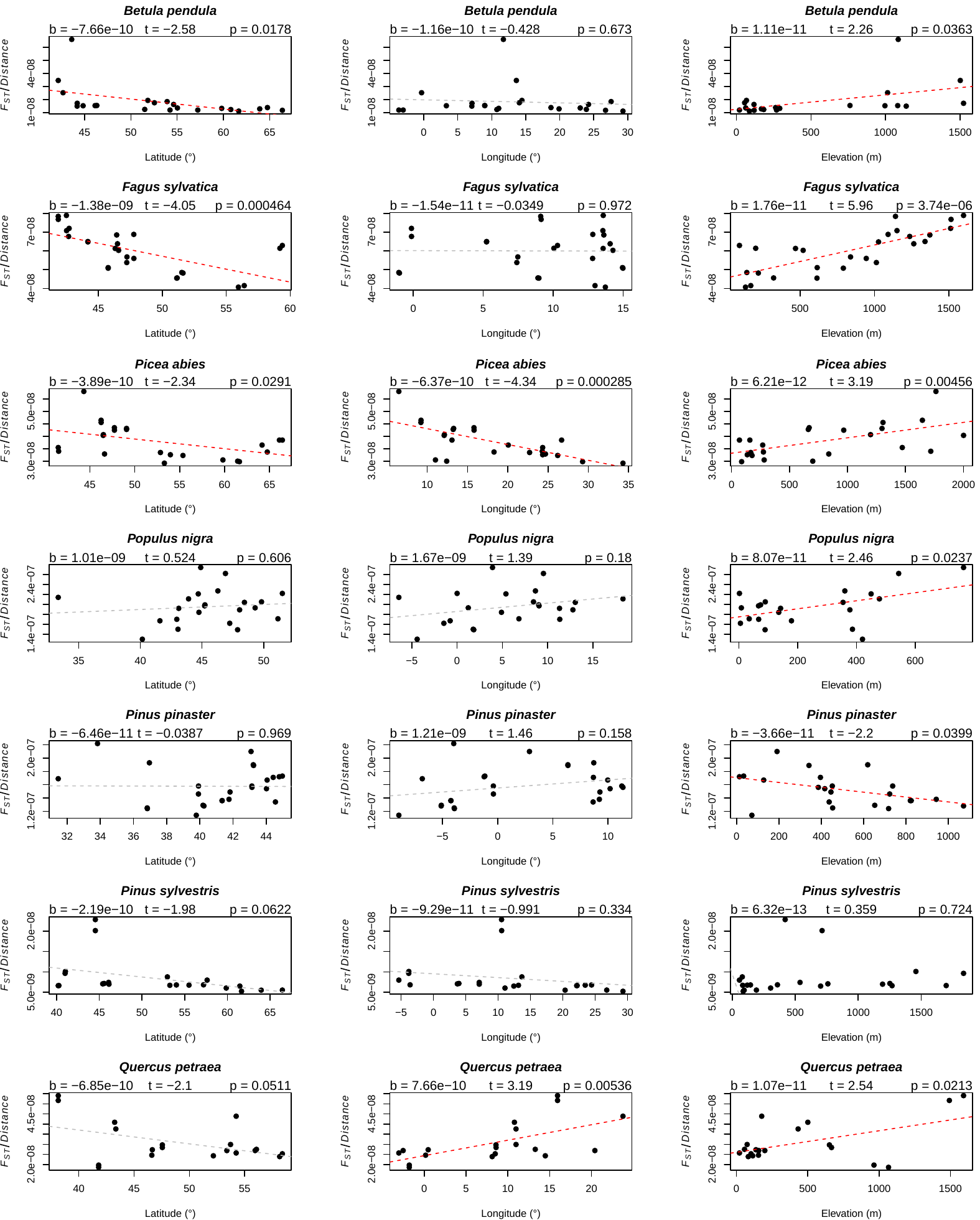

**Figure S2. Variation in population-specific *F*_ST_ across space.** Population-specific *F_ST_* (see section ‘scaling and computing population-specific *F****_ST_***’) were regressed over latitude, longitude or elevation. The value of the slope of the linear regressions (b) and the *t*- and *P*-values are presented above the plot. Where *P* < 0.05 the regression line is shown in red.

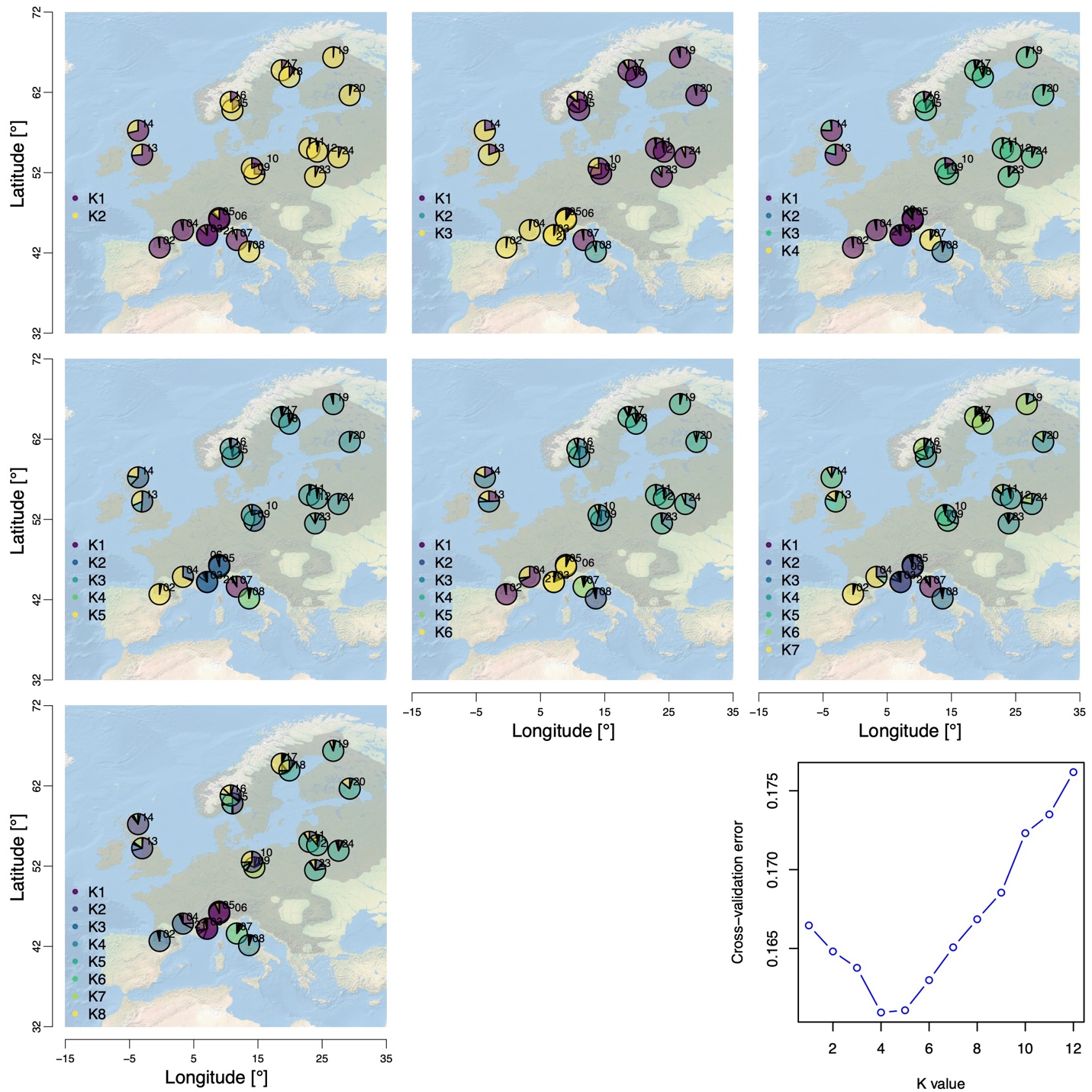

Figure S3. Admixture analysis among populations of *Betula pendula* based on the targeted capture SNP dataset. Geographic distribution of the level of admixture for each population, with colors in pie charts reflecting average assignment probabilities to the respective genetic group (*K* = 2–8). Population codes are explained in Table S1. The variation in cross-validation error across *K* values is represented in the bottom right panel, with the lowest value being the optimal number of genetic groups.

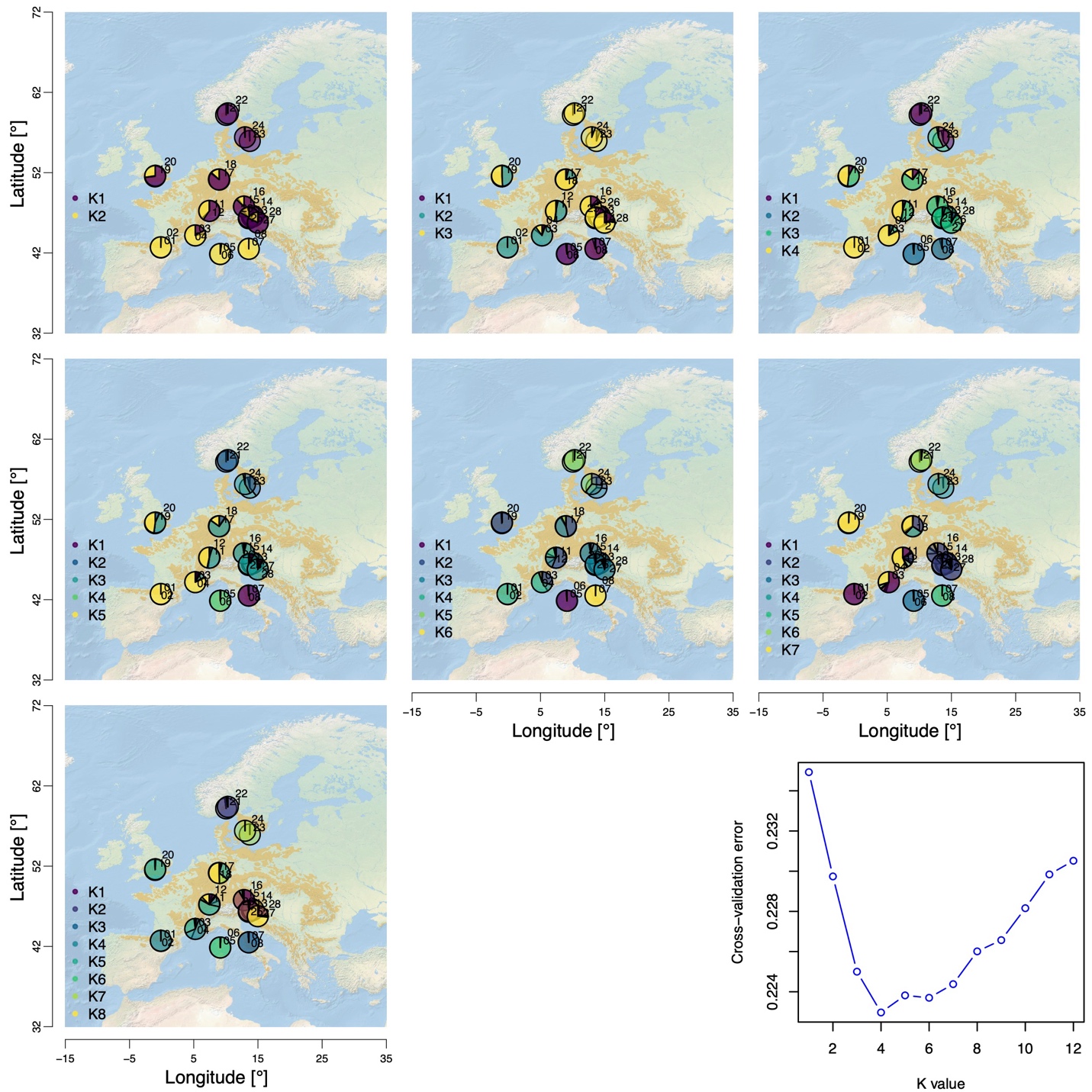

Figure S4. Admixture analysis among populations of *Fagus sylvatica* based on the targeted capture SNP dataset. Geographic distribution of the level of admixture for each population, with colors in pie charts reflecting average assignment probabilities to the respective genetic group (*K* = 2–8). Population codes are explained in Table S1. The variation in cross-validation error across *K* values is represented in the bottom right panel, with the lowest value being the optimal number of genetic groups.

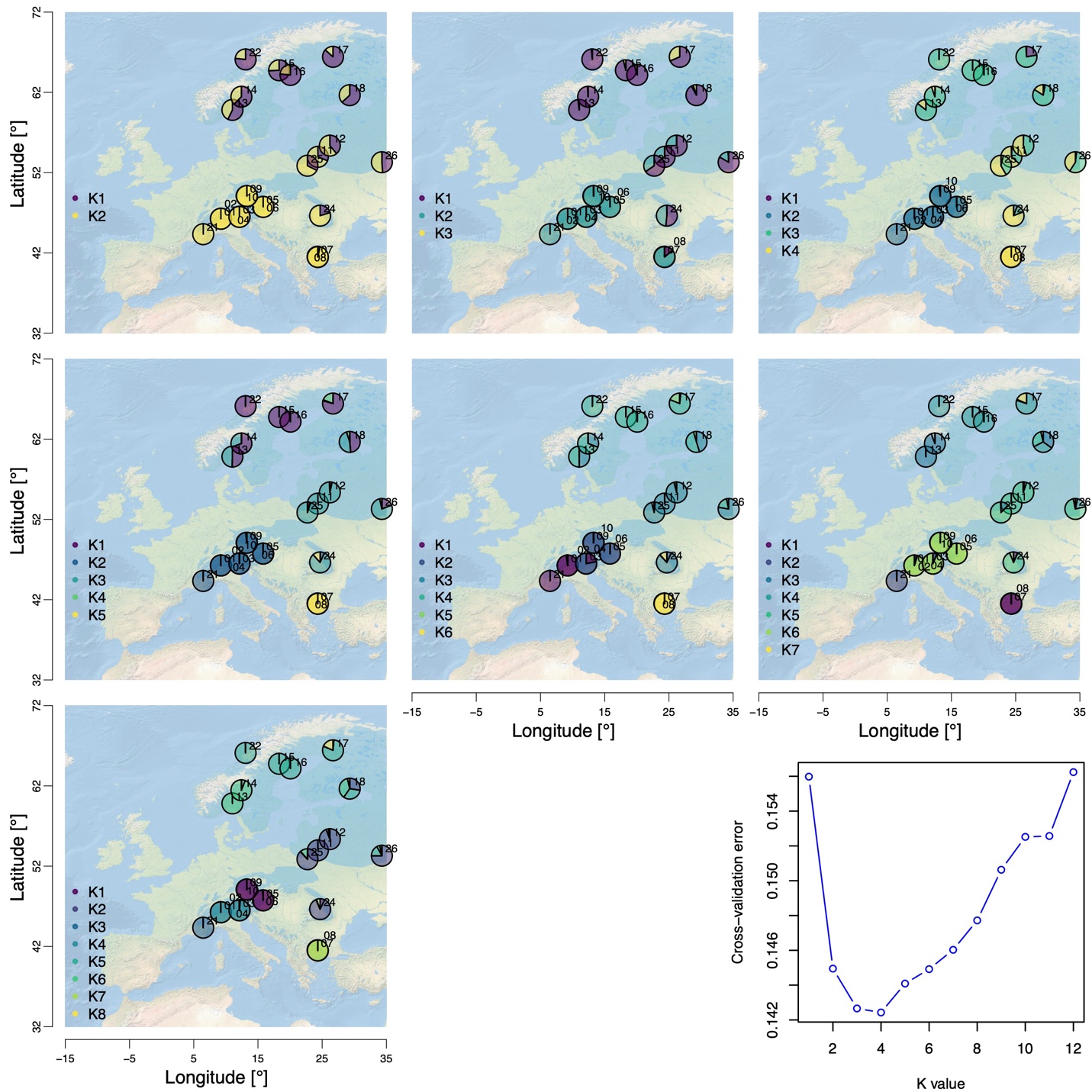

**Figure S5**. Admixture analysis among populations of *Picea abies* based on the targeted capture SNP dataset. Geographic distribution of the level of admixture for each population, with colors in pie charts reflecting average assignment probabilities to the respective genetic group (*K* = 2–8). Population codes are explained in Table S1. The variation in cross-validation error across *K* values is represented in the bottom right panel, with the lowest value being the optimal number of genetic groups.

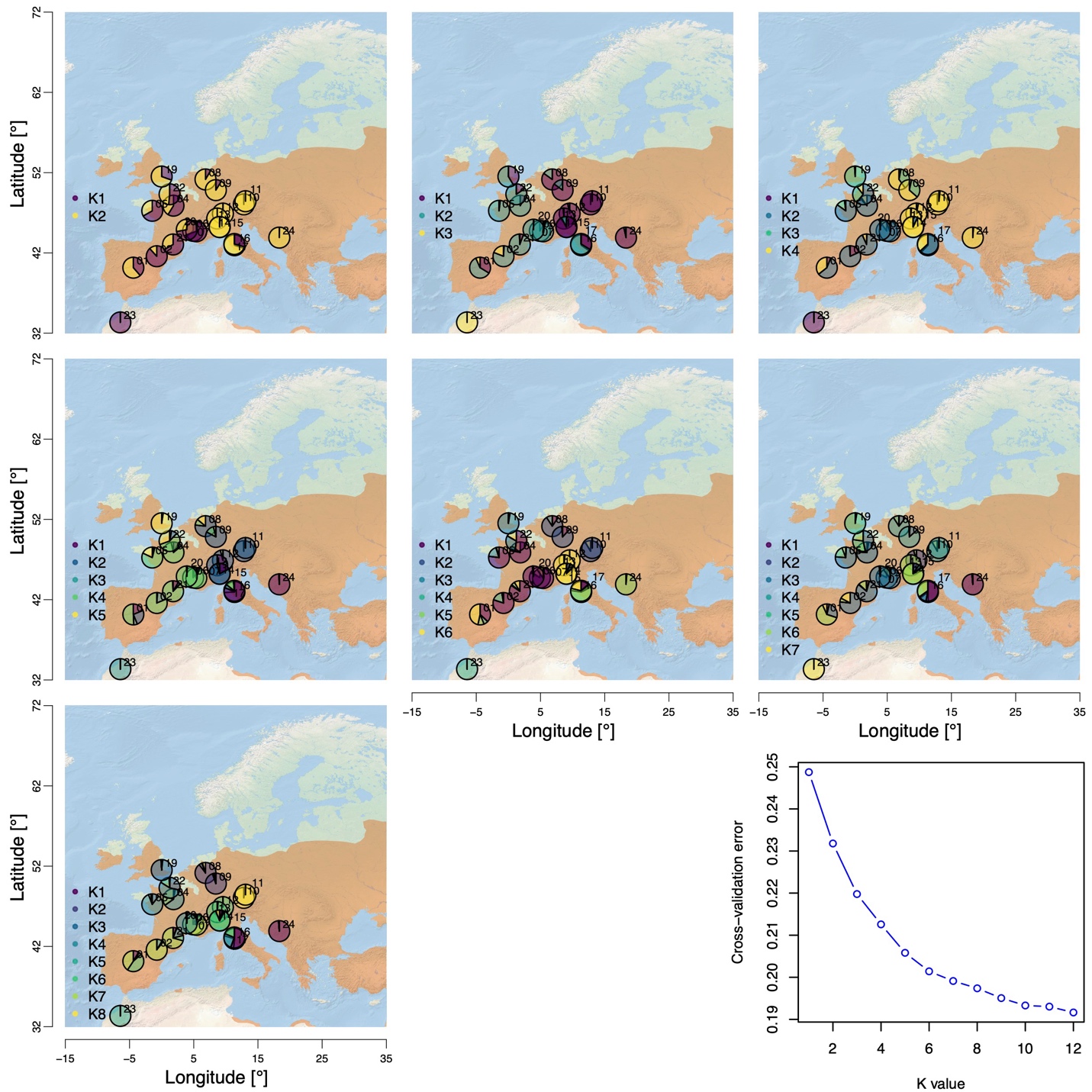

**Figure S6.** Admixture analysis among populations of *Populus nigra* based on the targeted capture SNP dataset. Geographic distribution of the level of admixture for each population, with colors in pie charts reflecting average assignment probabilities to the respective genetic group (*K* = 2–8). Population codes are explained in Table S1. The variation in cross-validation error across *K* values is represented in the bottom right panel, with the lowest value being the optimal number of genetic groups.

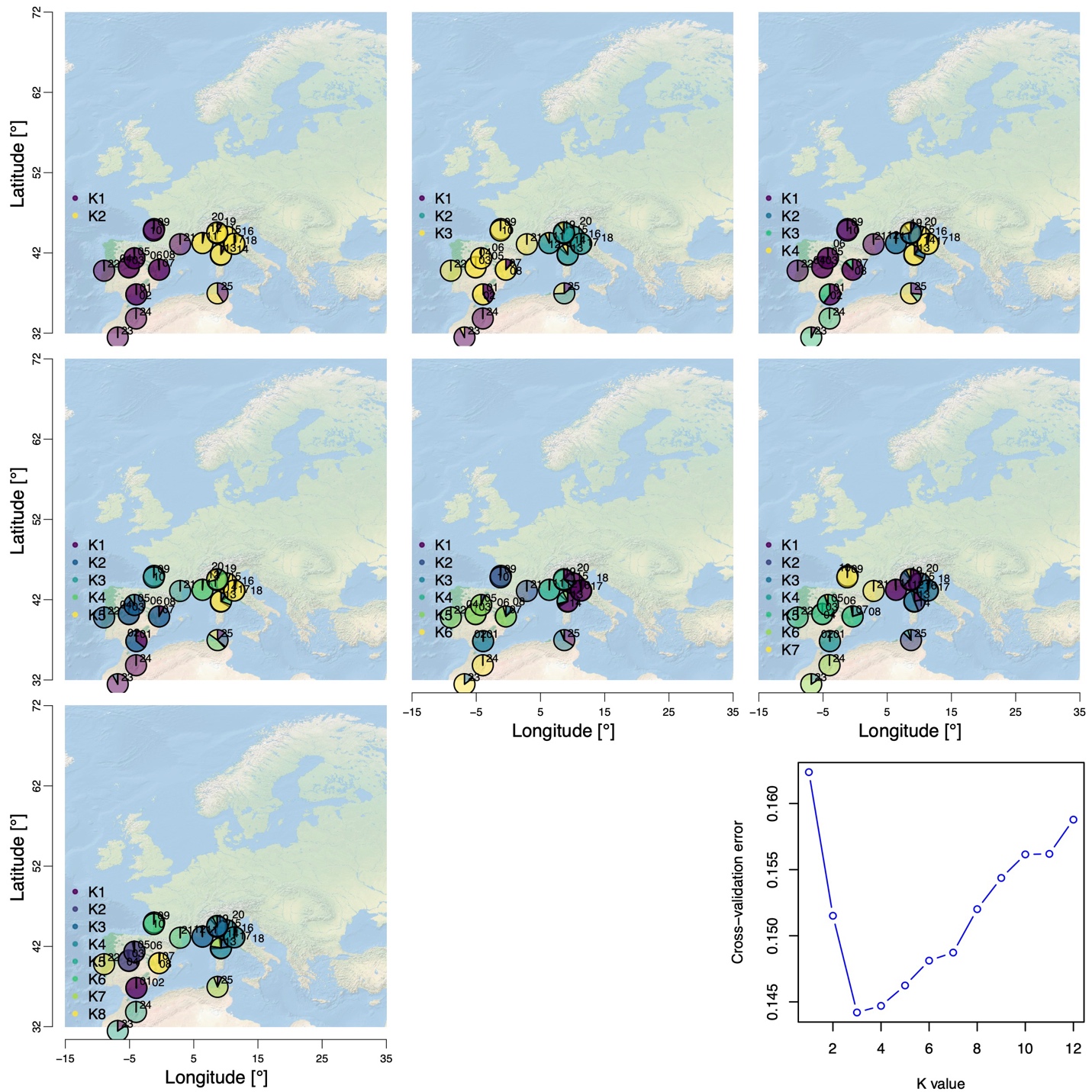

**Figure S7.** Admixture analysis among populations of *Pinus pinaster* based on the targeted capture SNP dataset. Geographic distribution of the level of admixture for each population, with colors in pie charts reflecting average assignment probabilities to the respective genetic group (*K* = 2–8). Population codes are explained in Table S1. The variation in cross-validation error across *K* values is represented in the bottom right panel, with the lowest value being the optimal number of genetic groups.

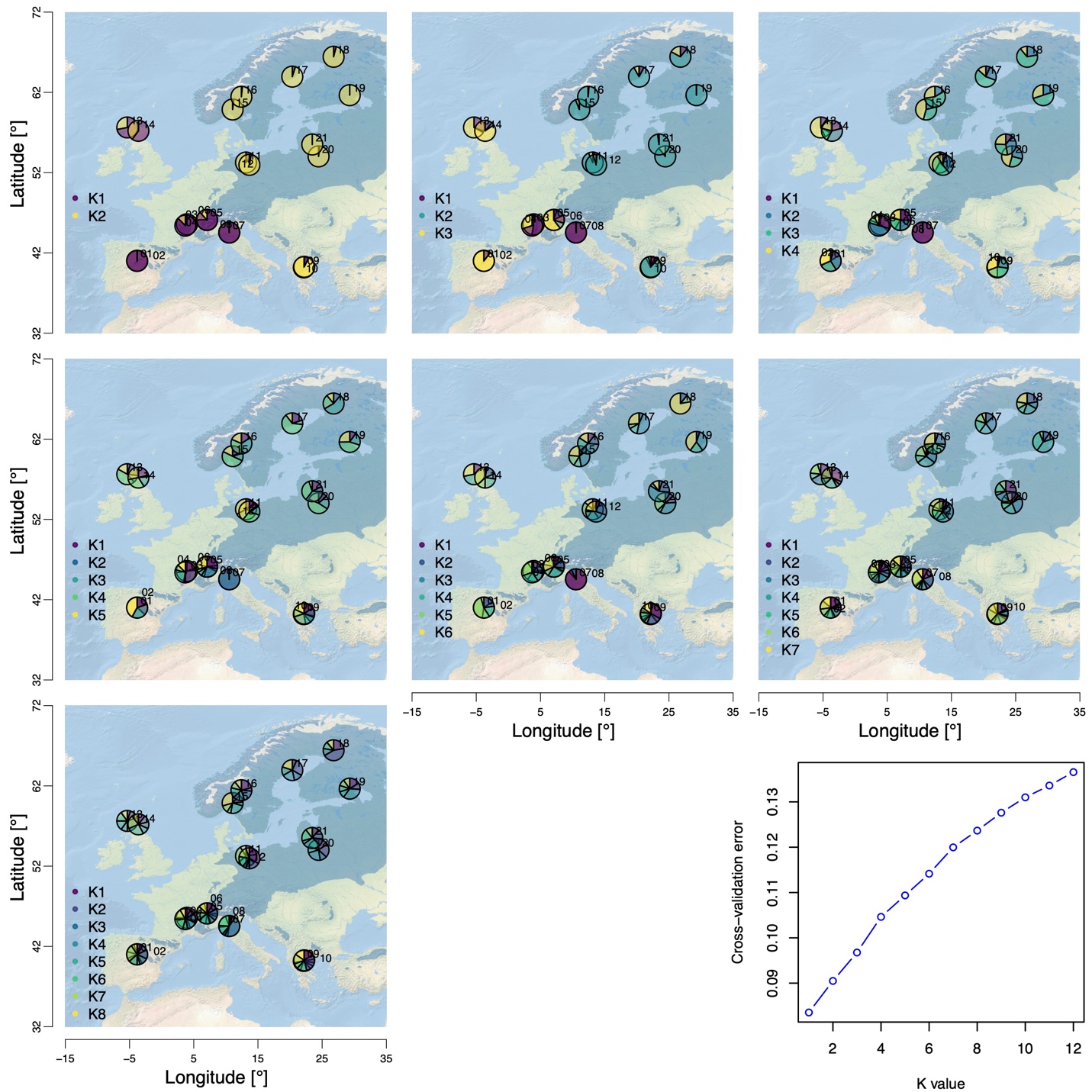

**Figure S8.** Admixture analysis among populations of *Pinus sylvestris* based on the targeted capture SNP dataset. Geographic distribution of the level of admixture for each population, with colors in pie charts reflecting average assignment probabilities to the respective genetic group (*K* = 2–8). Population codes are explained in Table S1. The variation in cross-validation error across *K* values is represented in the bottom right panel, with the lowest value being the optimal number of genetic groups.

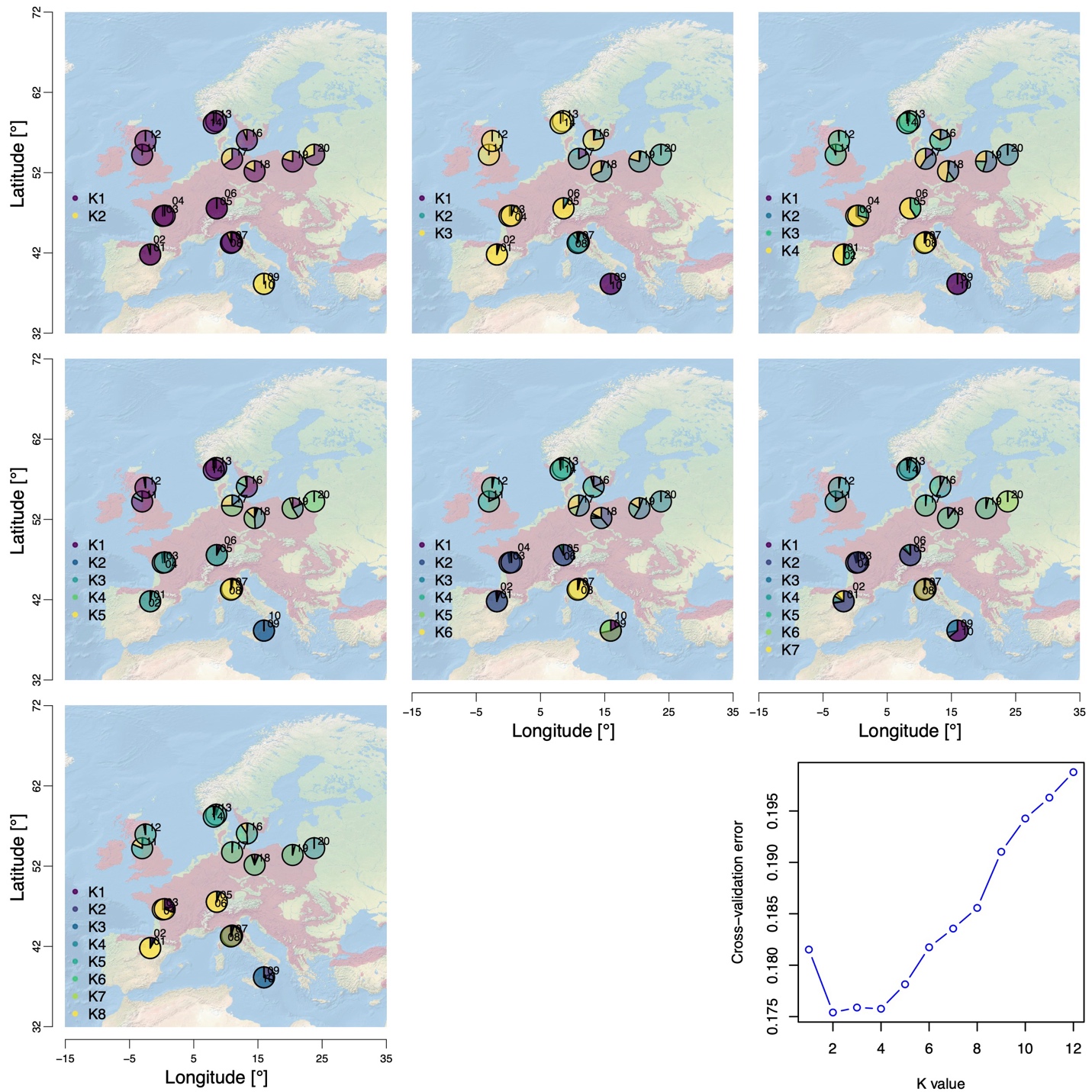

**Figure S9.** Admixture analysis among populations of *Quercus petraea* based on the targeted capture SNP dataset. Geographic distribution of the level of admixture for each population, with colors in pie charts reflecting average assignment probabilities to the respective genetic group (*K* = 2–8). Population codes are explained in Table S1. The variation in cross-validation error across *K* values is represented in the bottom right panel, with the lowest value being the optimal number of genetic groups.

Figure S10. Level of admixture within and among individuals of *Betula pendula* based on the targeted capture SNP dataset. Colors reflect assignment probabilities (i.e. *Q* scores) to respective genetic group (*K* = 2–8). Population codes are explained in Table S1. Singletons were excluded in the admixture analysis.

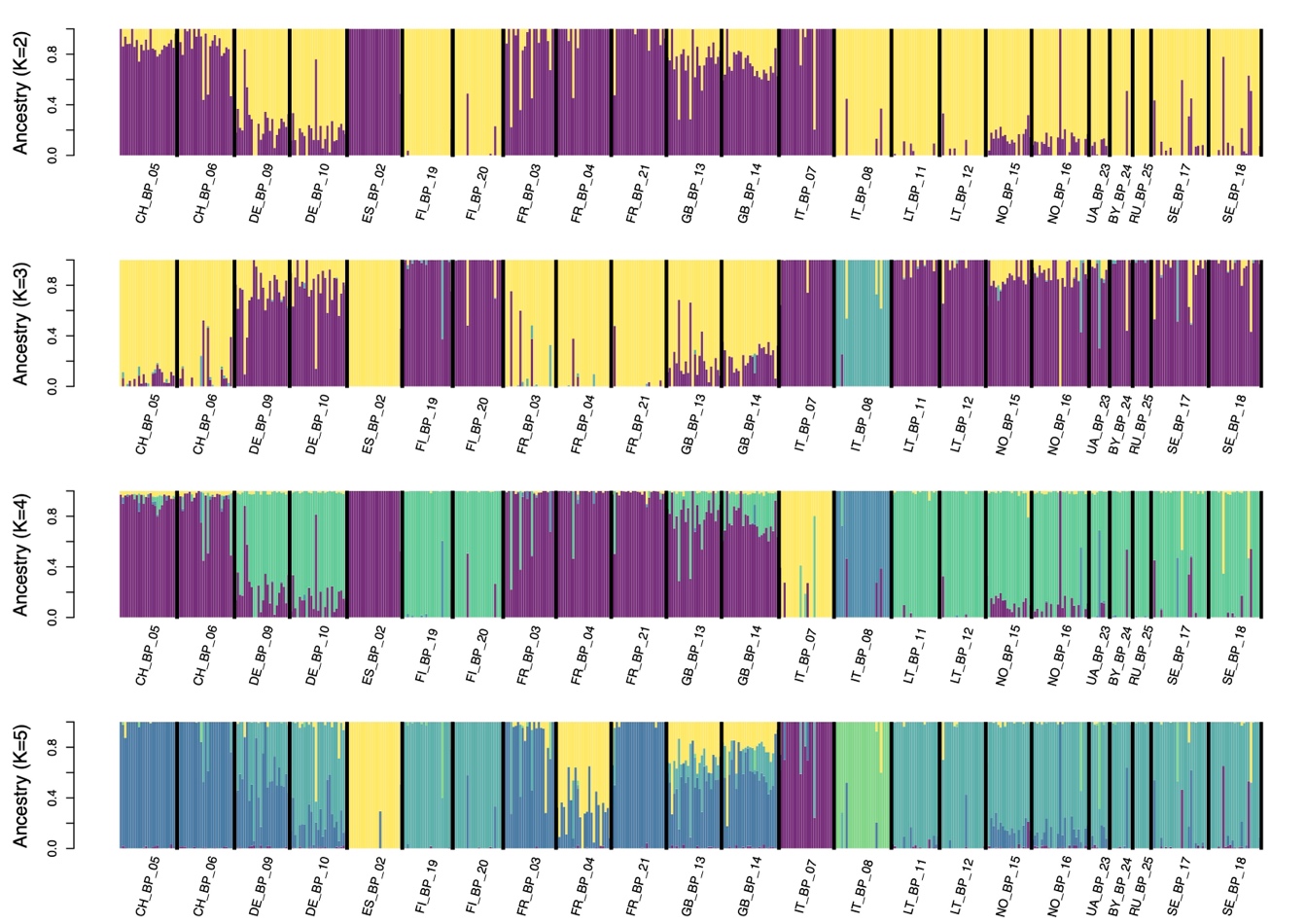

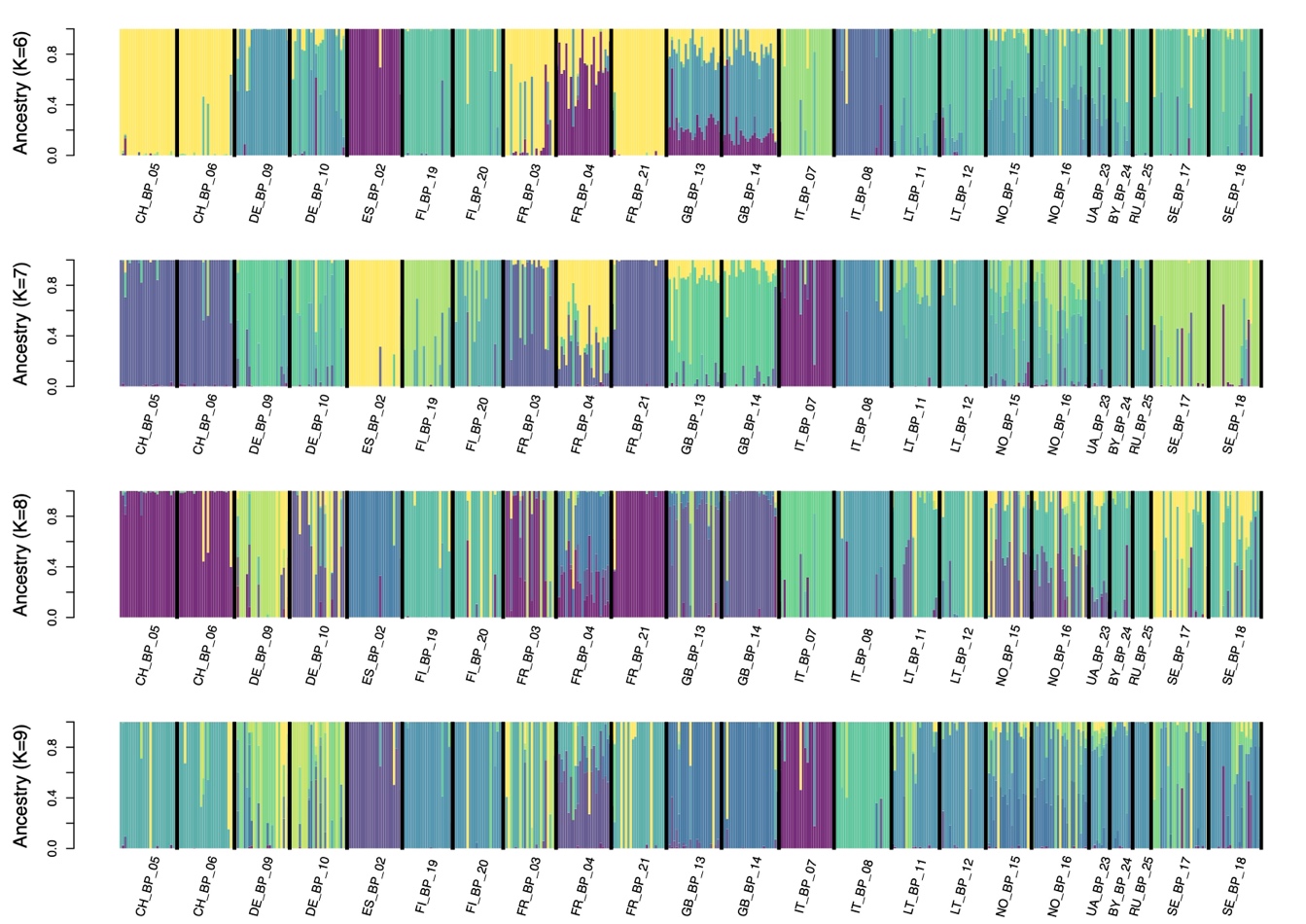

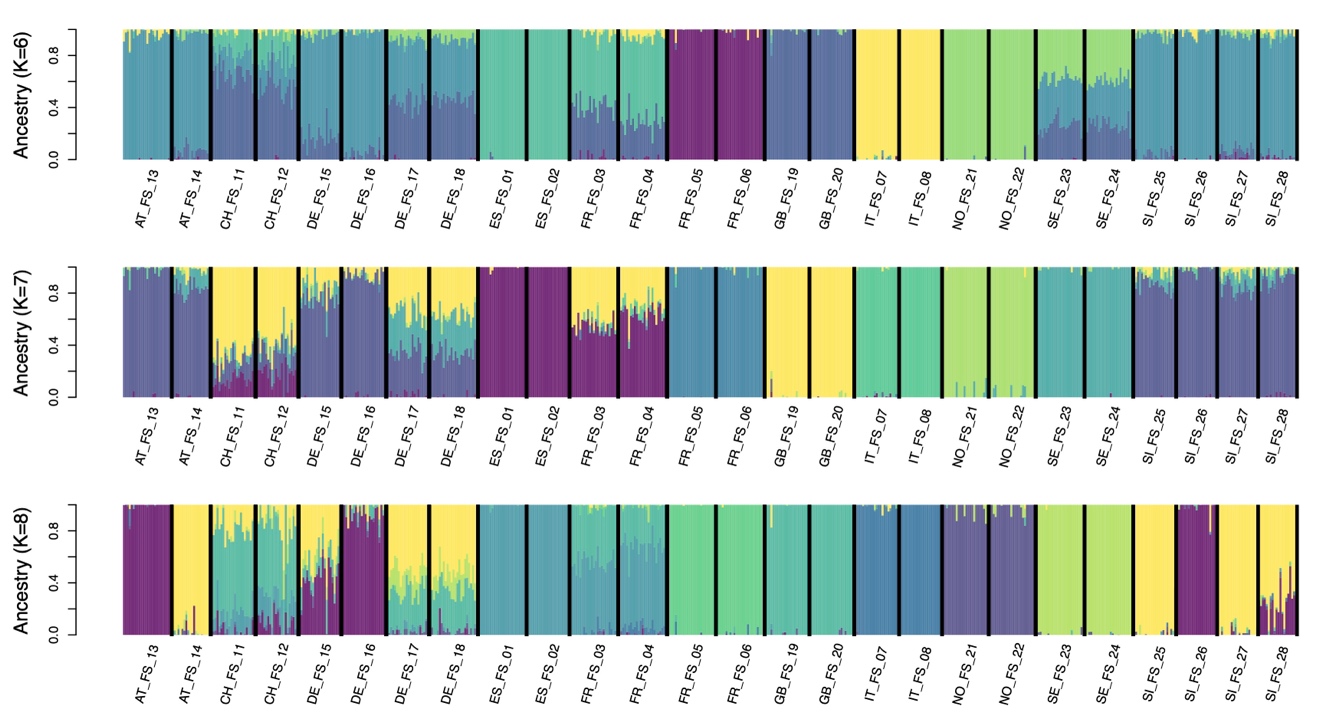

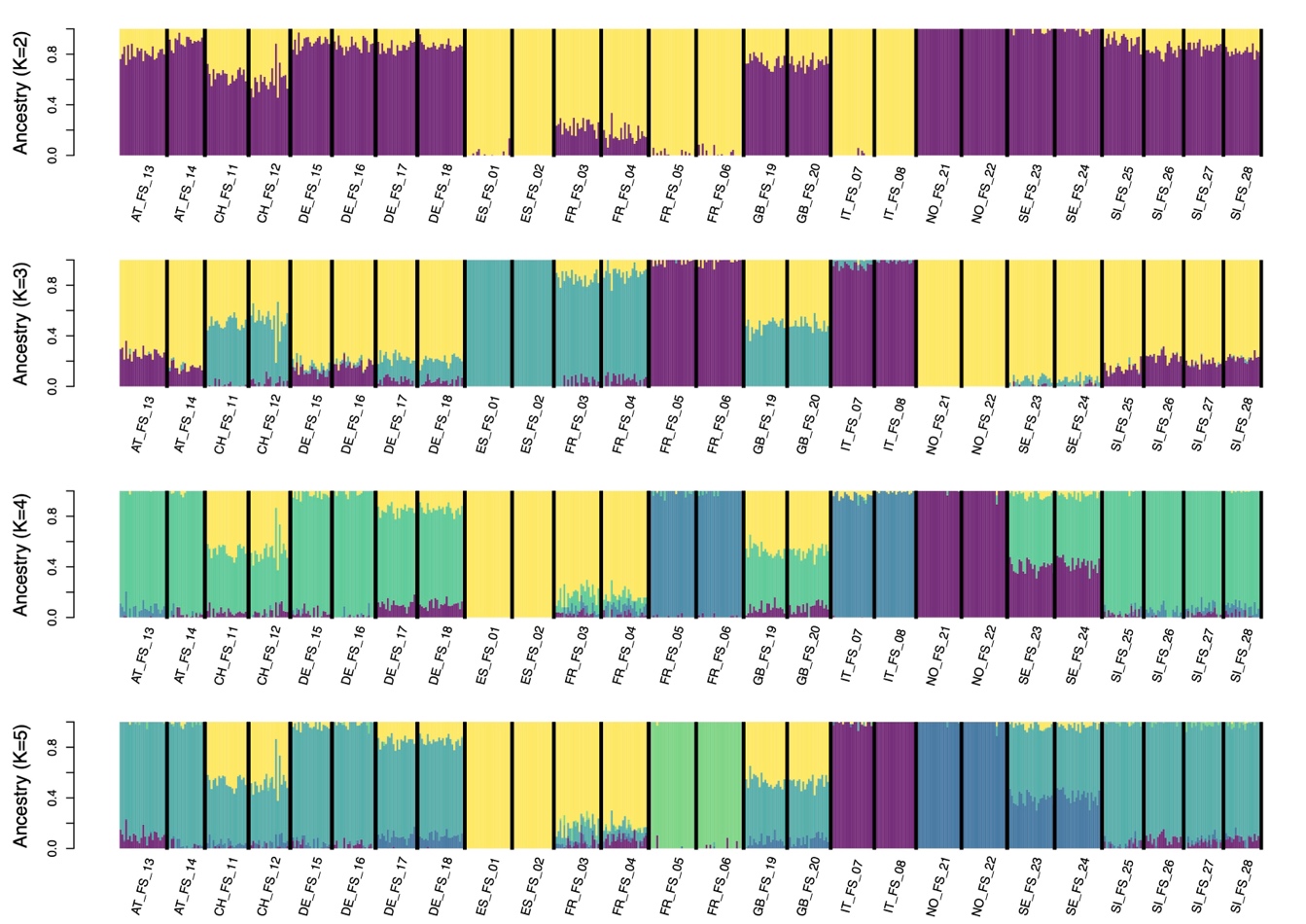

Figure S11. Level of admixture within and among individuals of *Fagus sylvatica* based on the targeted capture SNP dataset. Colors reflect assignment probabilities (i.e. *Q* scores) to respective genetic group (*K* = 2–8). Population codes are explained in Table S1. Singletons were excluded in the admixture analysis.

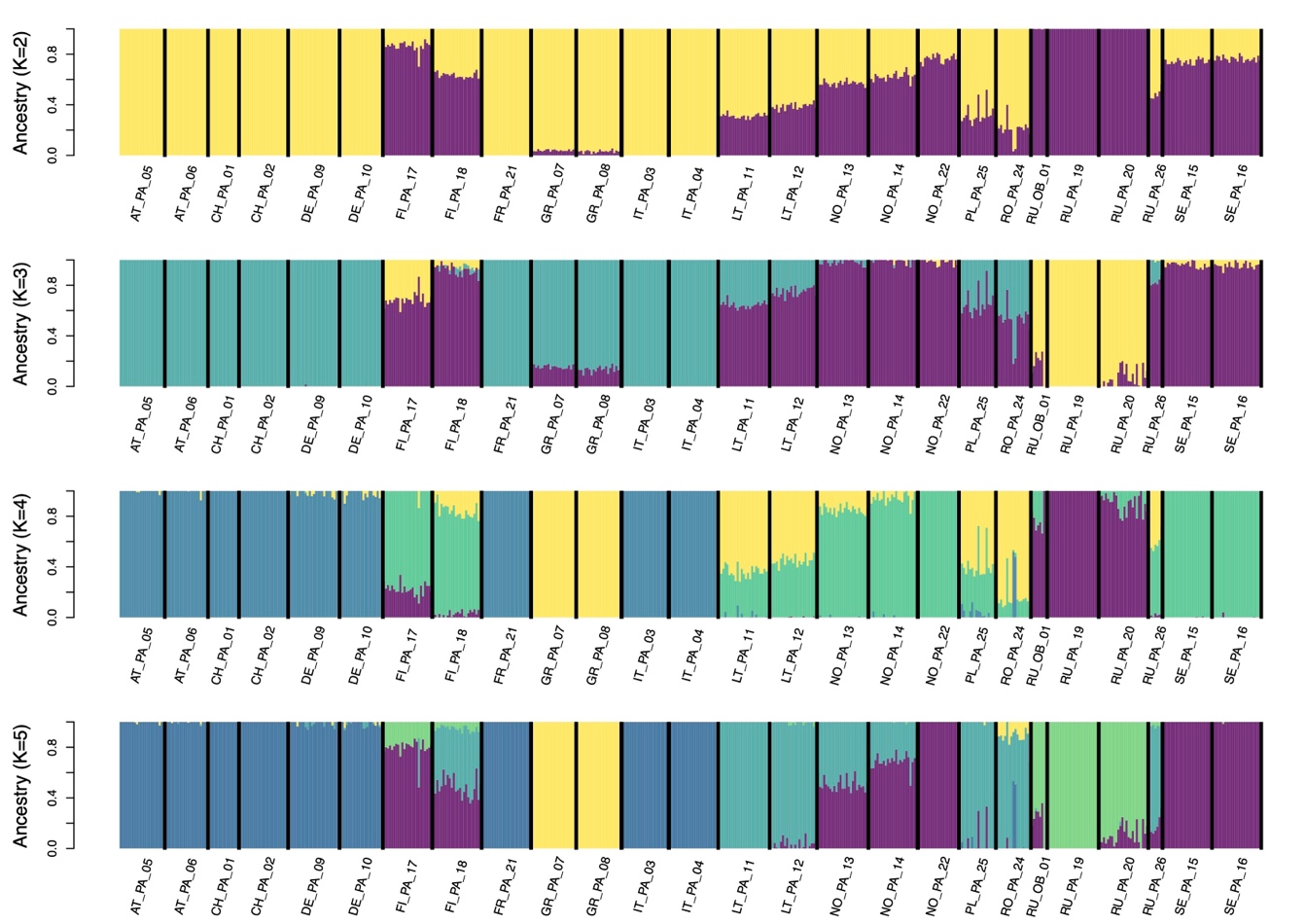

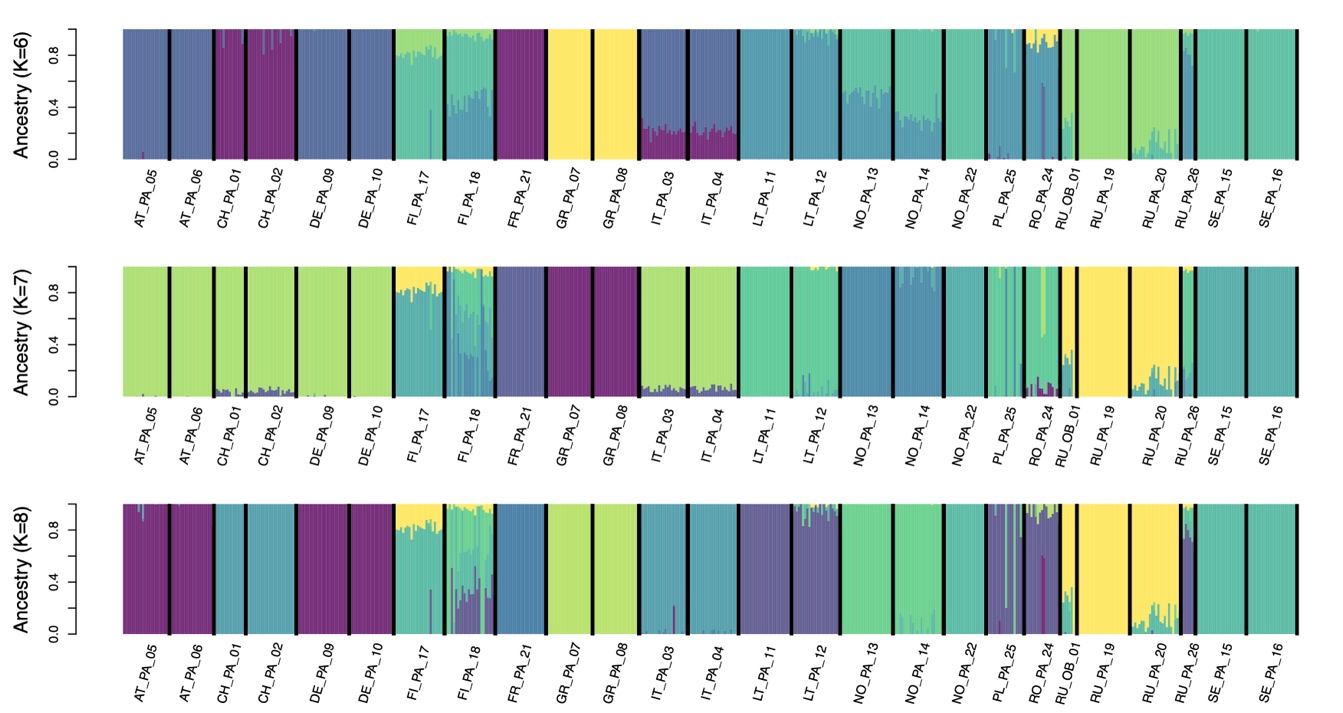

Figure S12. Level of admixture within and among individuals of *Picea abies* based on the targeted capture SNP dataset. Colors reflect assignment probabilities (i.e. *Q* scores) to respective genetic group (*K* = 2–8). Population codes are explained in Table S1.

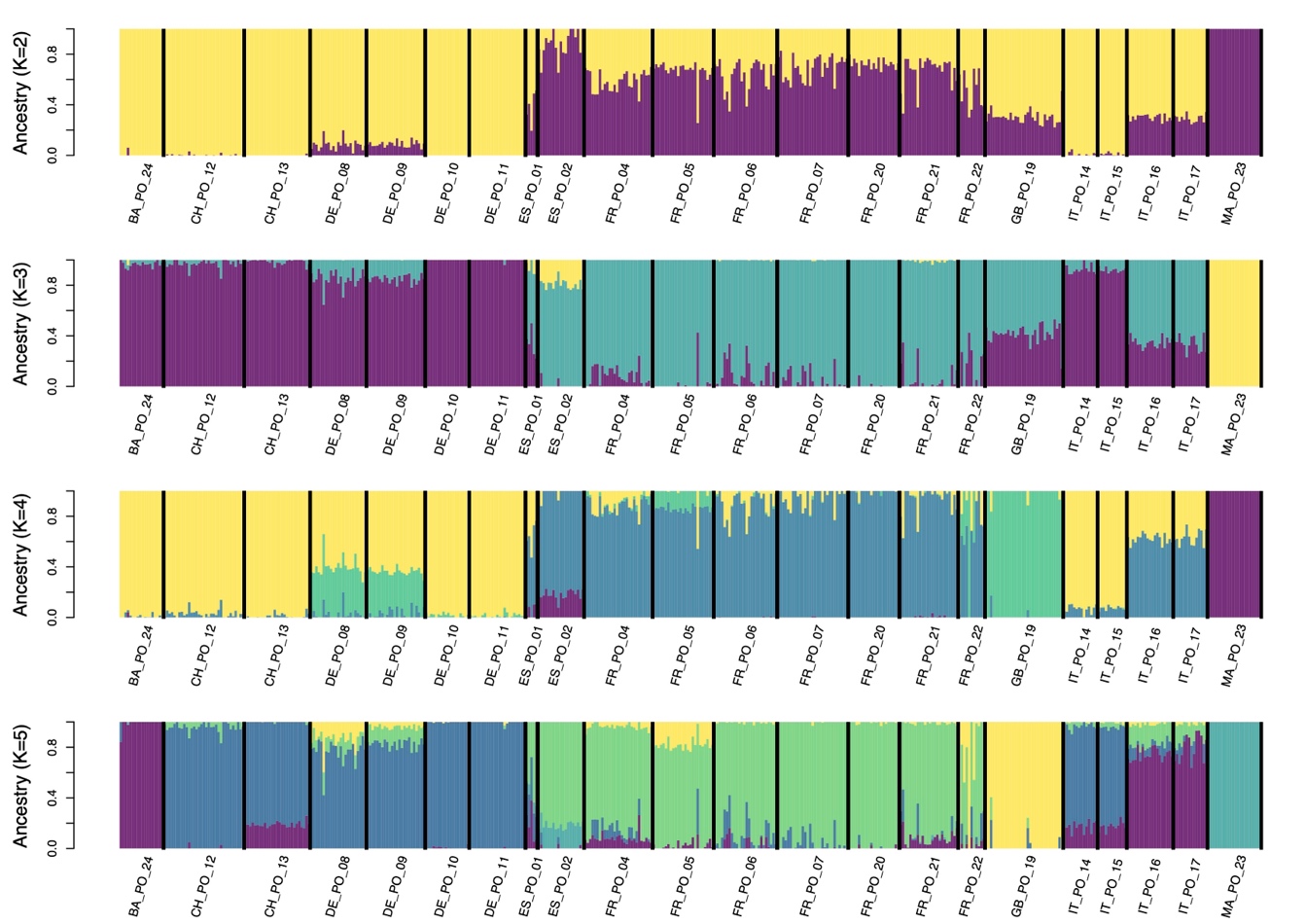

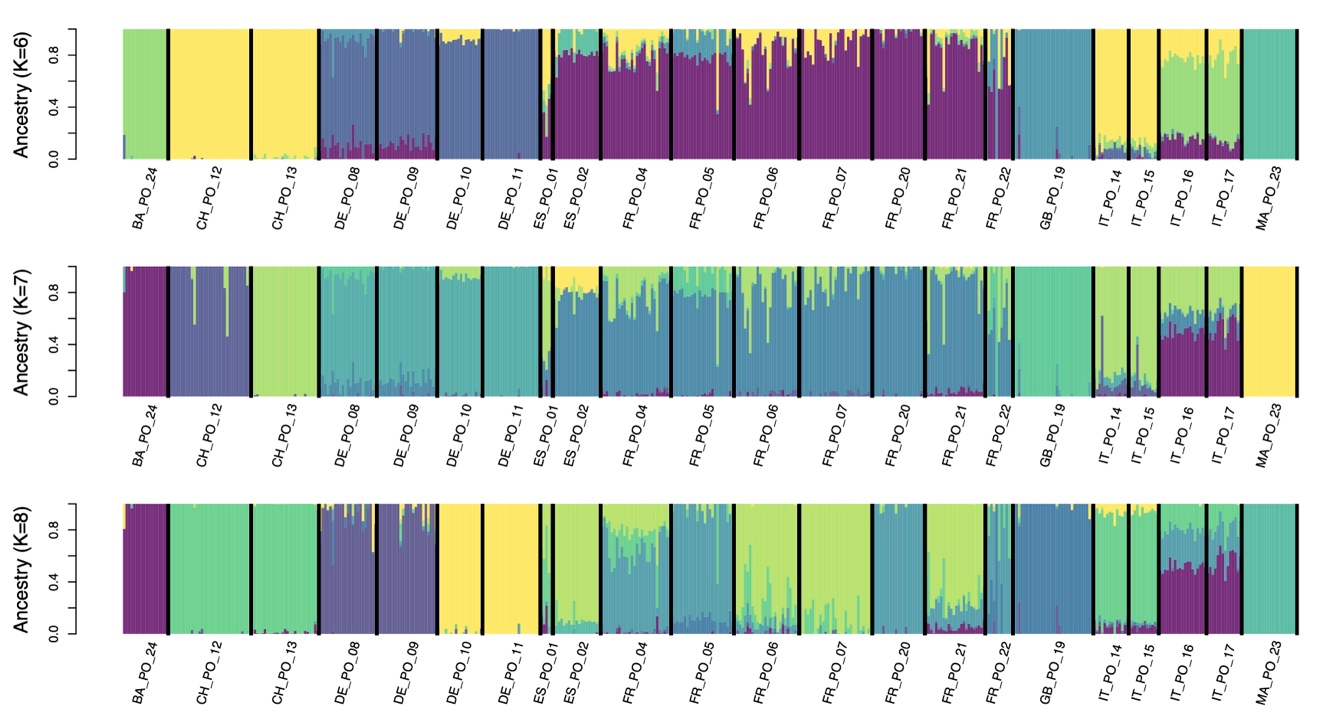

Figure S13. Level of admixture within and among individuals of *Populus nigra* based on the targeted capture SNP dataset. Colors reflect assignment probabilities (i.e. *Q* scores) to respective genetic group (*K* = 2–8). Population codes are explained in Table S1.

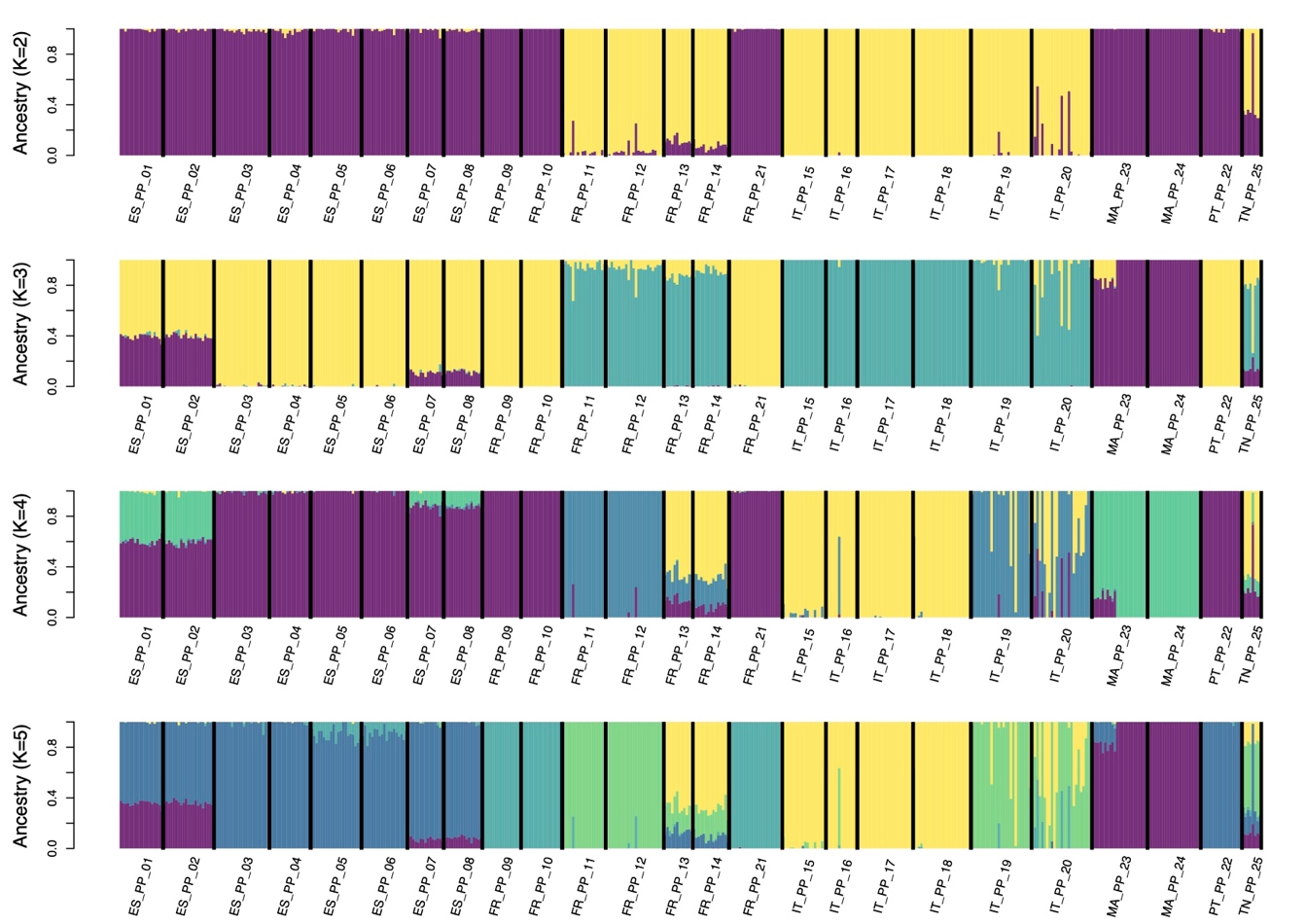

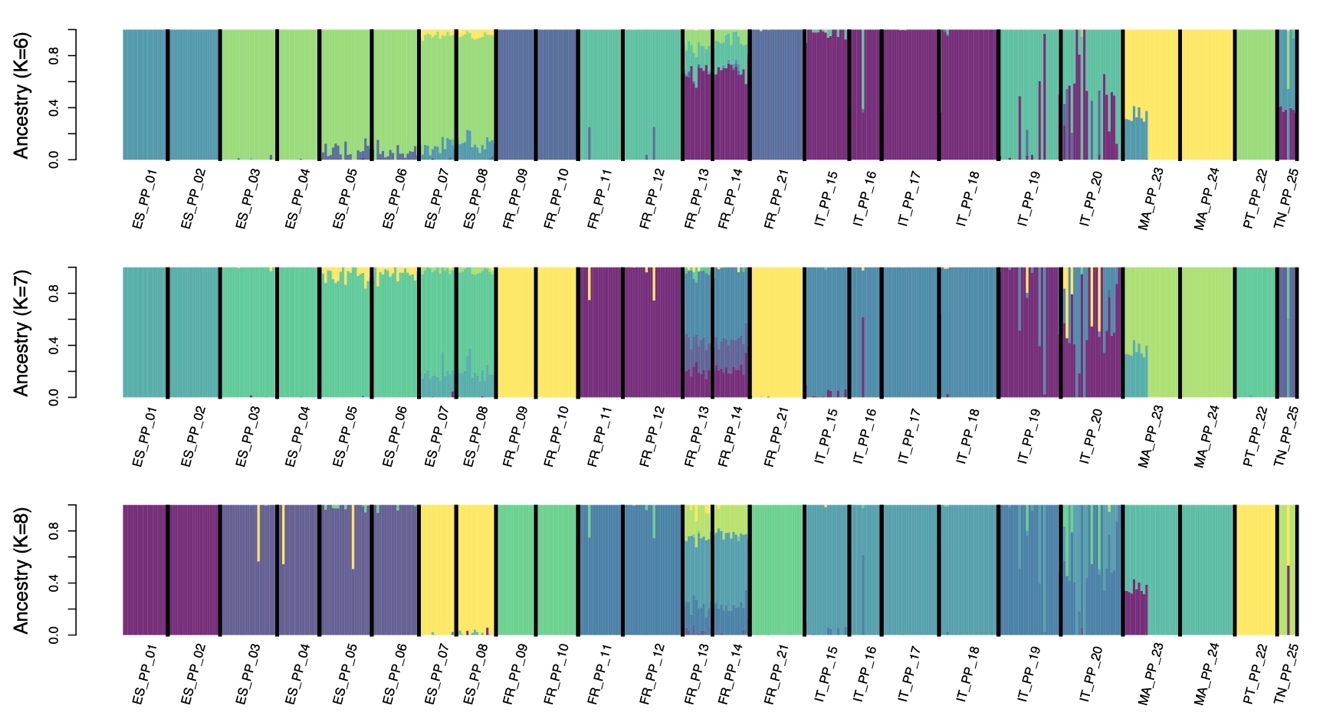

Figure S14. Level of admixture within and among individuals of *Pinus pinaster* based on the targeted capture SNP dataset. Colors reflect assignment probabilities (i.e. *Q* scores) to respective genetic group (*K* = 2–8). Population codes are explained in Table S1

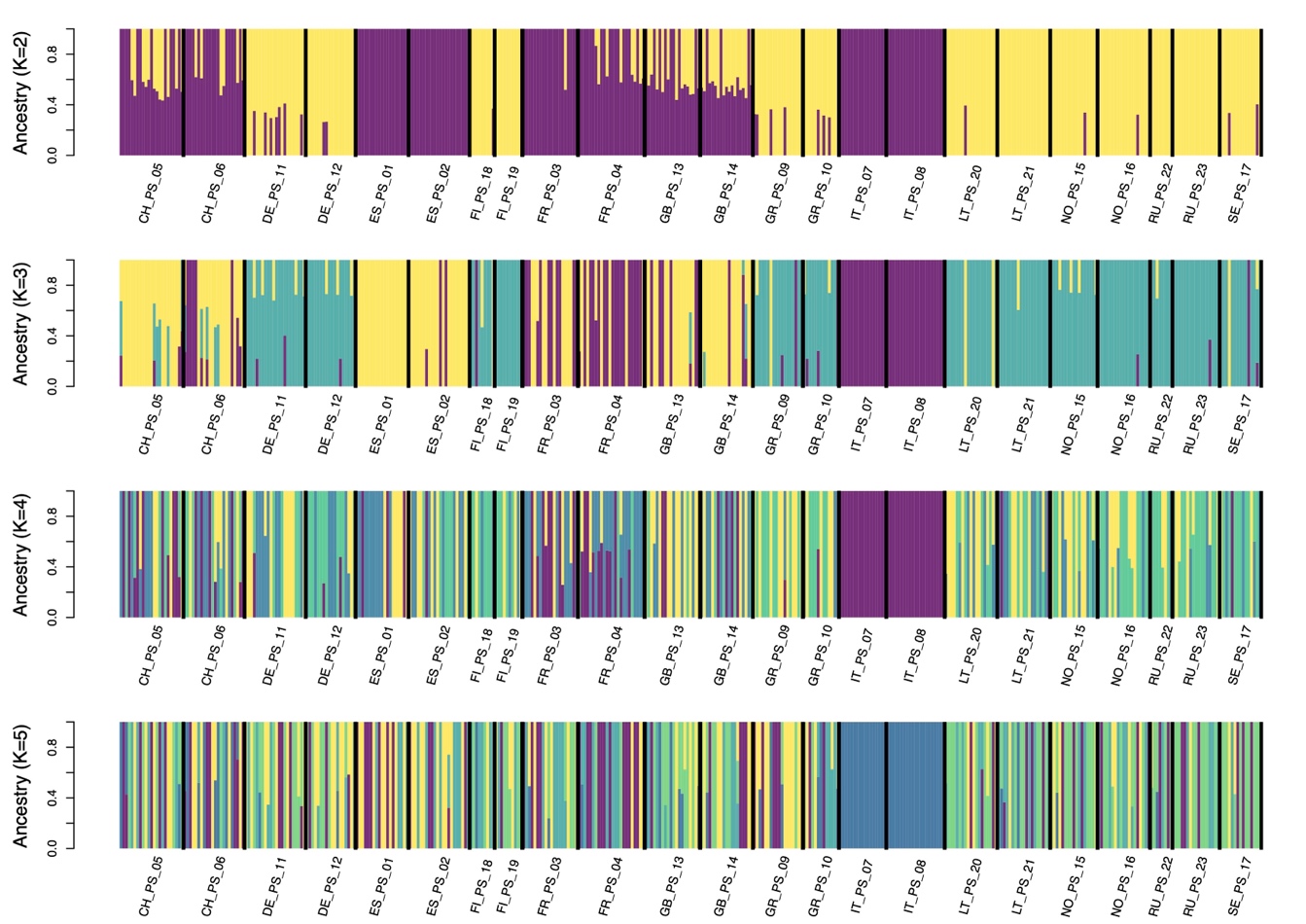

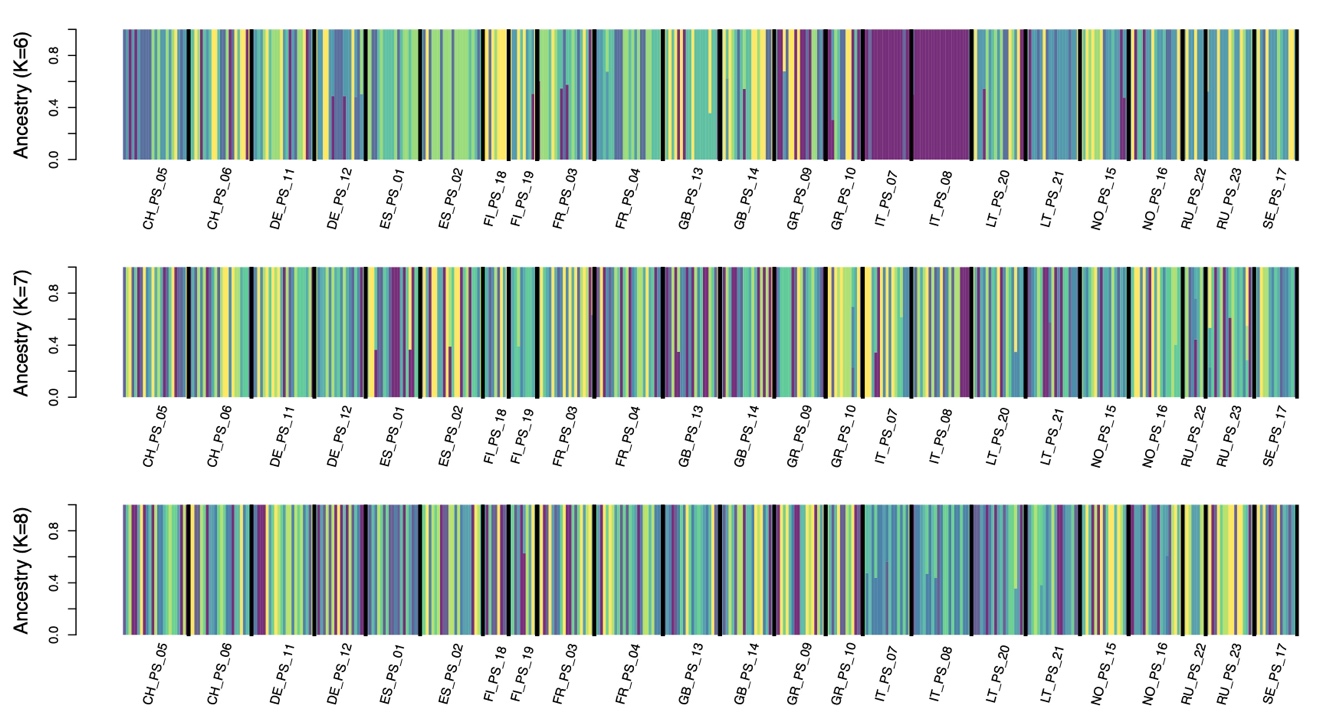

Figure S15. Level of admixture within and among individuals of *Pinus sylvestris* based on the targeted capture SNP dataset. Colors reflect assignment probabilities (i.e. *Q* scores) to respective genetic group (*K* = 2–8). Population codes are explained in Table S1.

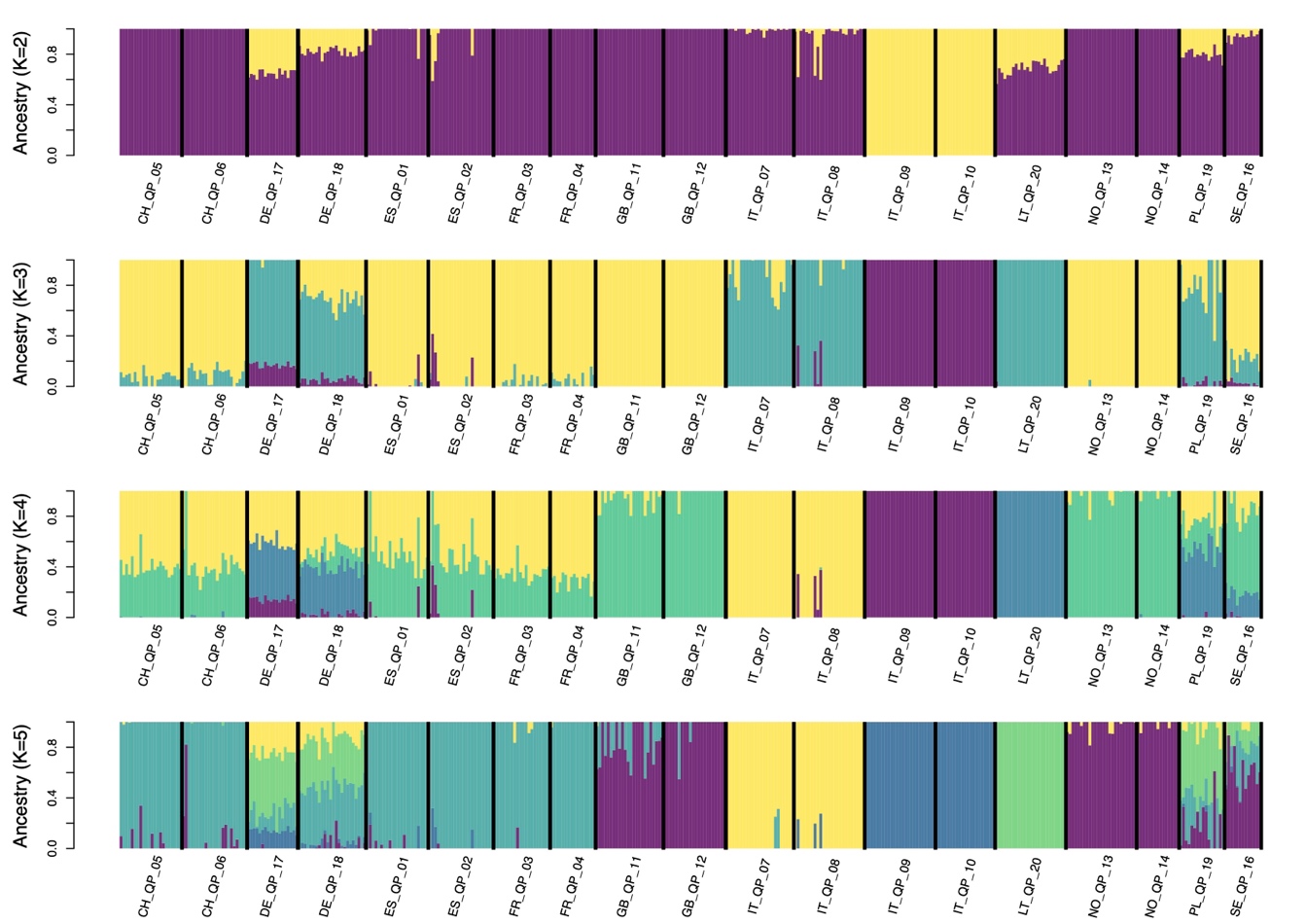

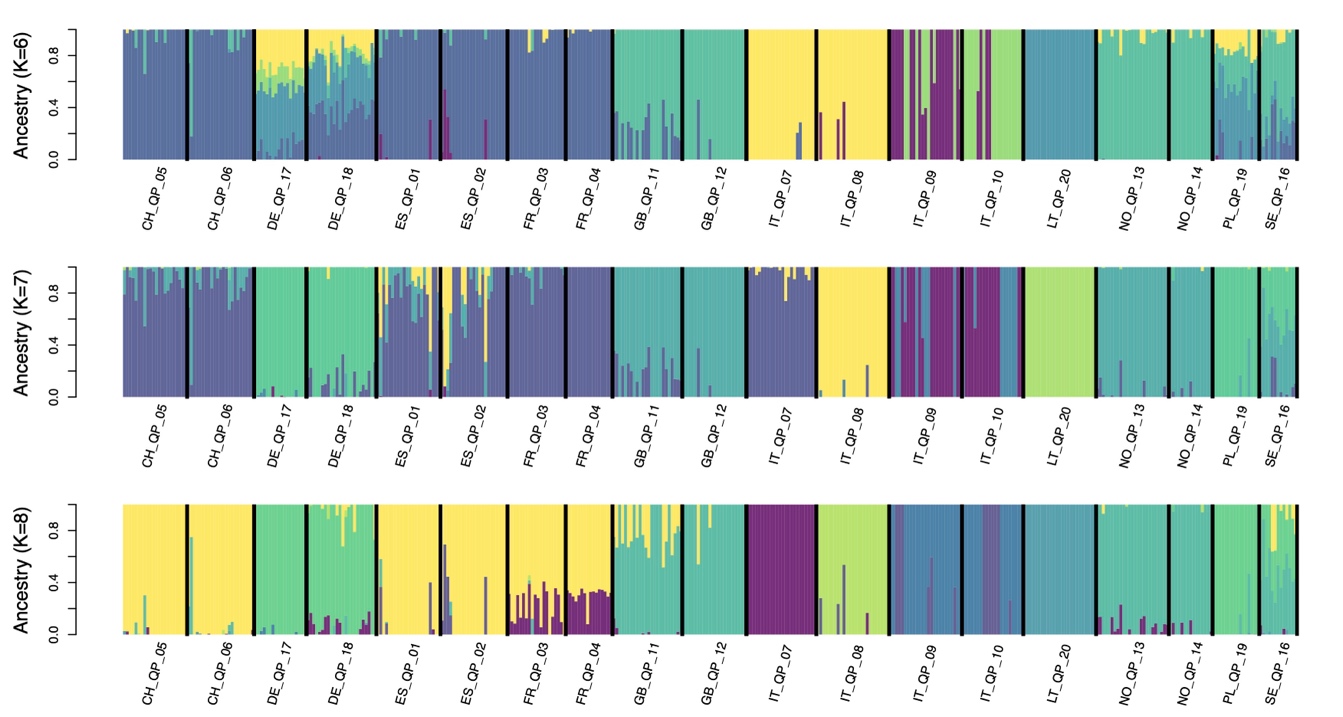

Figure S16. Level of admixture within and among individuals of *Quercus petraea* based on the targeted capture SNP dataset. Colors reflect assignment probabilities (i.e. *Q* scores) to respective genetic group (*K* = 2–8). Population codes are explained in Table S1.

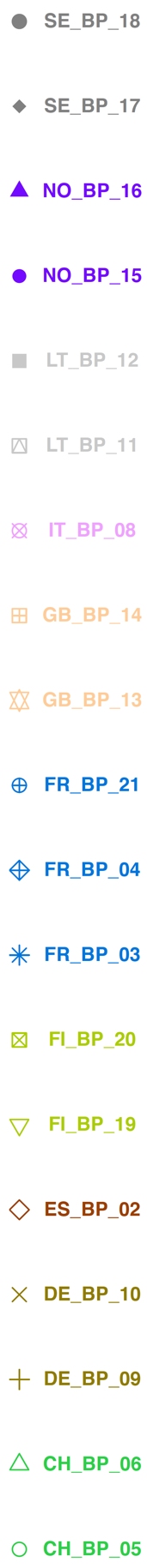

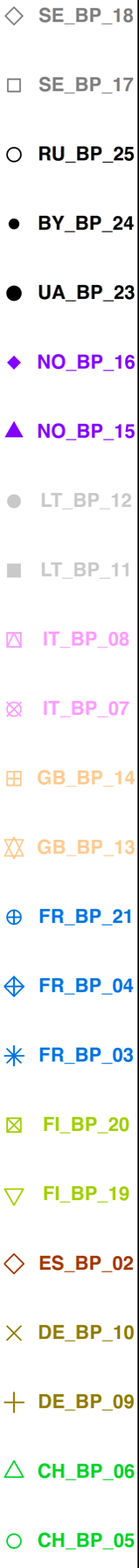

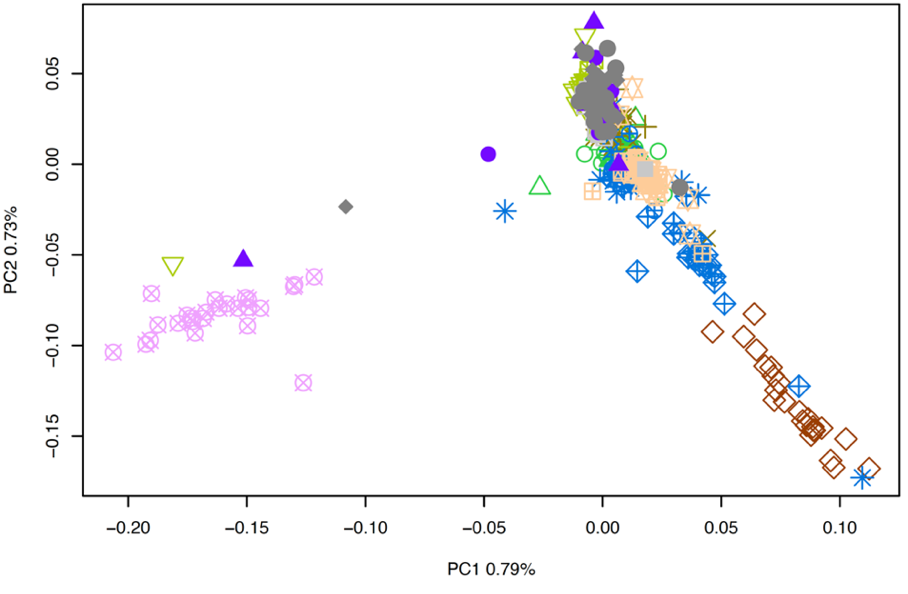

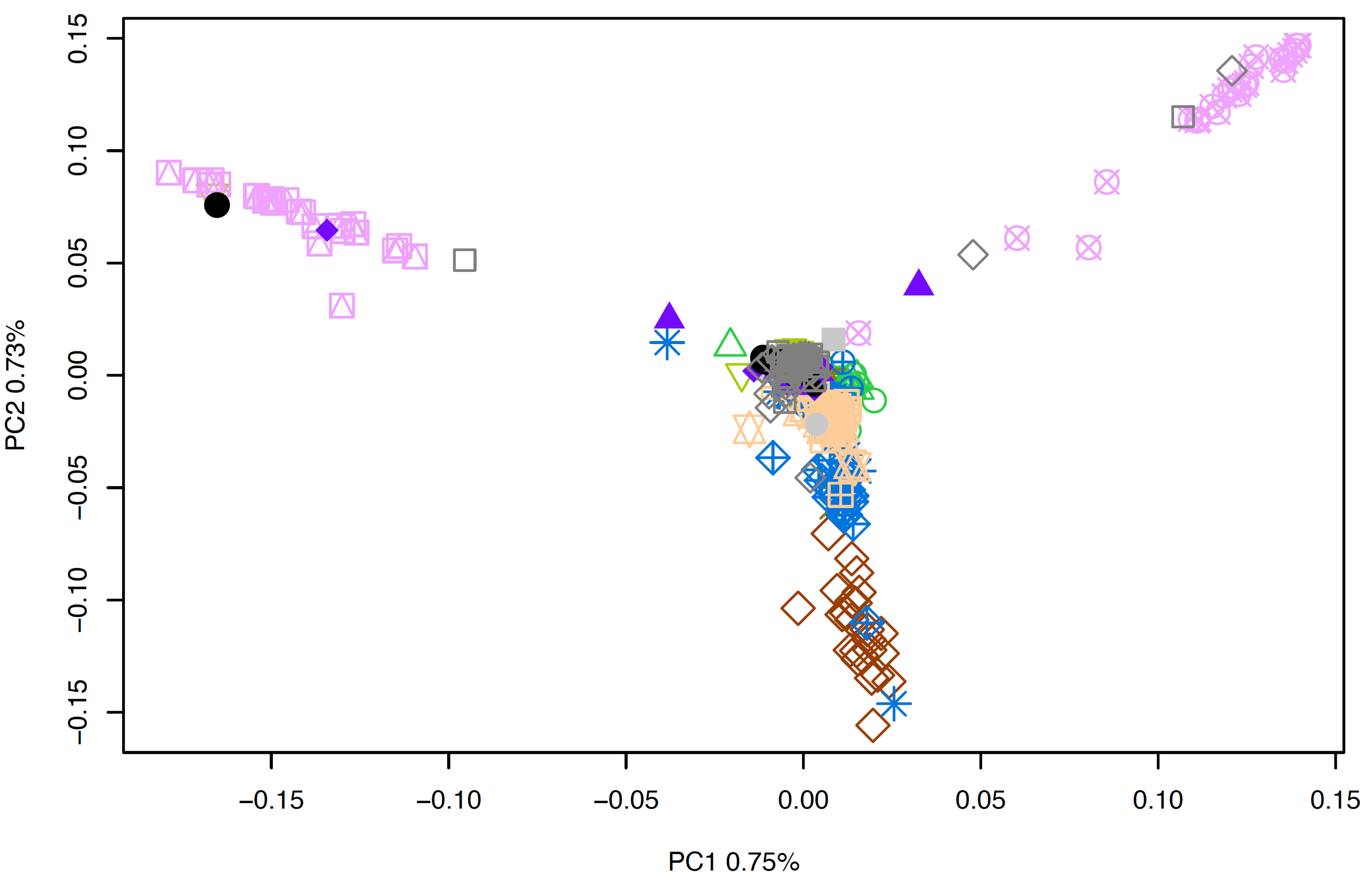

*Betula pendula*

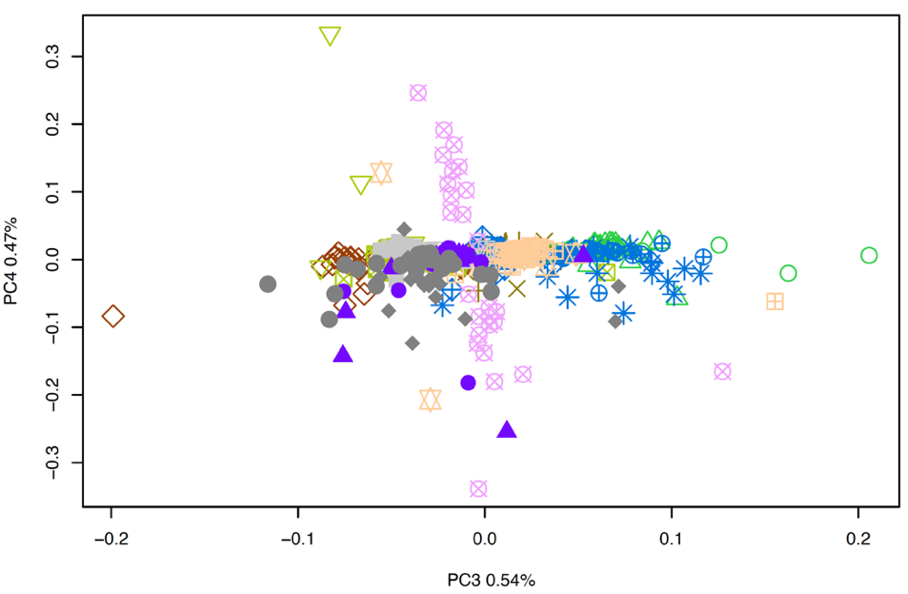

**Figure S17.** Principal component analysis of population structure of *Betula pendula*. Top panel is for all populations included in SNP set v.5.3. Middle and bottom panels show PC1 and 2 and PC3 and 4 of a PCA excluding outlier populations and individuals from top panel and focusing on Western Europe populations (SNP set v.6.3.1). Completely excluded populations are boxed. Leftmost most legend is for top panel, rightmost is for middle and bottom panels.

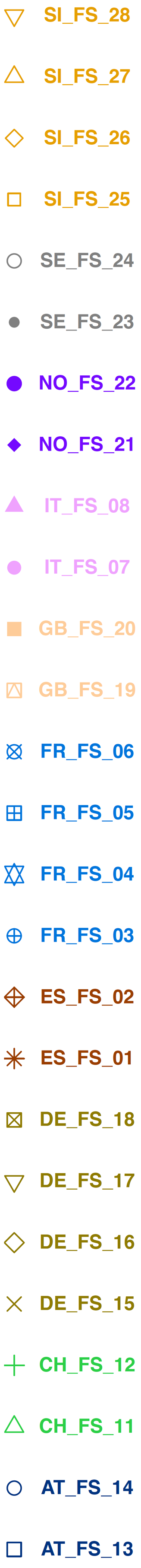

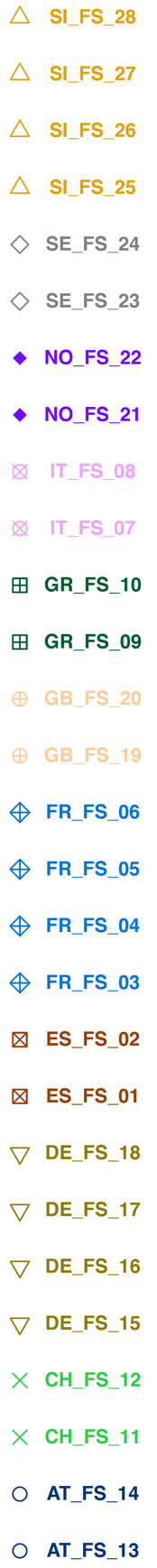

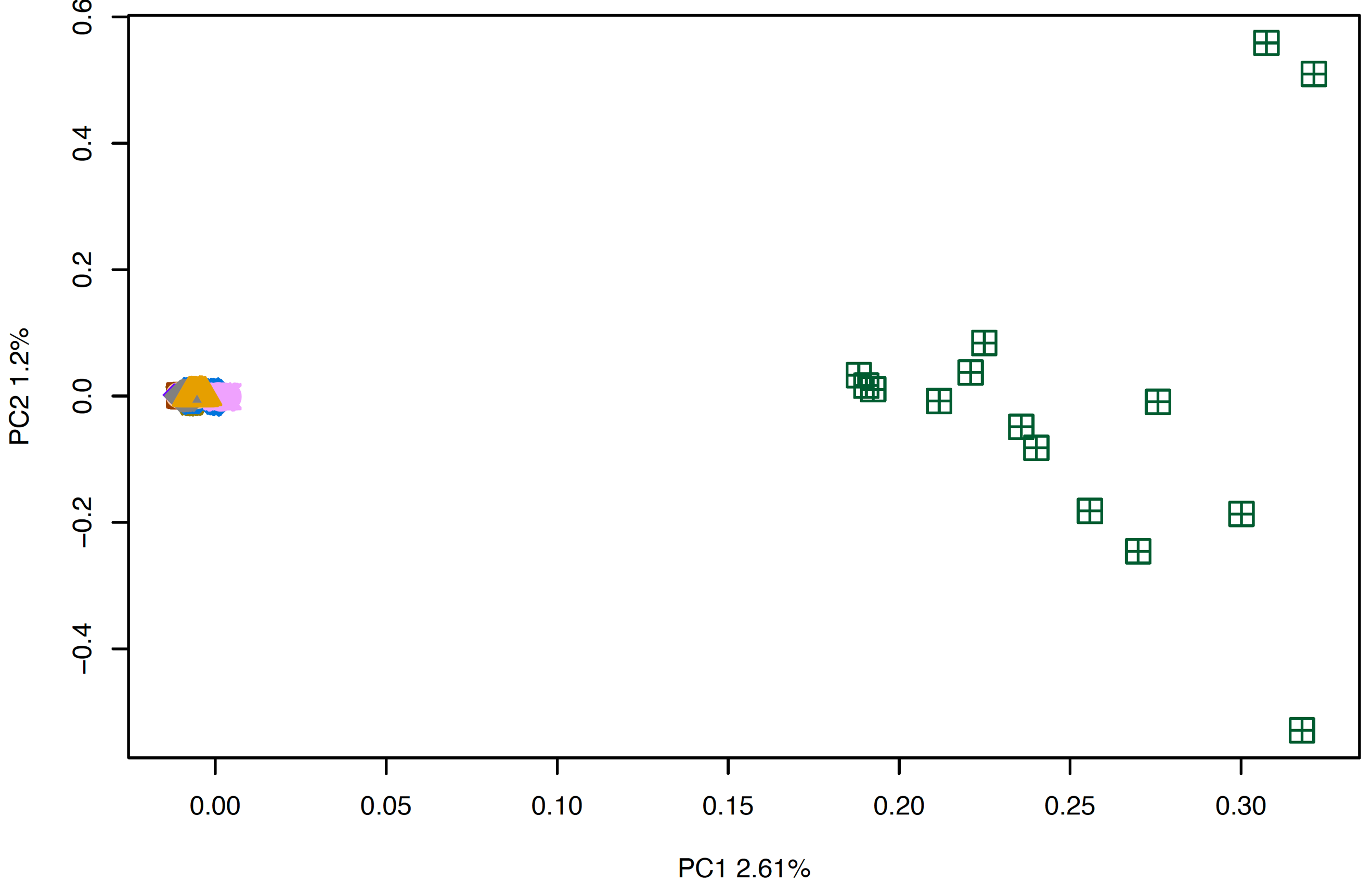

*Fagus sylvatica*

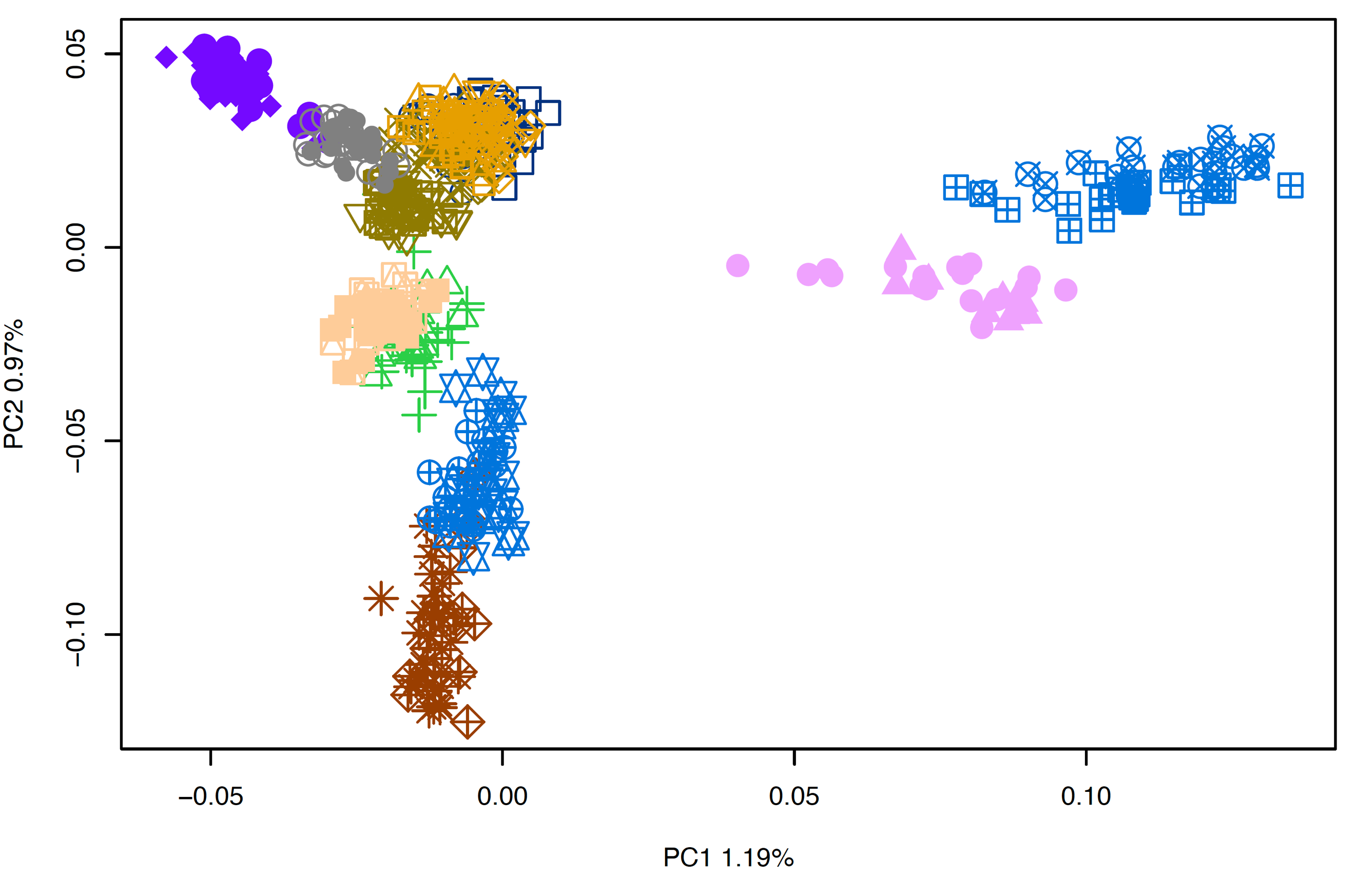

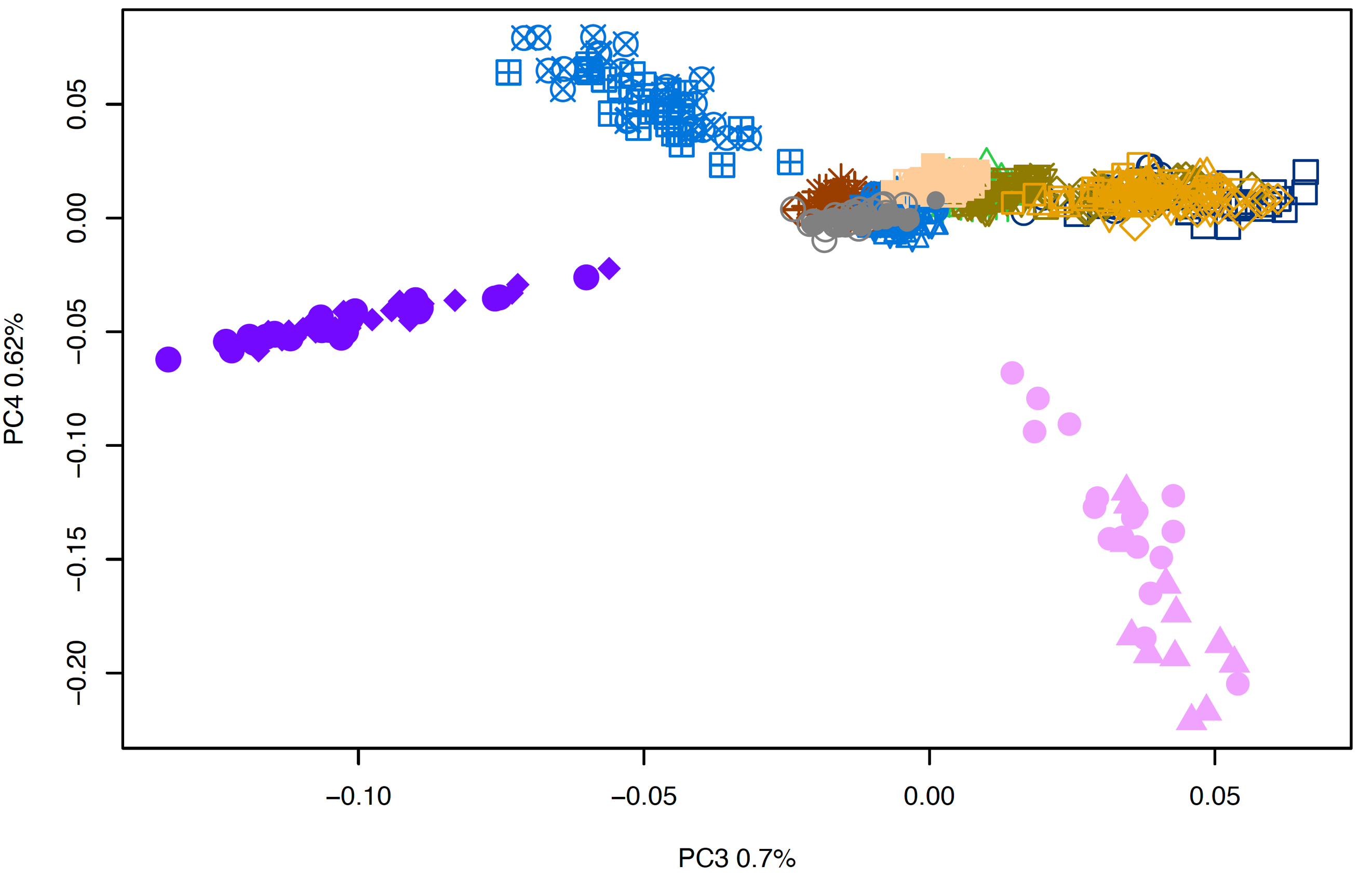

**Figure S18.** Principal component analysis of population structure of *Fagus sylvatica*. Top panel is for all populations included in SNP set v.5.3. Middle and bottom panels show PC1 and 2 and PC3 and 4 of a PCA excluding outlier populations and individuals from top panel and focusing on Western Europe populations (SNP set v.6.3.1). Completely excluded populations are boxed. Leftmost most legend is for top panel, rightmost is for middle and bottom panels.

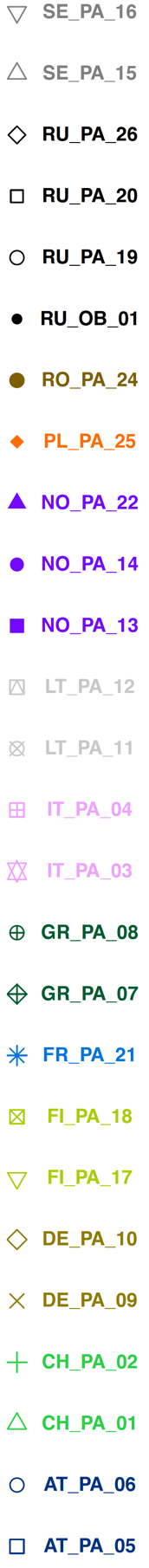

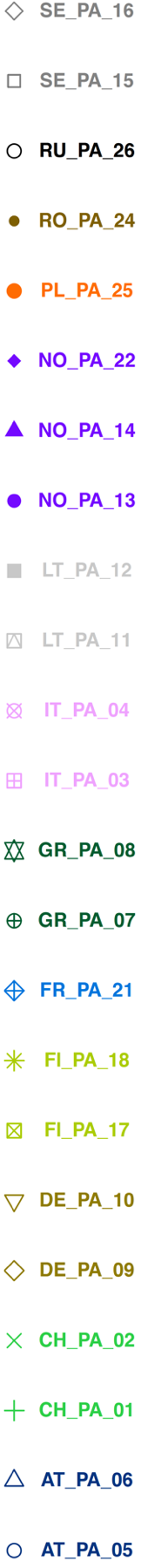

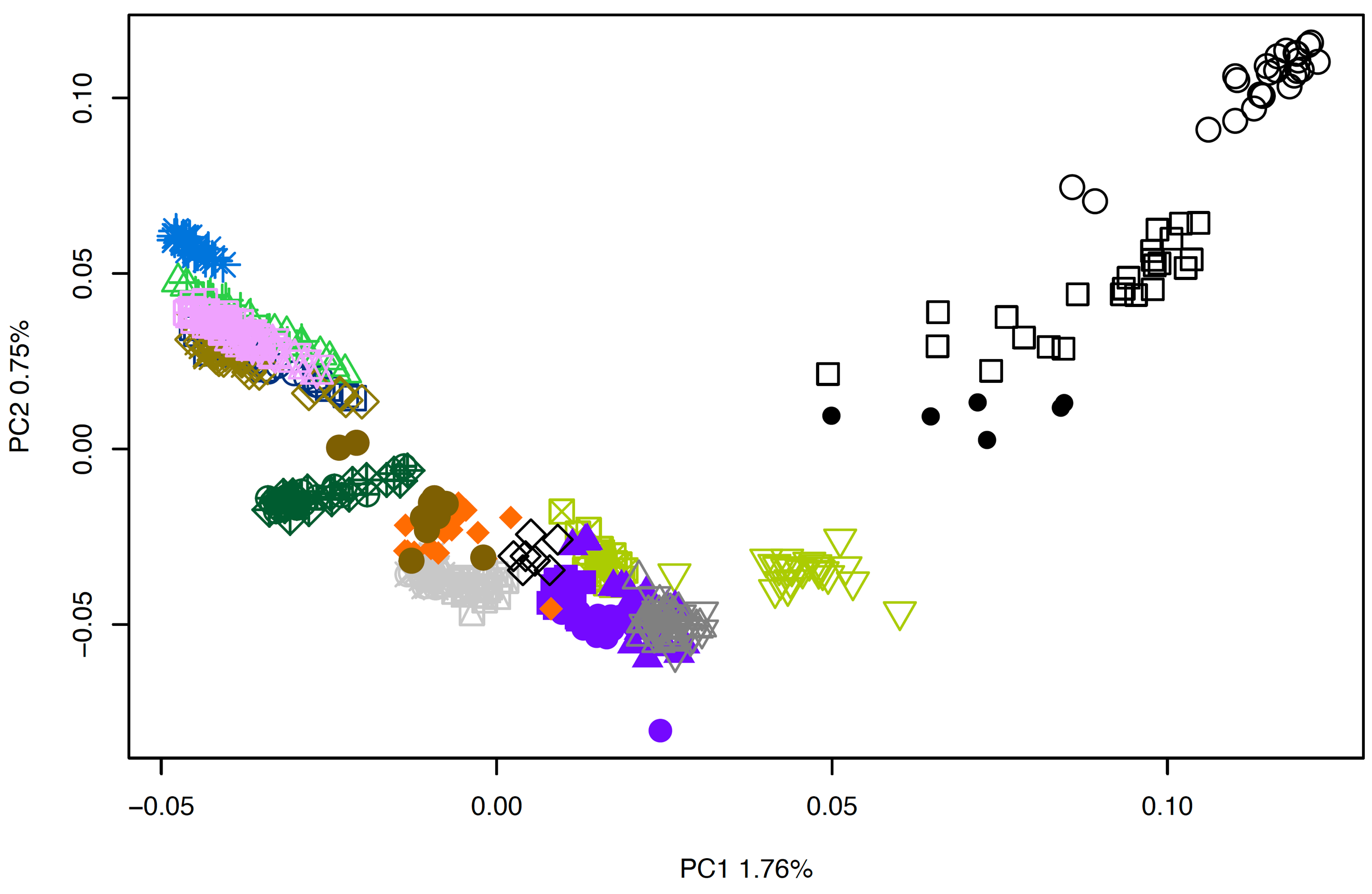

*Picea abies*

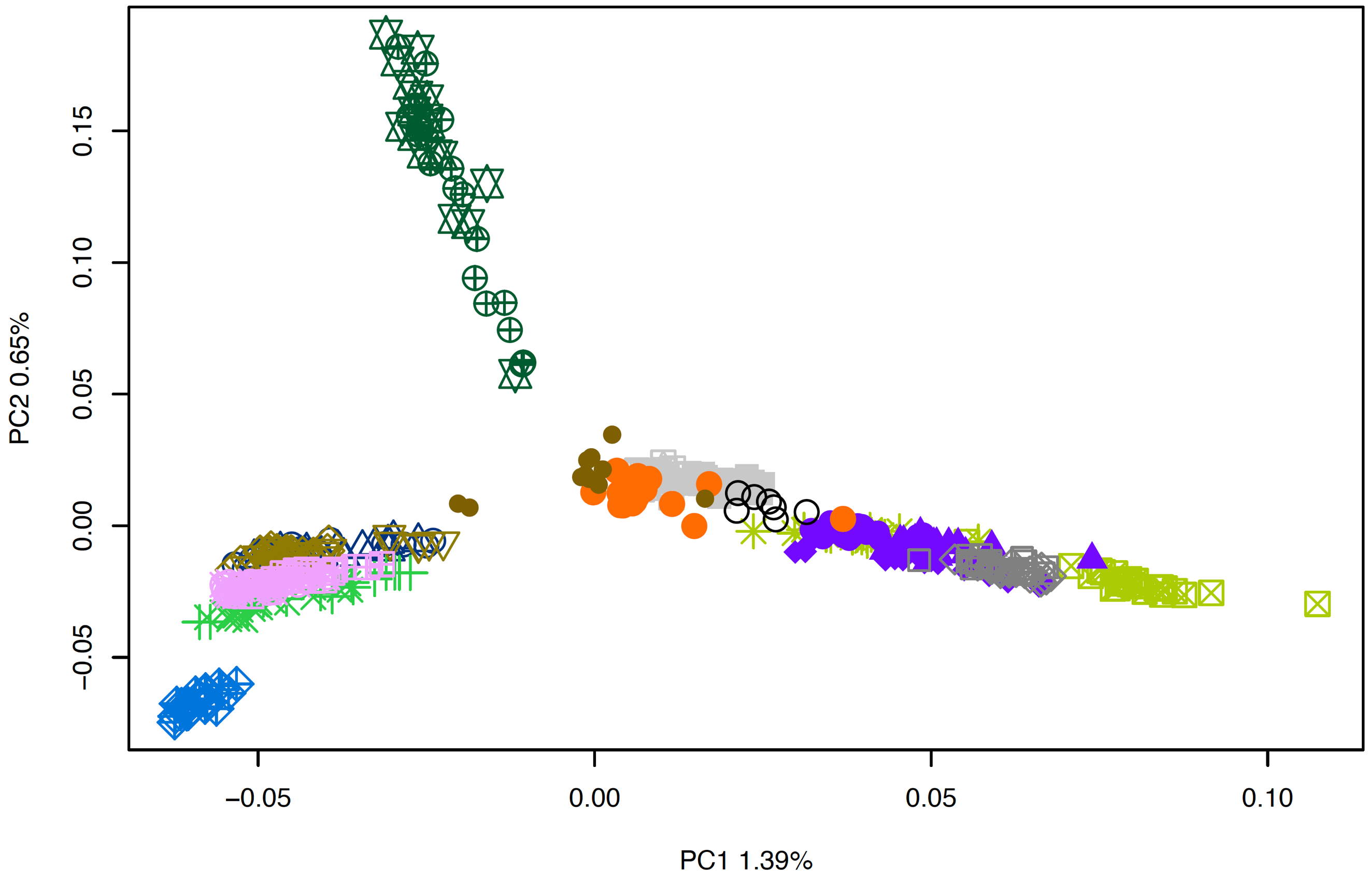

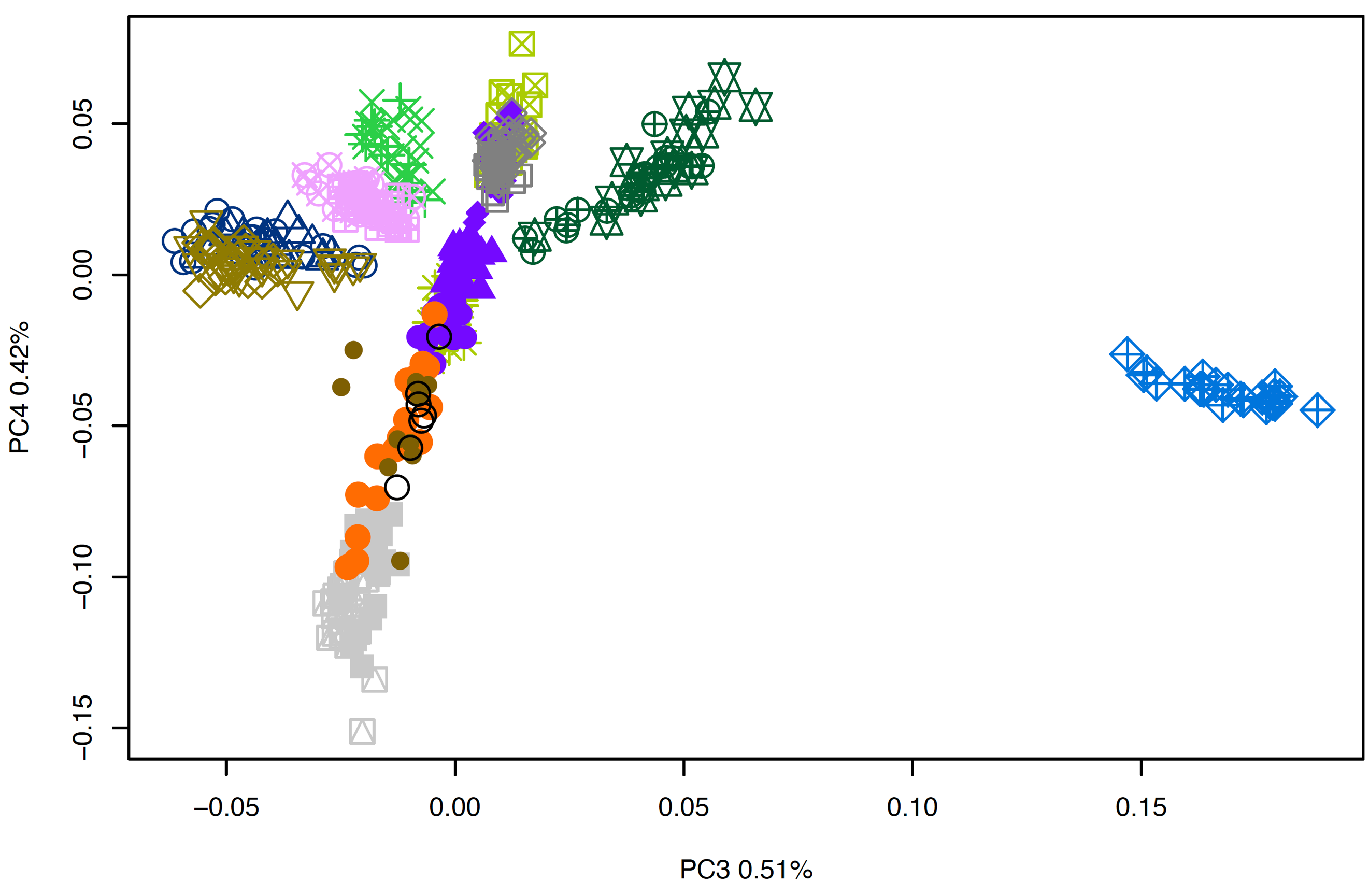

**Figure S19**. Principal component analysis of population structure of *Picea abies*. Top panel is for all populations included in SNP set v.5.3. Middle and bottom panels show PC1 and 2 and PC3 and 4 of a PCA excluding outlier populations and individuals from top panel and focusing on Western Europe populations (SNP set v.6.3.1). Completely excluded populations are boxed. Leftmost most legend is for top panel, rightmost is for middle and bottom panels.

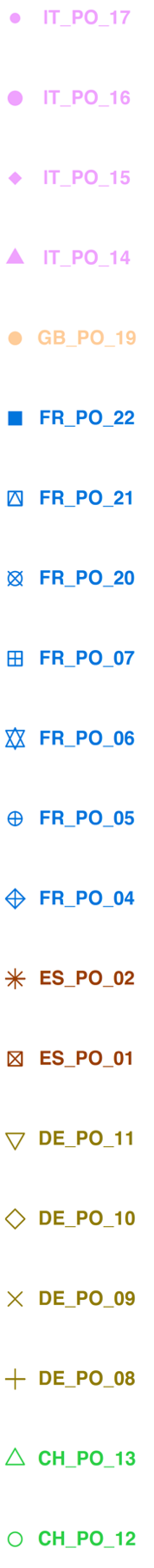

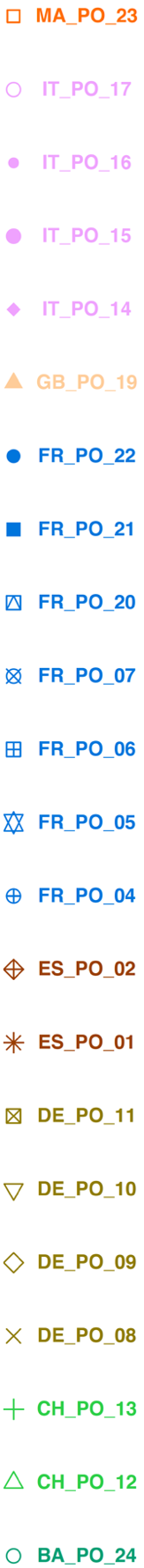

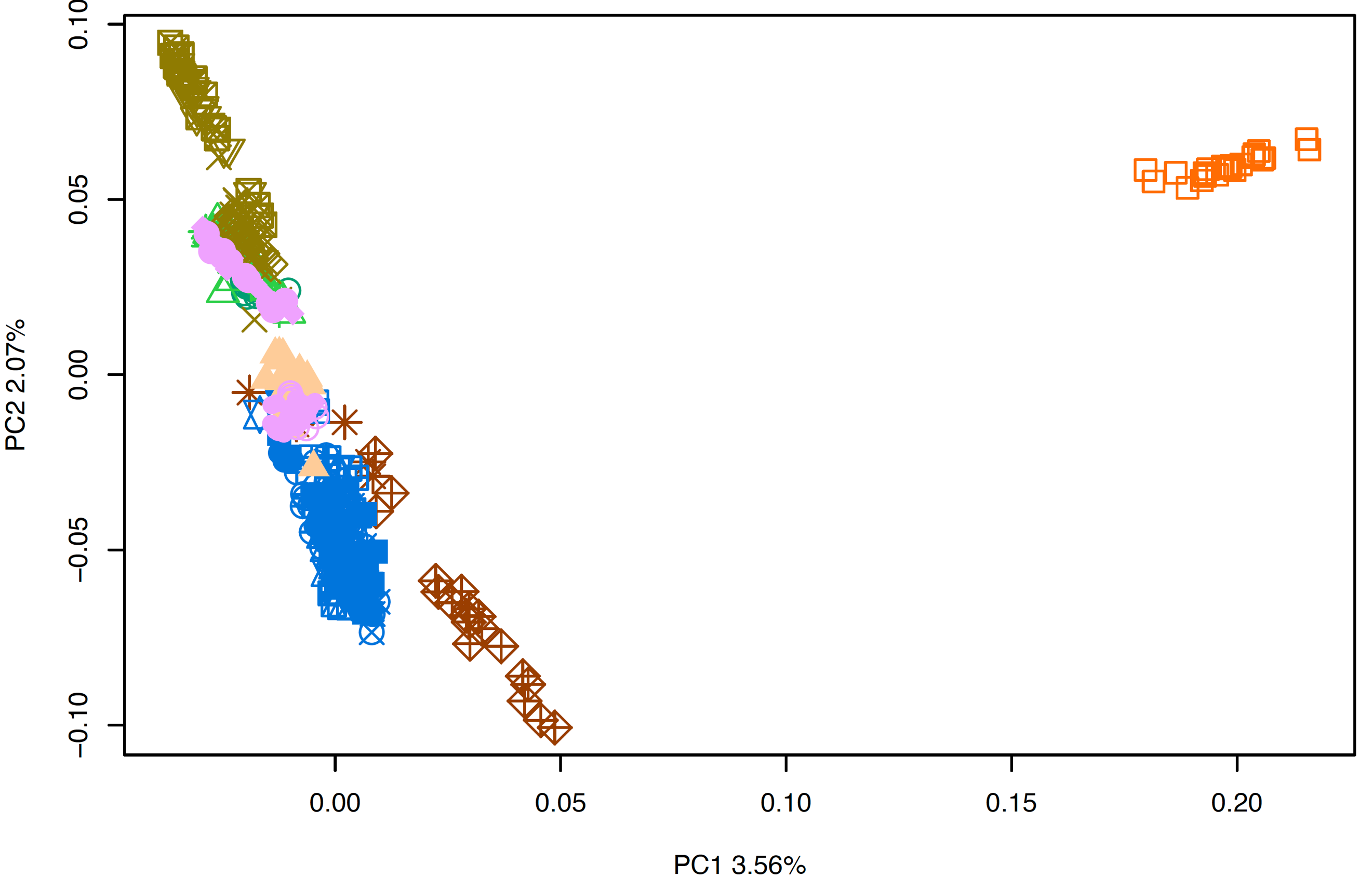

*Populus nigra*

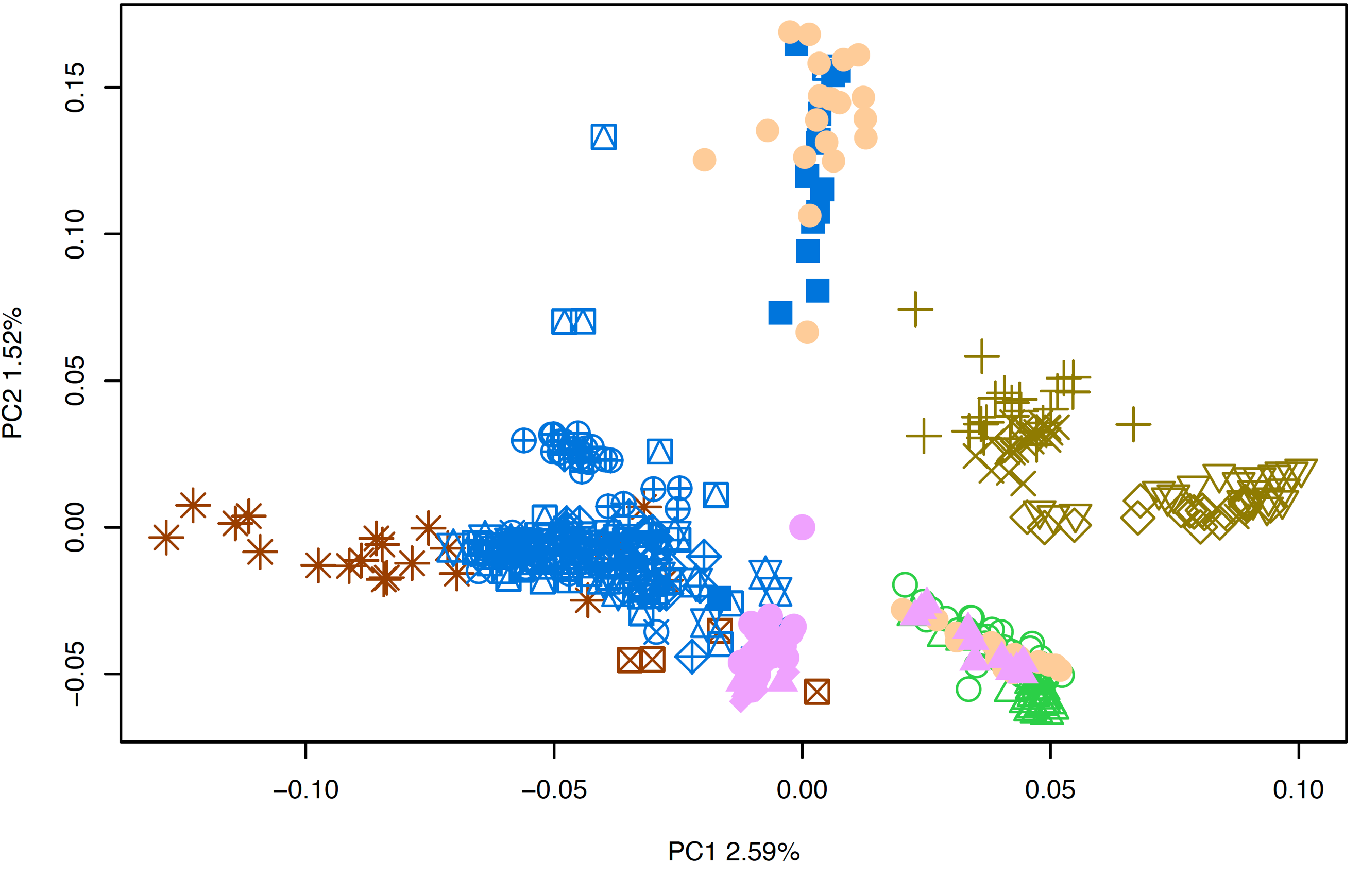

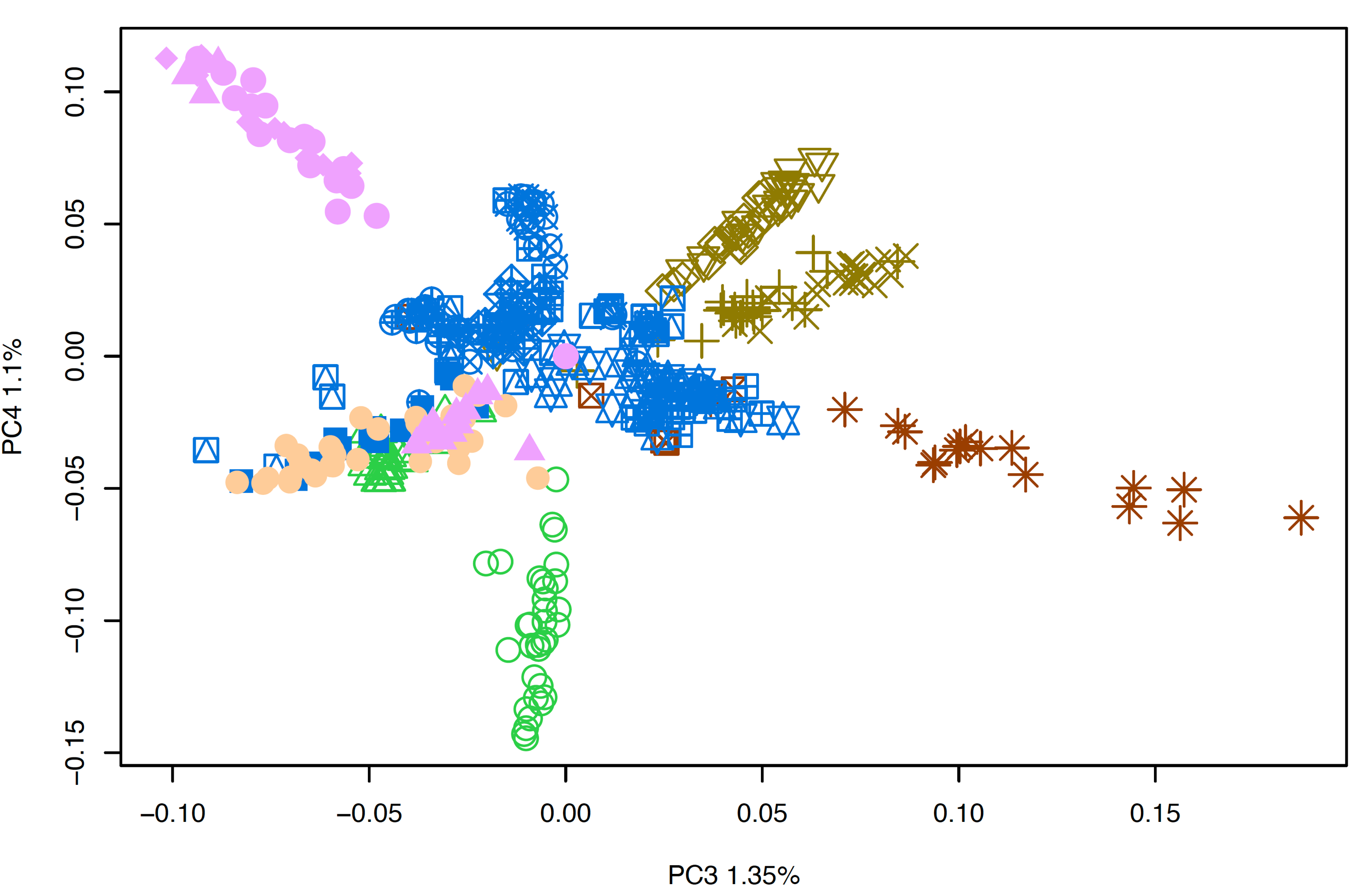

**Figure S20.** Principal component analysis of population structure of *Populus nigra*. Top panel is for all populations included in SNP set v.5.3. Middle and bottom panels show PC1 and 2 and PC3 and 4 of a PCA excluding outlier populations and individuals from top panel and focusing on Western Europe populations (SNP set v.6.3.1). Completely excluded populations are boxed. Leftmost most legend is for top panel, rightmost is for middle and bottom panels.

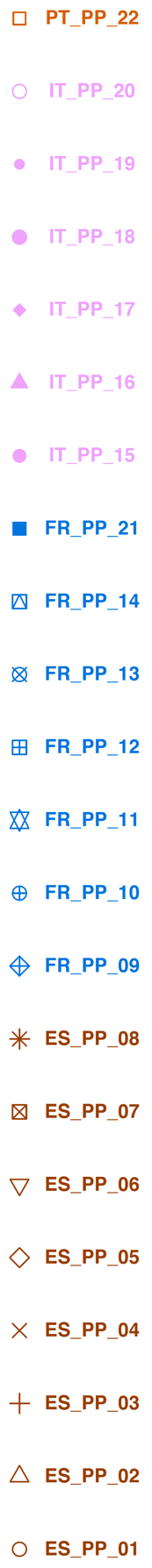

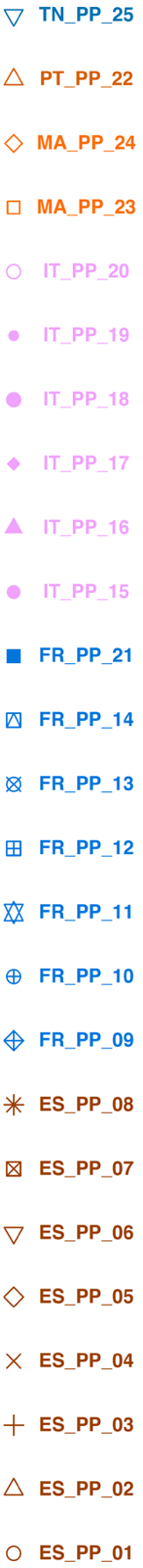

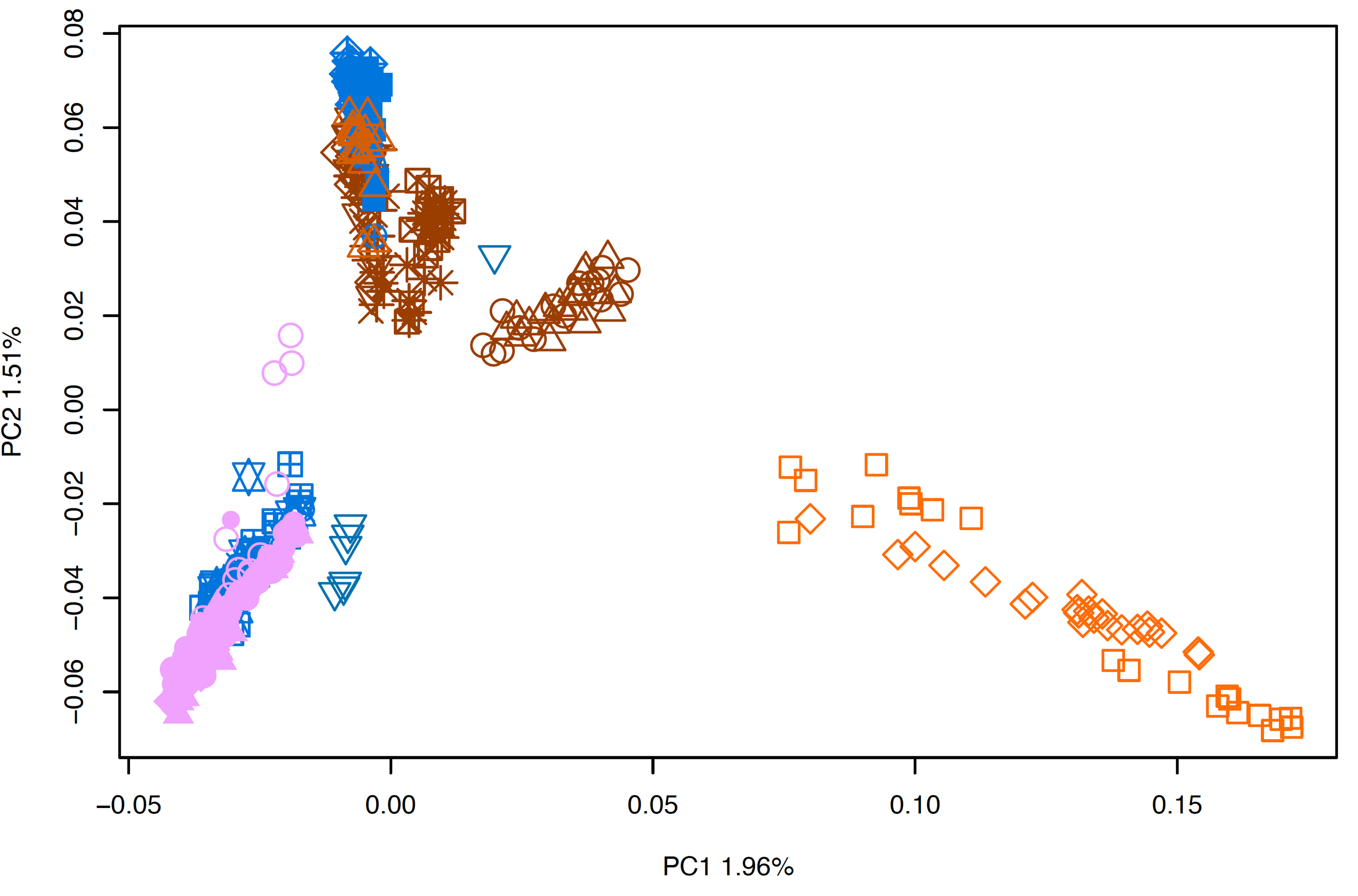

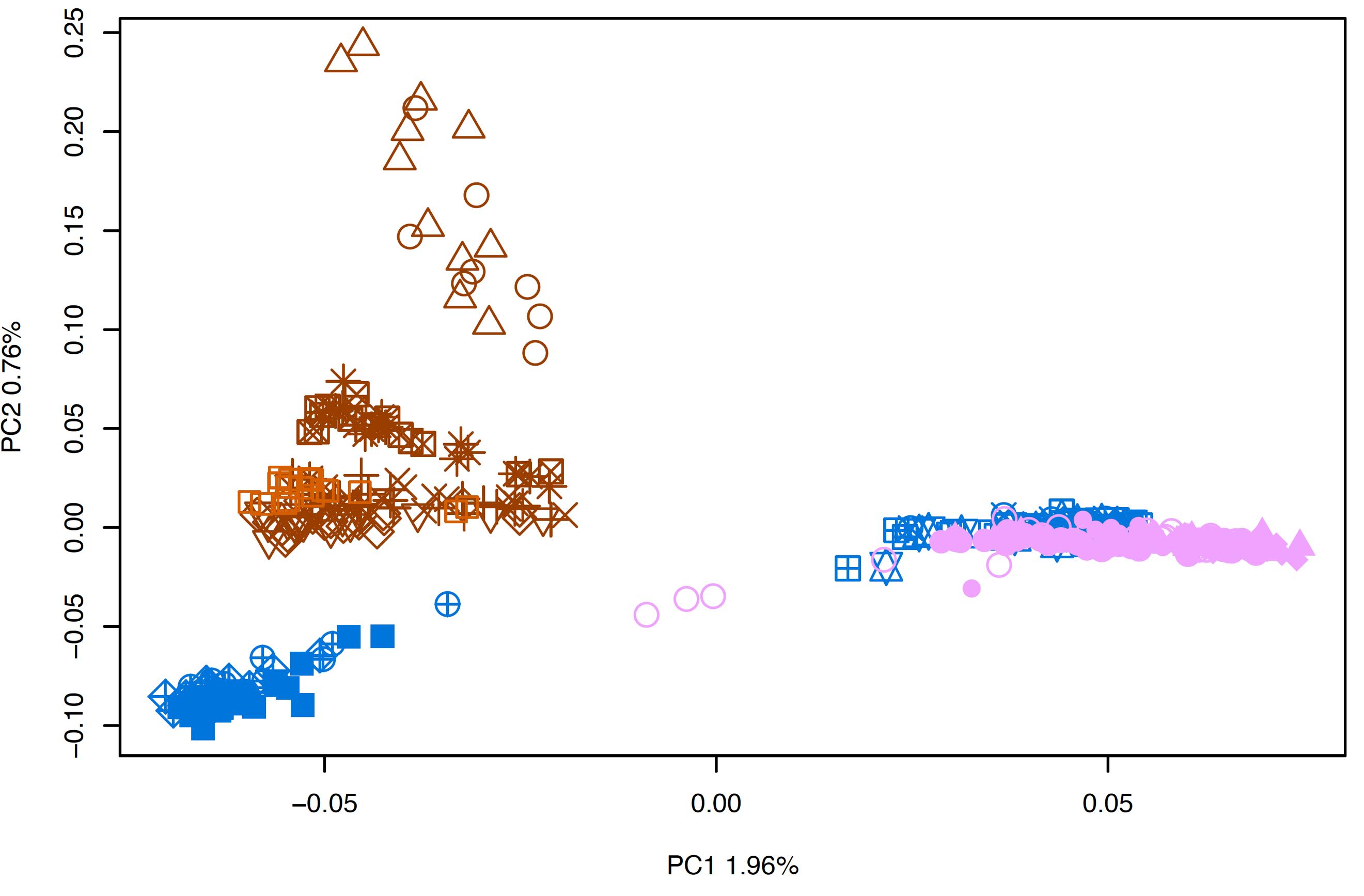

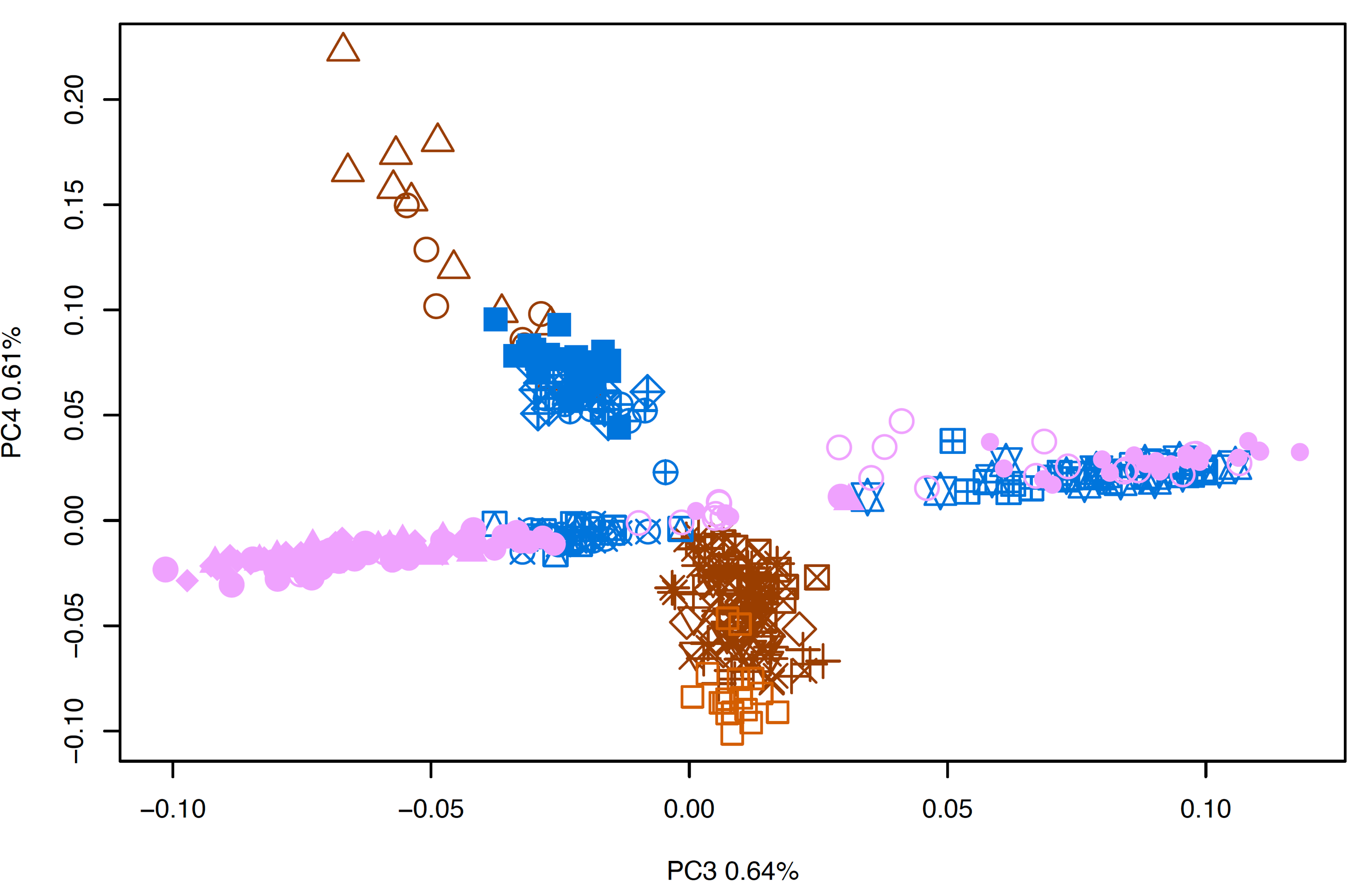

*Pinus pinaster*

**Figure S21.** Principal component analysis of population structure of *Pinus pinaster*. Top panel is for all populations included in SNP set v.5.3. Middle and bottom panels show PC1 and 2 and PC3 and 4 of a PCA excluding outlier populations and individuals from top panel and focusing on Western Europe populations (SNP set v.6.3.1). Completely excluded populations are boxed. Leftmost most legend is for top panel, rightmost is for middle and bottom panels.

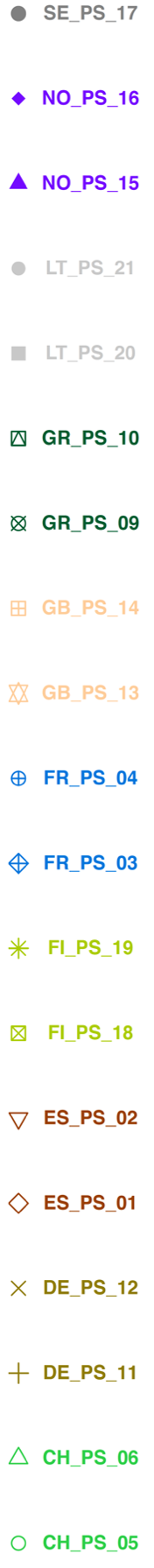

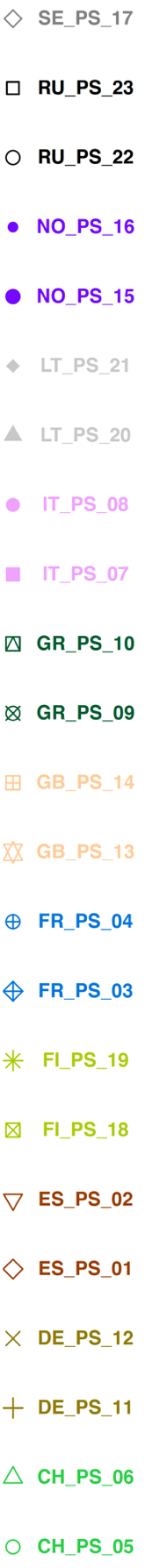

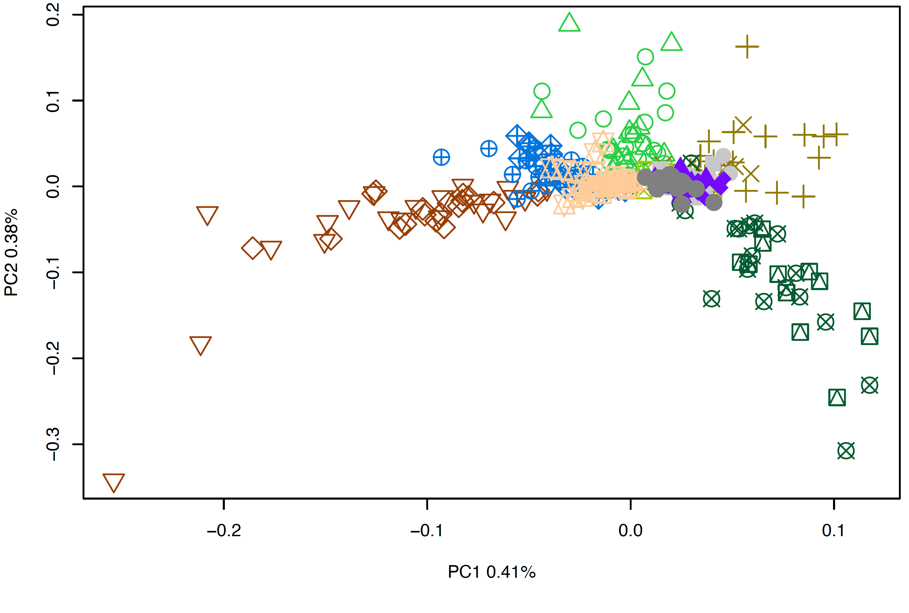

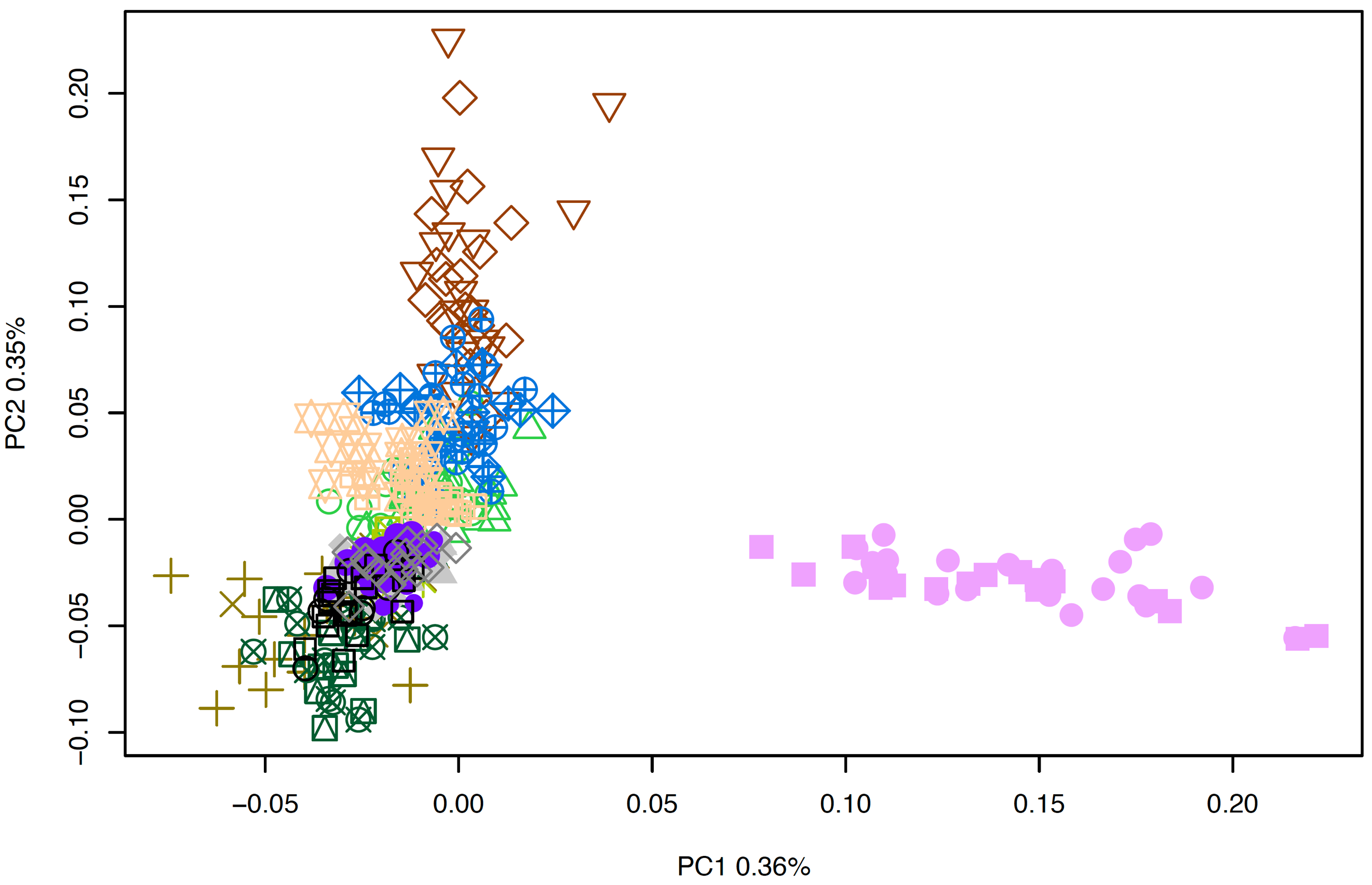

*Pinus sylvestris*

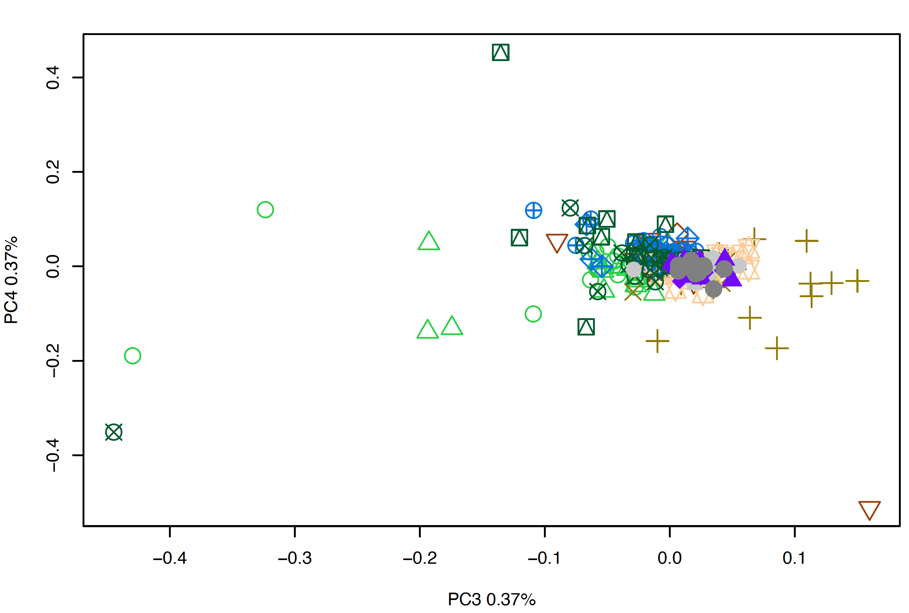

**Figure S22.** Principal component analysis of population structure of *Pinus sylvestris*. Top panel is for all populations included in SNP set v.5.3. Middle and bottom panels show PC1 and 2 and PC3 and 4 of a PCA excluding outlier populations and individuals from top panel and focusing on Western Europe populations (SNP set v.6.3.1). Completely excluded populations are boxed. Leftmost most legend is for top panel, rightmost is for middle and bottom panels.

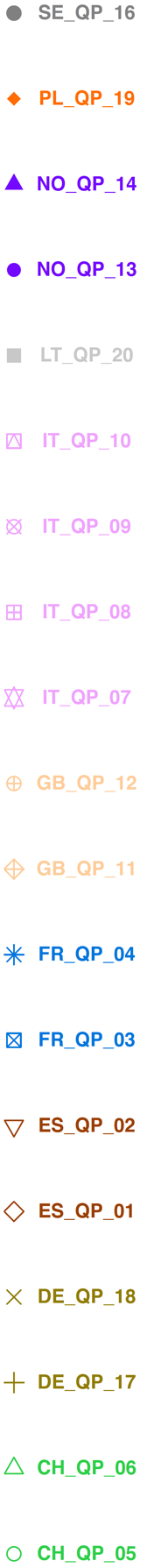

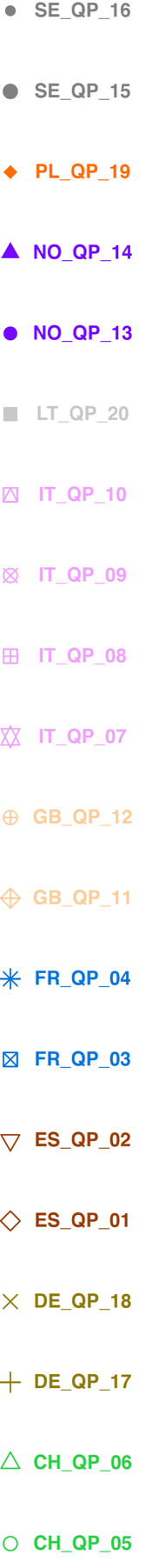

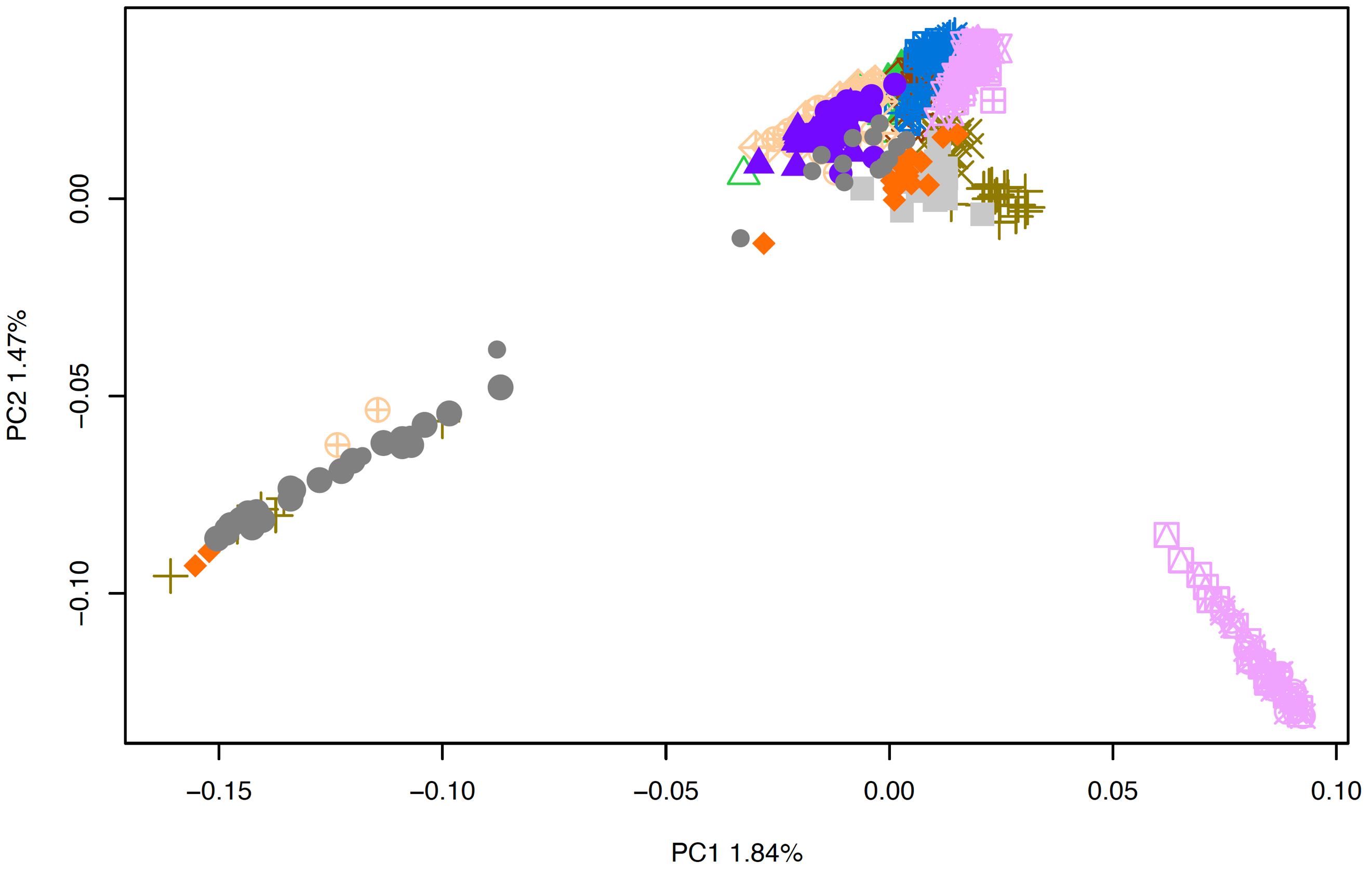

*Quercus petraea*

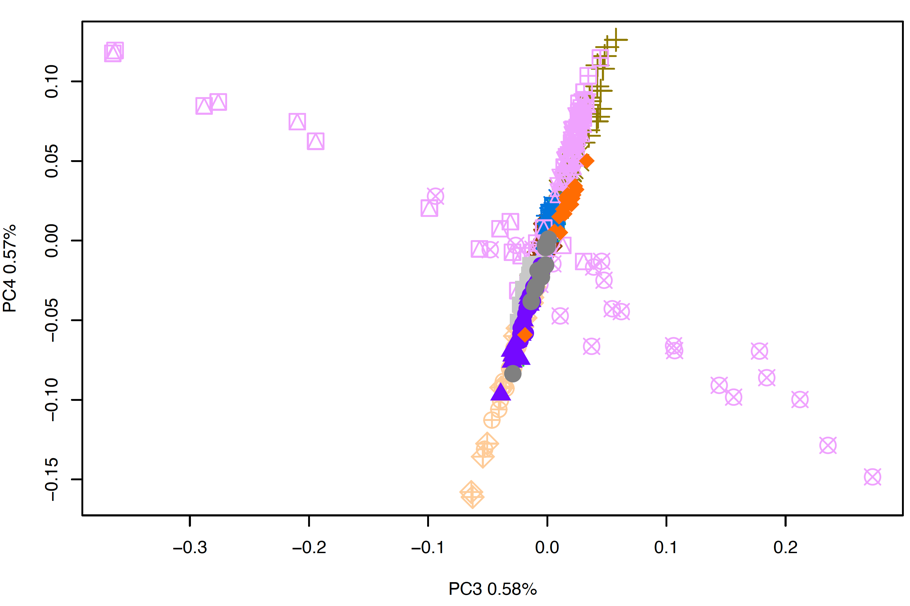

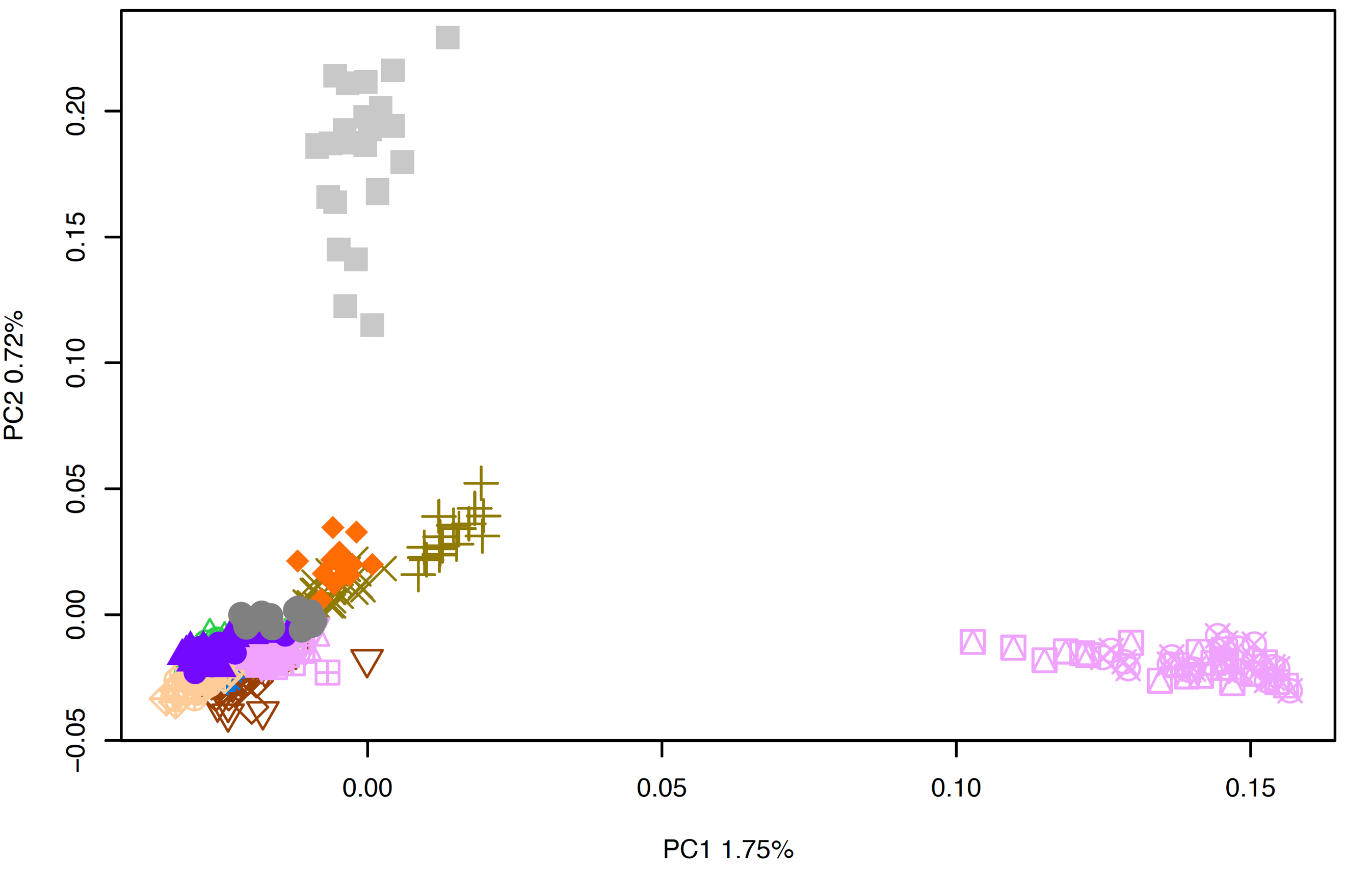

**Figure SX:** **Principal component analysis of population**

**Figure S23.** Principal component analysis of population structure of *Quercus petraea*. Top panel is for all populations included in SNP set v.5.3. Middle and bottom panels show PC1 and 2 and PC3 and 4 of a PCA excluding outlier populations and individuals from top panel and focusing on Western Europe populations (SNP set v.6.3.1). Completely excluded populations are boxed. Leftmost most legend is for top panel, rightmost is for middle and bottom panels.

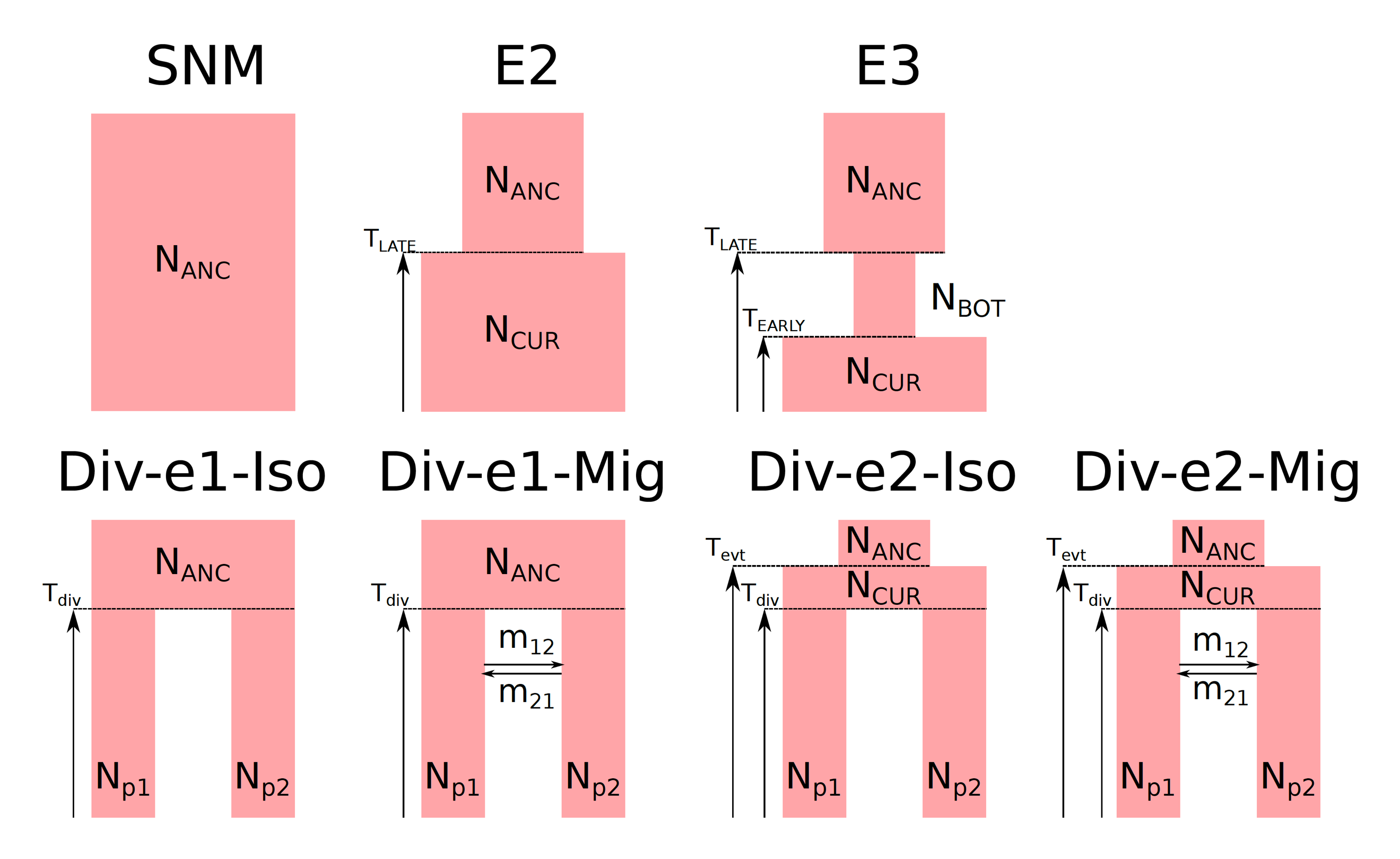

**Figure S24.** Schematics of the Fastsimcoal 2 models used in this study and their parameters. The blocks represent the evolution of populations with their lengths along the horizontal axis reflecting the population size and the time on the vertical axis from past (top) to present (bottom). Model abbreviations: SNM = Standard Neutral Model, E2 = 2 epoch model, E3 = 3 epoch model, Div-e1-Iso and Div-e2-Iso = Divergence model of two isolated populations both deriving from the same ancestral population that didn't or did experience a single demographic event in the past respectively, Div-e1-Mig and Div-e2-Mig = Divergence model of two populations with migration between both deriving from the same ancestral population that didn't or did experience a single demographic event in the past respectively. Parameter abbreviations: N_ANC_, N_CUR_, N_BOT_, N_p1_, N_p2_: effective population size of respectively the ancestral, current, bottleneck, diverged population 1 or diverged population 2. T_LATE_ T_EARLY_: time of the latest or earliest (backward in time) demographic event. T_evt_ and T_div_: time of the demographic or divergence event. m_12_ m_21_: migration rate from population 1 to 2 and 2 to 1 respectively (backward in time).

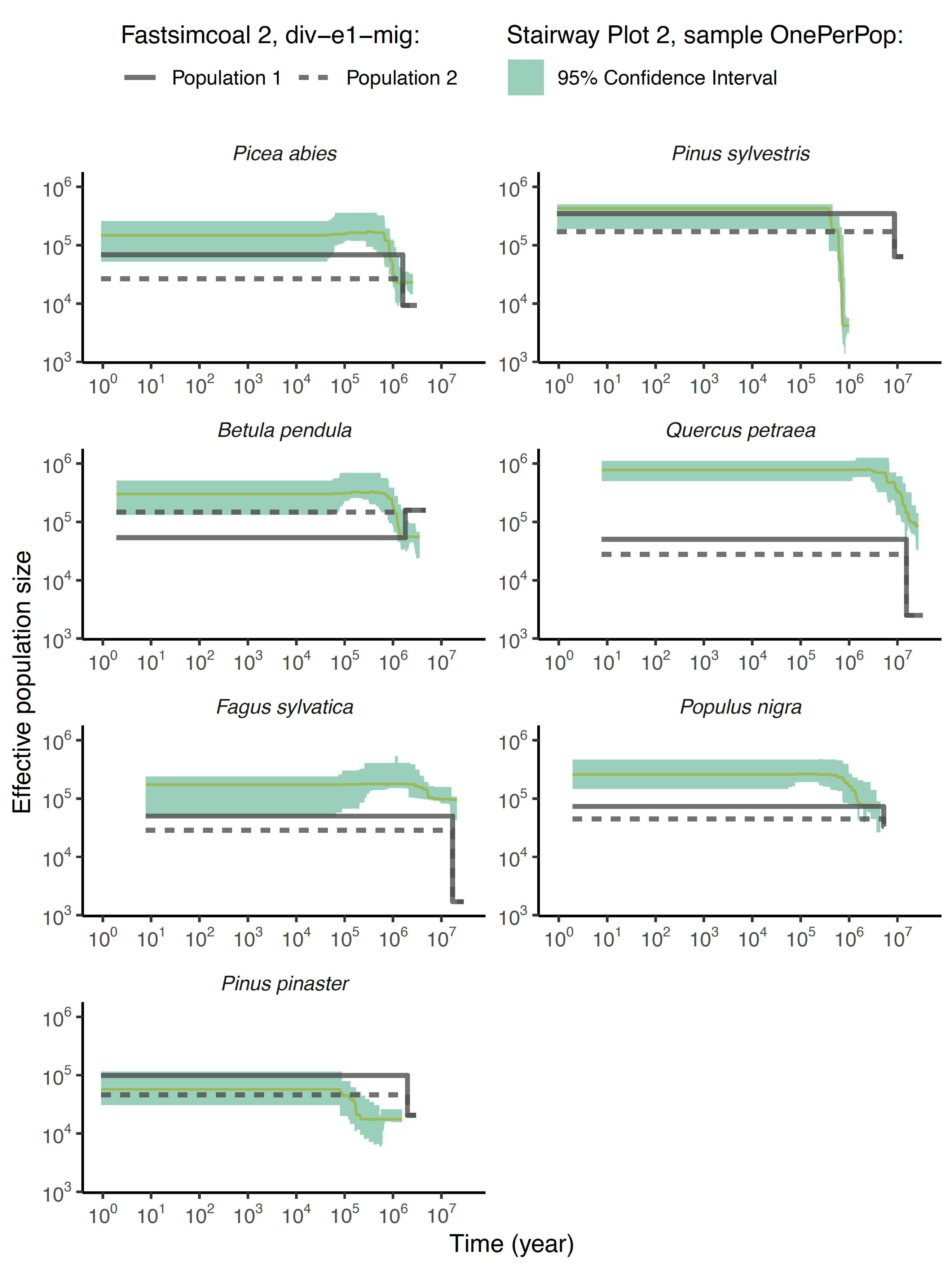

**Figure S25.** Comparisons of the results obtained with Stairway Plot 2 and the divergence models (div-e1-mig) implemented with fastsimcoal2.

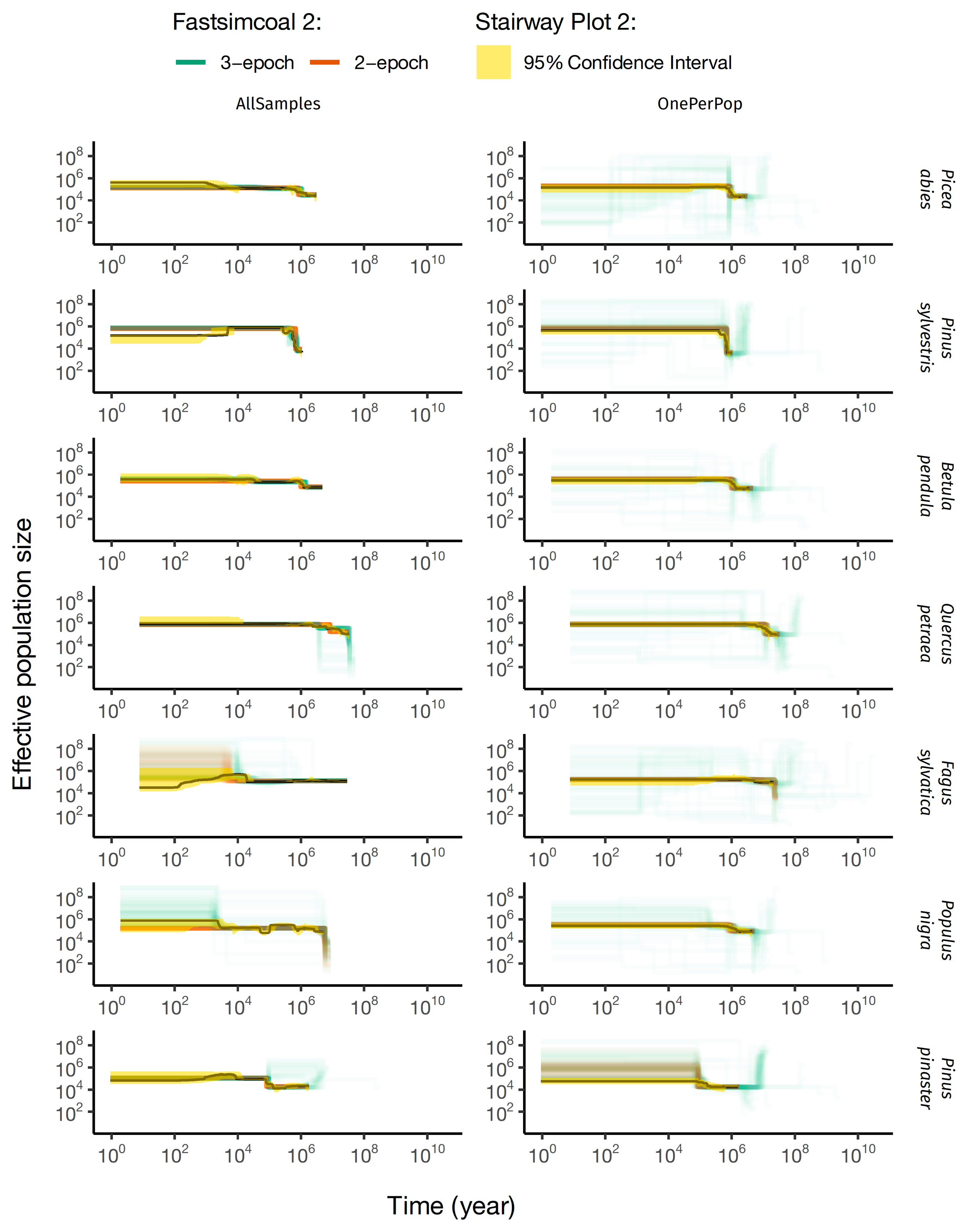

**Figure S26.** Comparison of the results obtained for the demographic dynamics of the seven forest tree species using two different sampling sets and three approaches. More specifically, results are presented for the Stairway Plot 2 model (in yellow) and for two demographic models tested using fastsimcoal2, i.e. 2-epoch (in red) and 3-epoch (in green), which allow one and two demographic events in the population, respectively. The analyses were performed using the site frequency spectrum computed either over all samples left panel) or over a random subset of one haplotype per location (OneperPop, right panel).

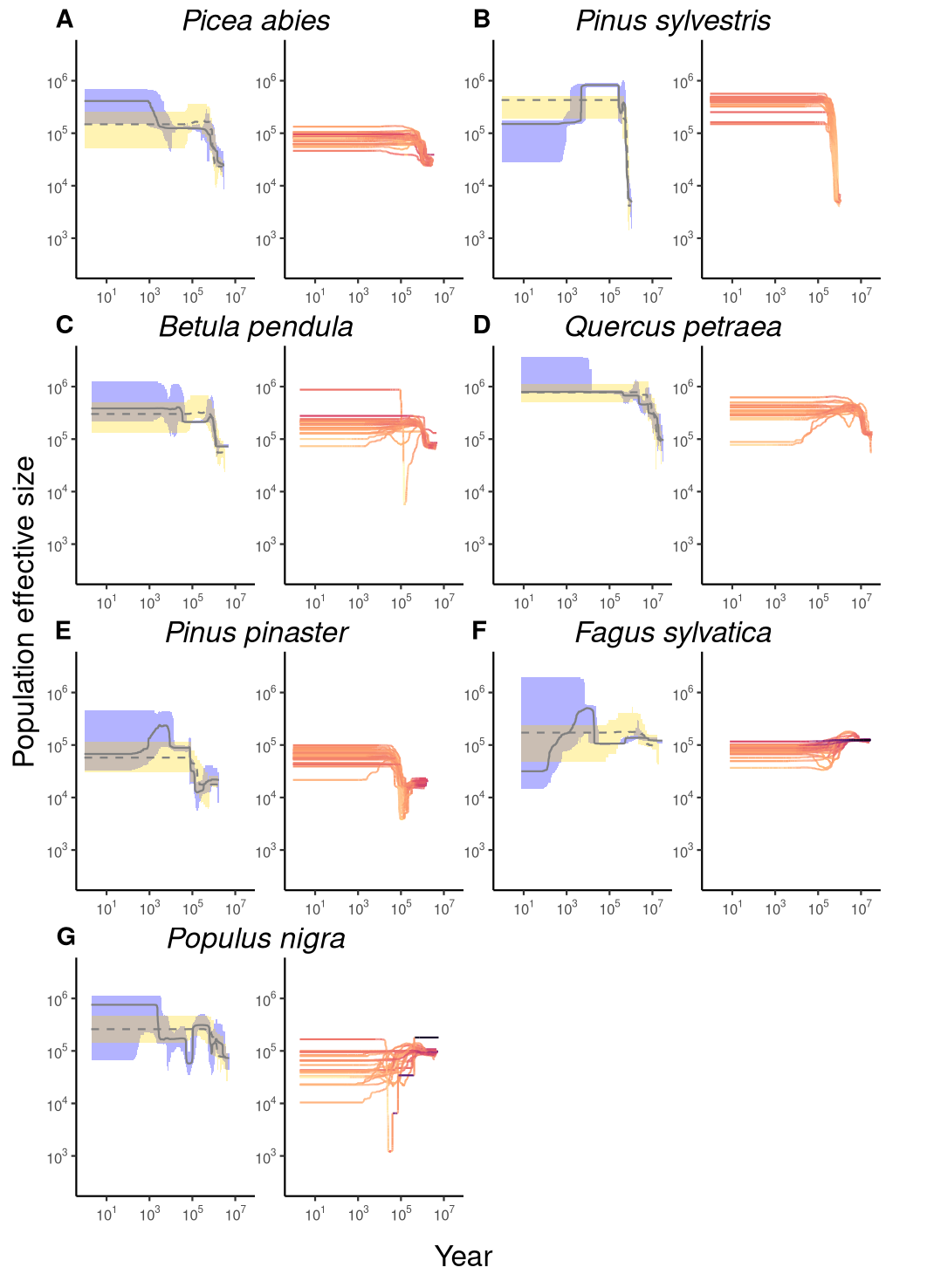

**Figure S27.** Stairway Plot 2 results for each species and across different sampling designs. For each species, results at the global and population levels are presented. At the global level (left plot in each panel), all-sample results (in blue) and one-per-population results (in yellow) are represented, with a line for the median level and a band for the 95% confidence interval. At the population level (right plot in each panel), each colored line represents the median result of the Stairway Plot 2 analysis run on a single population. Lighter to darker line colors represent areas with larger to smaller confidence intervals.

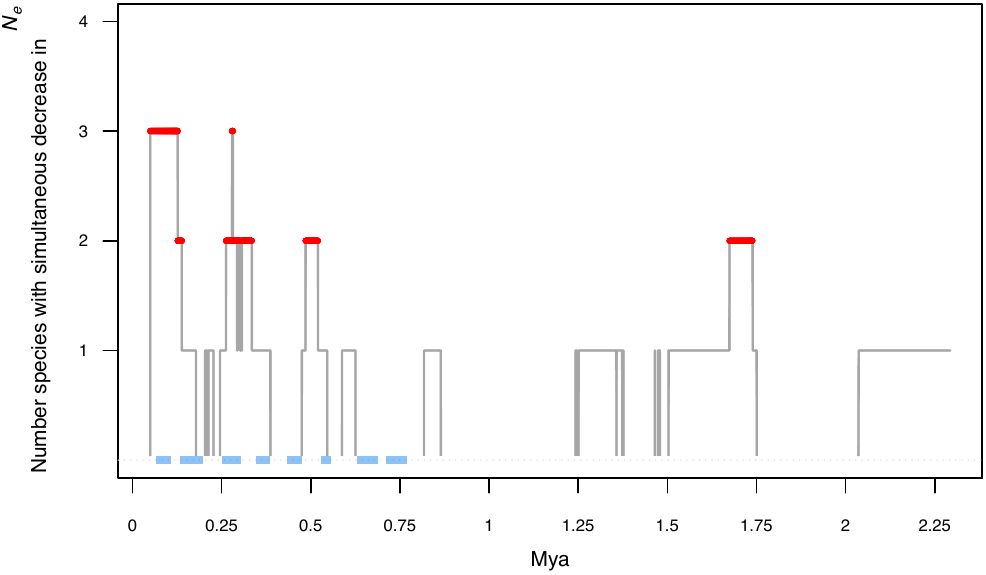

**Figure S28.** Synchronicity in decreasing phase in *N_e_* between the four species showing the highest correlations in *N_e_* dynamics (*Betula pendula*, *Picea abies*, *Pinus sylvestris* and *Populus nigra*). The solid line represents the number of species experiencing a decrease in *N_e_* at a given time point. The direction of change is given by the average change in *N_e_* across 250 time points using sliding windows see methods § “*Synchronicity analysis*”. Periods where the synchronicity in decreasing *N_e_* is larger than expected considering the actual change in *N_e_* over time, are highlighted in red. Blue areas along the x axis delineate glacial periods over the last 0.8 Mya. No significant synchronicity in decrease in *N_e_* was observed for *Q. petraea* and *F. sylvatica*.

Table S2. Number of populations and individuals per species. The complete list of sampled populations is given in Table S1. Populations not included in Opgenoorth et al. (2021) are listed as ‘additional populations’.

| **Species** | **Populations** | **Individuals** | **Additional populations** |
| --- | --- | --- | --- |
| *Betula pendula* (BP) | 23 | 497 | Belarus (BY_BP_24), Russia (RU_BP_25), Ukraine (UA_BP_23) |
| *Fagus sylvatica* (FS) | 26 | 602 | Austria (AT_FS_13, AT_FS_14), Slovenia (SI_FS_25, SI_FS_26, SI_FS_27, SI_FS_28) |
| *Populus nigra* (PO) | 22 | 467 | Great Britain (GB_PO_19), France (FR_PO_21), Morocco (MA_PO_23), Bosnia and Herzegovina (BA_PO_24) |
| *Picea abies* (PA) | 26 | 555 | Italy (IT_PA_03, IT_PA_04), Norway (NO_PA_22), Poland (PL_PA_25), Romania (RO_PA_24), Russia (RU_OB_01, RU_PA_19, RU_PA_20, RU_PA_26) |
| *Pinus pinaster* (PP) | 25 | 472 | France (FR_PP_21), Morocco (MA_PP_23, MA_PP_24), Portugal (PT_PP_22), Tunisia (TN_PP_25) |
| *Pinus sylvestris* (PS) | 23 | 411 | Russia (RU_PS_22, RU_PS_23) |
| *Quercus petraea* (QP) | 19 | 403 | - |

**Table S9** Synchronicity in decreasing phase in *N_e_* between the four species showing the highest correlations in *N_e_* dynamics (*Betula pendula*, *Picea abies*, *Pinus sylvestris* and *Populus nigra*). Span (number of consecutive $\Delta$*N_e_*) of synchronous decrease in *N_e_* for a given number of species are compared to the 95th percentiles of the distribution of the maximum spans obtained over 10,000 simulations where observed $\Delta$*N_e_* were directly randomized. Values are bolded and italicized when the duration of a given span is longer than that obtained through simulations, the signal for synchronicity in decreasing *N_e_* is hence considered as being significant. No significant synchronicity in decrease in *N_e_* was observed for *Q. petraea* and *F. sylvatica*.

| Two species | | | | |  | | Three species | | | | | |
| --- | --- | --- | --- | --- | --- | --- | --- | --- | --- | --- | --- | --- |
| Years (Kya) | | |  | Duration | |  | | Years (Kya) | | |  | Duration |
| Start |  | End |  |  |  |  | | Start |  | End |  |  |
| ***49.8*** |  | ***138.3*** |  | ***661*** | |  | | 49.8 |  | ***126.7*** |  | ***512*** |
| ***262.4*** |  | ***292.9*** |  | ***265*** | |  | | 278.9 |  | ***281.4*** |  | ***23*** |
| 294.4 |  | 301.8 |  | 79 | |  | | - |  | - |  | - |
| ***307*** |  | ***334.6*** |  | ***211*** | |  | | - |  | - |  | - |
| ***485.3*** |  | ***520*** |  | ***177*** | |  | | - |  | - |  | - |
| ***1674*** |  | ***1738.9*** |  | ***131*** | |  | | - |  | - |  | - |
| 95th percentile simulated span | | | | 103 | |  | | 95th percentile simulated span | | | | - |

Table S10. DNA extraction details for each species.

| **Species** | **DNA extraction method** | **Laboratory** |
| --- | --- | --- |
| *Betula pendula* | E.Z.N.A.® SP Plant DNA Kit (Omega Bio-tek) | University of Oulu, Finland |
| *Fagus sylvatica* | CTAB | Bavarian Office for Forest Seeding and Planting, ASP, Teisendorf, Germany |
| *Populus nigra* | Nucleospin Plant II Mini kit (Macherey-Nagel) | INRAE, BioForA, Orléans, France (except Greek samples extracted by Aristotle University of Tessaloniki, Greece) |
| *Picea abies* | DNeasy Plant Mini Kit (QIAGEN) | Uppsala University, Sweden |
| *Pinus pinaster* | DNeasy Plant Mini Kit (QIAGEN) | INRAE Biogeco, Bordeaux, France |
| *Pinus sylvestris* | DNeasy 96 Plant kit (QIAGEN) | UK Centre for Ecology and Hydrology |
| *Quercus petraea* | sbeadex maxi plant kit (LGC Genomics, Berlin, Germany) | Swiss Federal Institute for Forest, Snow and Landscape Research WSL, Switzerland |

Table S11. Probe design information and statistics.

| **Species** | **Reference for probe design** | **Best orthologs**  **(% bp)** | **Other orthologs**  **(% bp)** | **Candidates**  **(% bp)** | **Random**  **(% bp)** | **Criteria for additional targets** |
| --- | --- | --- | --- | --- | --- | --- |
| *Betula pendula* | *Betula pendula* subsp. *pendula* v.1.2 scaffolds, id35079 | 60.975 | - | 39.025 | - | Genes identified as putative targets of selective sweeps in (*14*) |
| *Fagus sylvatica* | *Fagus sylvatica* transcriptome assemblies (*67*, *68*); *F. sylvatica* candidate gene set (*69*) | 21.644 | 1.063 | 7.091 | 70.202 | Genes identified as differentially expressed in drought experiment (*68*) and during budburst (*67*); genes showing adaptive divergence on an elevation gradient (*70*) |
| *Populus nigra* | *Populus trichocarpa* v.3.0 for CDS  *Populus nigra* v.1.0 for full genes | 65.793 | 17.659 | 12.809 | 3.739 | Candidate genes based on annotation and differential expression across populations (*71*) |
| *Picea abies* | *Picea abies* v.1.0 | 58.365 | 0.378 | 4.336 | 36.921 | Orthologous genes between the three conifers;  genes identified as putative targets for positive selection in (*72*) |
| *Pinus pinaster* | Reference transcriptome at Gymno PLAZA v.1.0 | 42.276 | 6.802 | 10.063 | 40.859 | Orthologous genes between the three conifers; genes with potential roles in adaptation from (*30*, *73–77*); expressional candidate genes from (*78*) |
| *Pinus sylvestris* | Reference transcriptome at Gymno PLAZA v.1.0 | 48.420 | 8.846 | 6.744 | 35.990 | Orthologous genes between the three conifers;  genes with potential roles in adaptation from (*79*, *80*) |
| *Quercus petraea* | *Quercus robur* oak haplome v.2.3 (*72*) | 41.508 | - | 28.797 | 29.695  (14.219 intergenic sequence) | Candidate genes involved in local adaptation for response to water stress (*81*), and to temperature, precipitation and date of budburst (*82*);  intergenic sequences corresponding either to SNPs associated with local adaptation to temperature, precipitation and date of budburst or to random sequences. |

Table S15. Probe design information and statistics.

| **Species** | **CDS** | **UTR** | **Unclassified** | **Intergenic** |
| --- | --- | --- | --- | --- |
| *Betula pendula* | 1 | 0 | 0 | 0 |
| *Fagus sylvatica* | 1 | 0 | 0 | 0 |
| *Populus nigra* | 1 | 0 | 0 | 0 |
| *Picea abies* | 0.76 | 0.24 | 0 | 0 |
| *Pinus pinaster* | 0.99 | 0 | 0.01 | 0 |
| *Pinus sylvestris* | 0.96 | 0 | 0.04 | 0 |
| *Quercus petraea* | 0.63 | 0 | 0 | 0.37 |

Table S16. Reference genome used for mapping each species. mt; mitochondrial; cp: chloroplast

| **Species** | **Reference genome** | **Reference** |
| --- | --- | --- |
| *Betula pendula* | *Betula pendula* subsp. pendula v.1.2 scaffolds, id35079 (mt: LT855379.1, cp: LT855378.1) | (*14*) |
| *Fagus sylvatica* | *Fagus sylvatica* v.1.3 (cp: NCBI MK598696: mt (*Prunus avium*) NCBI NC_044768.1) | (*83*, *83*) |
| *Populus nigra* | *Populus trichocarpa* v.3.1 | (*84*) |
| *Picea abies* | *Picea abies* v.1.0 | (*85*, *85*) |
| *Pinus pinaster* | *Pinus taeda* v.2.01 (mt: NC_039746.1, cp: NC_0214401.1) | (*86*, *87*) |
| *Pinus sylvestris* | *Pinus taeda* v.2.01 (mt: NC_039746.1, cp: NC_0214401.1) | (*86*, *87*) |
| *Quercus petraea* | *Quercus robur* v.2.3 | (*81*) |

Table S17. Sequencing and mapping statistics. For each species we report the total number of bp sequenced, the number of Illumina reads mapping uniquely in the genome, and the portion of the reference genome in which at least 50% of the samples of the given species have a minimum coverage of 8x and a minimum genotype quality value of 20.

| **Species** | **Gbp sequenced** | **Uniquely mapping reads** | **Available genome** |
| --- | --- | --- | --- |
| *Betula pendula* | 395.41 | 2,802,450,105 | 6,041,548 |
| *Fagus sylvatica* | 644.50 | 2,673,500,437 | 6,569,688 |
| *Populus nigra* | 507.92 | 2,435,020,528 | 6,106,889 |
| *Picea abies* | 706.78 | 1,517,348,029^a^ | 4,974,709 |
| *Pinus pinaster* | 1146.77 | 2,156,038,221 | 3,068,914 |
| *Pinus sylvestris* | 538.90 | 1,401,611,778 | 1,407,443 |
| *Quercus petraea* | 531.30 | 2,558,016,107 | 6,323,296 |

^a^ Computed after duplicated reads were removed.

**Table S18: Proportion of SNPs in the various categories (SNP set v.5.3.1).**

| **Species** | **Best-orthologs** | **Other-orthologs** | **Candidates** | **Random** | **Intergenic** |
| --- | --- | --- | --- | --- | --- |
| *Betula pendula* | 0.66 | 0.00 | 0.26 | - | 0.08 |
| *Fagus sylvatica* | 0.16 | 0.01 | 0.04 | 0.39 | 0.40 |
| *Picea abies* | 0.53 | 0.00 | 0.05 | 0.36 | 0.06 |
| *Populus nigra* | 0.70 | 0.11 | 0.08 | 0.02 | 0.09 |
| *Pinus pinaster* | 0.17 | 0.03 | 0.04 | 0.17 | 0.58 |
| *Pinus sylvestris* | 0.13 | 0.02 | 0.03 | 0.11 | 0.71 |
| *Quercus petraea* | 0.38 | 0.00 | 0.12 | 0.14 | 0.36 |

Table S19: Proportion of SNPs in different structural classes (SNP set v.5.3.1).

| **Species** | **0-fold** | **2-3 -fold** | **4-fold** | **intergenic** | **intron** | **top** | **up** | **down** |
| --- | --- | --- | --- | --- | --- | --- | --- | --- |
| *Betula pendula* | 0.15 | 0.08 | 0.10 | 0.17 | 0.50 | 0.00 | 0.00 | 0.00 |
| *Fagus sylvatica* | 0.16 | 0.08 | 0.10 | 0.21 | 0.44 | 0.00 | 0.00 | 0.00 |
| *Populus nigra* | 0.17 | 0.10 | 0.11 | 0.08 | 0.43 | 0.00 | 0.04 | 0.07 |
| *Picea abies* | 0.20 | 0.10 | 0.09 | 0.41 | 0.19 | 0.00 | 0.00 | 0.00 |
| *Pinus pinaster* | 0.16 | 0.08 | 0.09 | 0.52 | 0.16 | 0.00 | 0.00 | 0.00 |
| *Pinus sylvestris* | 0.14 | 0.06 | 0.05 | 0.70 | 0.05 | 0.00 | 0.00 | 0.00 |
| *Quercus petraea* | 0.10 | 0.06 | 0.06 | 0.39 | 0.35 | 0.00 | 0.02 | 0.02 |

Table S20: Number of SNPs is each SNP set and species.

| **Species** | **v5.3** | **v5.3.1** | **v5.3.2** | **v6.3.1** | **v6.3.2** |
| --- | --- | --- | --- | --- | --- |
| *Betula pendula* | 213250 | 213250 | 213250 | 88741 | 64697 |
| *Fagus sylvatica* | 223185 | 166939 | 166939 | 54039 | 40506 |
| *Populus nigra* | 197204 | 197204 | 197204 | 41841 | 33857 |
| *Picea abies* | 290166 | 290166 | 290166 | 135562 | 98624 |
| *Pinus pinaster* | 111937 | 111937 | 111937 | 50108 | 33418 |
| *Pinus sylvestris* | 89903 | 89903 | 89903 | 65205 | 40650 |
| *Quercus petraea* | 507026 | 479619 | 479619 | 174110 | 134010 |
